# AI-Accelerated Structure Elucidation of Boavistamides A–C, Cyclic Depsipeptides from a Marine Filamentous Cyanobacterium Collected in Cabo Verde

**DOI:** 10.64898/2026.06.13.732064

**Authors:** Marine Cuau, Nicole E. Avalon, Byeol Ryu, Evgenia Glukhov, Jehad Almaliti, Adriana Rego, Thaiz R. Teixeira, Mai Shingyoji, Mariana L. De Souza, Alma Trinidad-Javier, Krittikorn Kumpornsin, Joseph Chen, Case W. McNamara, Conor R. Caffrey, Elizabeth A. Winzeler, Vitor M. Vasconcelos, Pedro N. Leão, William H. Gerwick

**Affiliations:** Center for Marine Biotechnology and Biomedicine, Scripps Institution of Oceanography, University of California San Diego, La Jolla, California 92093, United States; Interdisciplinary Centre of Marine and Environmental Research (CIIMAR), University of Porto, Matosinhos 4450-208, Portugal; Department of Pharmaceutical Sciences, School of Pharmacy & Pharmaceutical Sciences, University of California, Irvine, 856 Health Sciences Road, Suite 5400, Irvine, California 92697-3958, United States; Robert A. Mah Molecular Innovation Center, University of California, Irvine, Irvine, California 92697, United States; Skaggs School of Pharmacy and Pharmaceutical Sciences, University of California San Diego, La Jolla, California 92093, United States; Center for Discovery and Innovation in Parasitic Diseases, Skaggs School of Pharmacy and Pharmaceutical Sciences, University of California San Diego, La Jolla, California 92093, United States; Department of Pediatrics, School of Medicine, University of California, San Diego, La Jolla, California 92093, United States; Calibr-Skaggs Institute for Innovative Medicines, A Division of Scripps Research, La Jolla, California 92093, United States

## Abstract

Boavistamide A (**1**), a new alkyne-containing cyclic depsipeptide featuring the rare 3-amino-2-methyl-7-octynoic acid (AMOYA) moiety, was discovered along with two structurally related analogs, boavistamides B and C (**2** and **3**), from a filamentous marine cyanobacterium collected on Boa Vista Island, Cabo Verde. Their isolation was guided by antiplasmodial activity, GNPS MS/MS molecular networking, LC-MS profiling, and dereplication using the MarinLit database. The planar structures of boavistamides A–C (**1**–**3**) were elucidated through comprehensive HRMS and 1D/2D NMR analyses, with annotation support from AI-based tools SMART-NMR 2.1 and DeepSAT. The absolute configurations were established using Marfey’s analysis and L-Phe-OMe coupling, complemented by NMR-based conformational studies. Boavistamides A and B exhibited moderate antiplasmodial activity with no mammalian cell cytotoxicity. Microscopic observations and metagenomic binning identified the producer strain as belonging to the genus *Okeania* (Microcoleaceae). These results expand the chemical diversity of AMOYA-containing cyanobacterial metabolites and highlight the utility of integrated metabolomics and AI-assisted workflows for natural product discovery from environmental samples.

**GRAPHICAL ABSTRACT:** 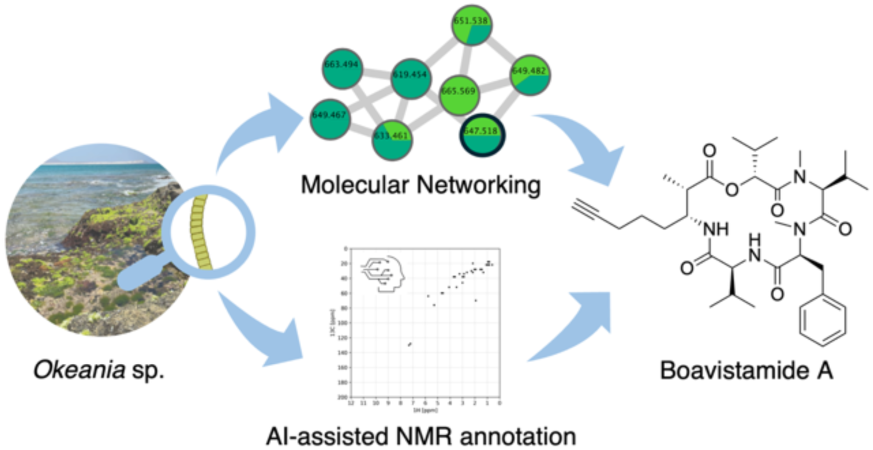

---

Marine cyanobacteria are prolific producers of chemically diverse natural products (NPs), many of which display bioactive properties and pharmaceutical potential. Among them, filamentous forms such as those from the genus *Okeania* have yielded a broad spectrum of bioactive metabolites.^1,2,3^ This chemodiversity is largely attributed to their rich biosynthetic machinery, particularly hybrid nonribosomal peptide synthetase (NRPS) and polyketide synthase (PKS) pathways, coupled with ecological adaptation across marine environments.^4,5,6^ Cyclic depsipeptides represent a prominent class of cyanobacterial metabolites, often featuring modified residues such as *β*-hydroxy acids, *N*-methyl amino acids, and unusual lipophilic moieties. Rare among these are metabolites with a 3-amino-2-methyl-7-octynoic acid (AMOYA) residue, which has been reported in only a few NPs to date.^7,8,9,10,11,12^ Advances in artificial intelligence (AI) are increasingly transforming NP structure elucidation. Accessible tools such as SMART-NMR and DeepSAT provide rapid, database-assisted annotation of experimental NMR spectra, drastically narrowing structural possibilities within seconds.^13,14^ Although these tools do not replace expert manual NMR interpretation, they have rapidly become an integral component of contemporary NP workflows; accelerating dereplication and improving the efficiency of planar structure assignment.^13,15,16,17,18,19^

In this study, a marine cyanobacterial sample collected in an underexplored region of Insular West Africa (Ilhéu de Sal Rei, Boa Vista Island, Cabo Verde) was initially profiled by LC–MS/MS. The crude extract was fractionated and the resulting fractions were analyzed using the Global Natural Products Social (GNPS) molecular networking tool.^20^ This molecular networking analysis revealed a distinct cluster of related metabolites with an intriguing molecular composition, which, together with antiparasitic activity observed in selected fractions, prompted its prioritization for further investigation. Subsequently, we isolated boavistamide A (**1**), an AMOYA-containing cyclic depsipeptide, along with two structurally related analogs, boavistamides B and C (**2** and **3**). The predominant filamentous cyanobacterium present in the sample was taxonomically identified as *Okeania* sp. based on microscopic and metagenomic evidence. Investigating marine cyanobacteria from underexplored regions such as Insular West Africa remains a valuable strategy for NP discovery, as habitat, ecological and taxonomic biodiversity often correlate with chemodiversity and novelty.^21^ The remote Cabo Verde archipelago harbors a rich and largely uncharacterized cyanobacterial diversity.^22^ Nevertheless, to the best of our knowledge, no NPs have previously been reported from marine organisms collected on Boa Vista Island, Cabo Verde.

Structural elucidation of the boavistamides was supported by high-resolution MS, 1D/2D NMR, and significantly accelerated by use of AI-based NMR annotation tools (SMART-NMR and DeepSAT). The absolute configuration of the amino acid and hydroxy acid residues was determined using chiral derivatization and comparison with authentic standards, while the configuration of the AMOYA unit was assigned based on NMR coupling constants and ROESY correlations. The biological activity of the boavistamides was evaluated in several assays and revealed moderate antiplasmodial activity for compounds **1** and **2** with no cytotoxicity in mammalian cell lines (HEK293T, HepG2, and SF188). Collectively, this study highlights how computational metabolomics, metagenomics, and AI tools can accelerate NP discovery and structure elucidation, and underscores the underexplored potential of Cabo Verdean marine filamentous cyanobacteria as reservoirs of new metabolites.

## RESULTS AND DISCUSSION

### A) Cyanobacterial Collection and Taxonomic Assignment

A dark green, filamentous biofilm was collected from a rocky intertidal platform on Ilhéu de Sal Rei, Boa Vista Island, Cabo Verde (Table S1, Figures S1–S2). Light microscopy revealed abundant long filaments of cyanobacteria, consistent with the family Microcoleaceae. Short-read shotgun metagenomic sequencing (DNBSEQ™) and genome-resolved analysis further supported this taxonomic assignment. The metagenome assembly and binning yielded 204 microbial bins, of which ten were taxonomically assigned to Cyanobacteria using GTDB-Tk (Figure S3). Quality assessment with CheckM indicated that four cyanobacterial bins met the criteria for high-quality (completeness > 90%, contamination < 5%) metagenome-assembled genomes (MAGs). Among these, the cyanobacterial MAG of highest quality (Bin 81; 94.6% completeness, 3.9% contamination) taxonomically assigned to the genus *Okeania*, was also the dominant cyanobacterial lineage according to Anvi’o abundance profiling (Figure S4, Table S2).

### B) Extraction and Bioassay Guided Isolation

The freeze-dried cyanobacterial biomass was extracted using a 2:1 mixture of CH_2_Cl_2_/MeOH. The resulting organic extract was subjected to vacuum liquid chromatography (VLC) on silica gel, yielding 12 fractions (A–K) based on increasing polarity (normal phase). Bioactivity screening of all fractions was performed against *Plasmodium falciparum, Trypanosoma brucei,* and a panel of microbial strains, including *Bacillus subtilis, Escherichia coli, Staphylococcus aureus, Salmonella typhimurium,* and *Candida albicans*. Fractions D (eluted with 40% EtOAc in hexanes) and E (eluted with 60% EtOAc in hexanes) exhibited selective antiparasitic activity. Fraction D was active against *P. falciparum*, while fraction E displayed dual activity against *P. falciparum* and *T. brucei.* Neither fraction showed significant antibacterial or antifungal activity, and only weak cytotoxicity was observed against H460 human lung carcinoma cells (Table S3, Figures S5–S8). Guided by the antiplasmodial activity and LC-MS data, fractions D and E were prioritized for purification. LC-MS analysis revealed three dominant peaks at *m/z* 625.36, 627.47, and 629.33 [M + H]^+^, corresponding to the molecular mass of compounds later named boavistamides A–C (**1–3)** (Figure 1A and Figures S9–S11). These fractions were subjected to reversed-phase HPLC, resulting in the isolation of boavistamides A–C (**1–3)** (Figure 1B).

**Figure 1.**
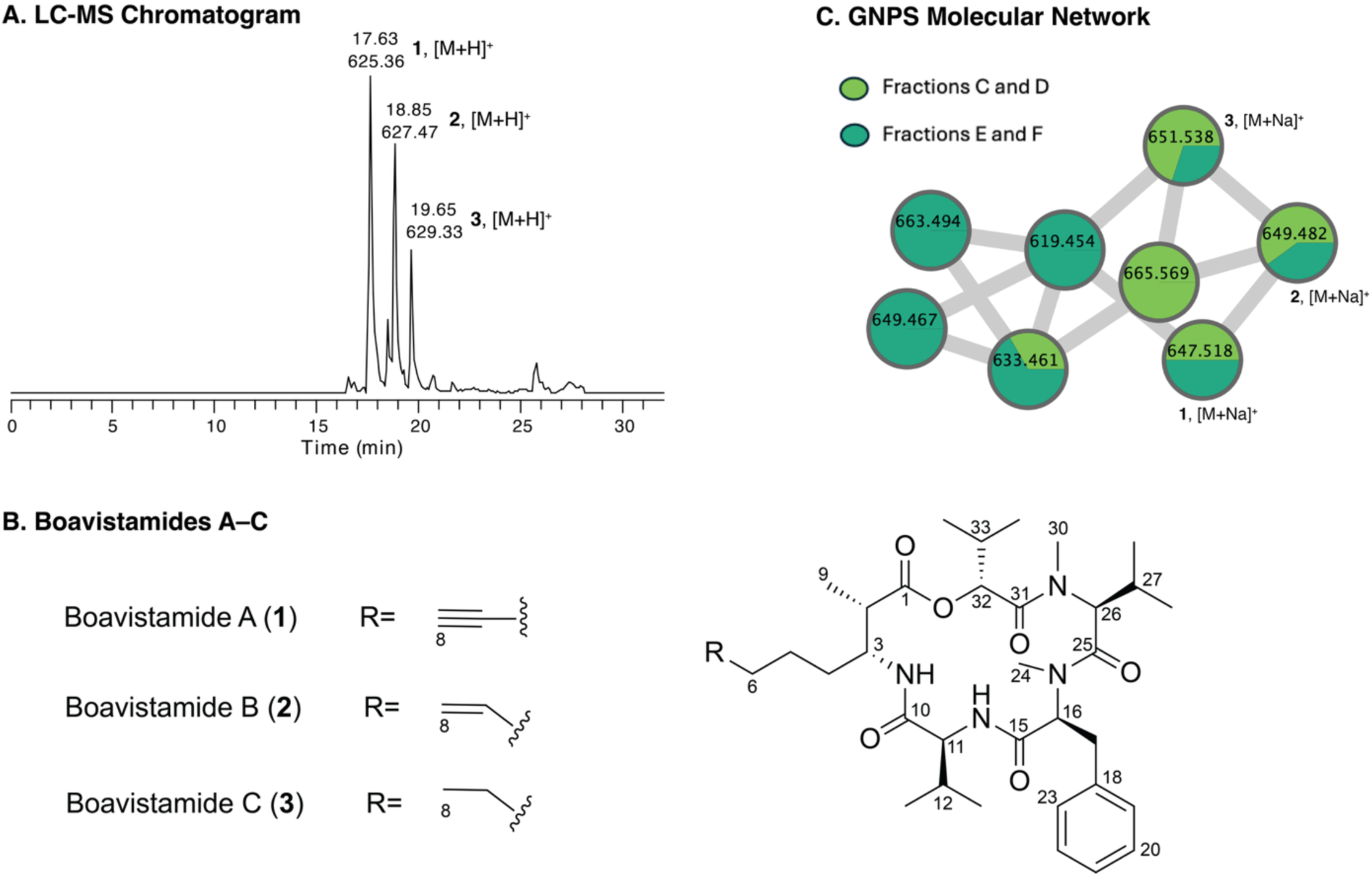
(A) Base peak chromatogram (retention time 0–32 min) of fraction D, showing boavistamides A–C (**1**–**3**) as the major components detected as protonated molecules at *m/z* 625 [M + H]^+^, 627 [M + H]^+^, and 629 [M + H]^+^, with the peak at *m/z* 625 corresponding to the 100% relative abundance peak. (B) Chemical structures of boavistamides A–C (**1**–**3**). (C) Selected portion of the classical molecular network generated from LC-MS/MS data of VLC fractions. Compounds (**1**–**3**) form a distinct cluster associated with fractions C–F. Note that the nodes corresponding to boavistamides A–C (**1**–**3**) are labeled in the molecular network with their respective precursor ion masses detected as sodium adducts at *m/z* 647 [M + Na]^+^, 649 [M + Na]^+^, and 651 [M + Na]^+^ rather than the [M + H]^+^ adducts as seen in Panel A. The full molecular network is shown in Figure S12.

### C) GNPS Molecular Networking

To assist in the dereplication and prioritization of secondary metabolites, MS/MS molecular networking was performed using the GNPS platform.^20^ A classical molecular network was generated from the LC-MS/MS data acquired from extract-derived VLC fractions, which was visualized using Cytoscape (Figures 1C and S12).^23^ This analysis revealed a distinct molecular family that did not match any reference spectra in the GNPS spectral libraries. Within this cluster, boavistamides A–C were identified by their precursor ion masses at *m/z* 647.5 [M + Na]^+^, 649.5 [M + Na]^+^, and 651.5 [M + Na]^+^, respectively. Preliminary dereplication using the NP database MarinLit (https://marinlit.rsc.org/) did not yield any matches to known compounds, suggesting that the metabolites were potentially novel and motivating further isolation and structure elucidation. Subsequent LC-MS analysis confirmed that fractions D and E contained the target metabolites in sufficient quantities for comprehensive structure elucidation, showing three major peaks at the corresponding *m/z* values of 625 [M + H]^+^, 627 [M + H]^+^, and 629 [M + H]^+^ (Figure 1A). These fractions were subsequently purified by HPLC, yielding boavistamides A–C (**1**–**3**) (Figure 1B). Together, the combination of molecular networking, LC-MS analysis, and database-assisted dereplication enabled efficient prioritization and targeted isolation of the boavistamides.

### D) Structure Elucidation

Boavistamide A (**1**), an amorphous white solid, was assigned the molecular formula C₃₅H₅₂N₄O₆ based on high-resolution electrospray ionization mass spectrometry (HRESIMS) (Figure S13). Initial insights into the structure of compound **1** were obtained from 1D and 2D NMR analyses, including ^1^H, ^13^C, COSY, HSQC, HMBC, and ROESY experiments (Tables 1 and S4, Figures S14–19). The ^1^H NMR spectrum revealed the presence of seven methyl groups (δ_H_ 1.21, 1.06, 0.95 0.91, 0.81, 0.79, 0.55), two amide protons (δ_H_ 5.85, 6.71), two sharp singlets consistent with N-methyl groups (δ_H_ 2.85, 3.21), and one terminal alkyne proton (δ_H_ 1.88). The ^13^C NMR and HSQC spectra confirmed the presence of five carbonyl signals (δ_C_ 173.5, 172.5, 171.8, 170.5, 169.6), a terminal alkyne carbon (δ_C_ 68.8), and an aromatic system (δ_C_ 137.9, 129.5, 128.9, 127.0).

To accelerate structure elucidation and refine structural hypotheses, we employed two AI-based annotation tools: SMART-NMR 2.1 (http://smart.ucsd.edu/classic) and DeepSAT (https://deepsat.ucsd.edu).^13,14^ SMART-NMR uses a convolutional neural network (CNN) trained on a large database of HSQC spectra to generate ranked structural predictions. SMART analysis of the standard ^1^H–^13^C HSQC spectrum of compound **1** returned several cyclic depsipeptides featuring terminal alkyne-containing fatty acyl chains and aromatic moieties among the top 10 hits (Figure 2A). The best matches included ulongapeptin, veraguamide F, and dudawalamide C, all with a high cosine similarity score of 0.94.^7,25,29^ In parallel, DeepSAT, which uses a multitask CNN to predict Morgan-type fingerprints from HSQC data, produced complementary results. Normal and multiplicity-edited ^1^H−^13^C HSQC data obtained for compound **1** were applied to DeepSAT; both spectral inputs showed the same top 10 matches with 100% oligopeptides containing a terminal alkyne chain (Figure 2B). Antanapeptin D emerged as the closest match with a high cosine similarity score of 0.91.^24^ Both SMART NMR and DeepSAT converged on closely related structural frameworks, reinforcing the initial observation that compound **1** was a cyclic alkyne-bearing depsipeptide. Additionally, these predictions, in combination with classical NMR data, supported the presence of the rare 3-amino-2-methyl-7-octynoic acid (AMOYA) unit as the lipophilic residue, previously identified in only a handful of cyanobacterial metabolites, including ulongapeptin.

**Figure 2.**
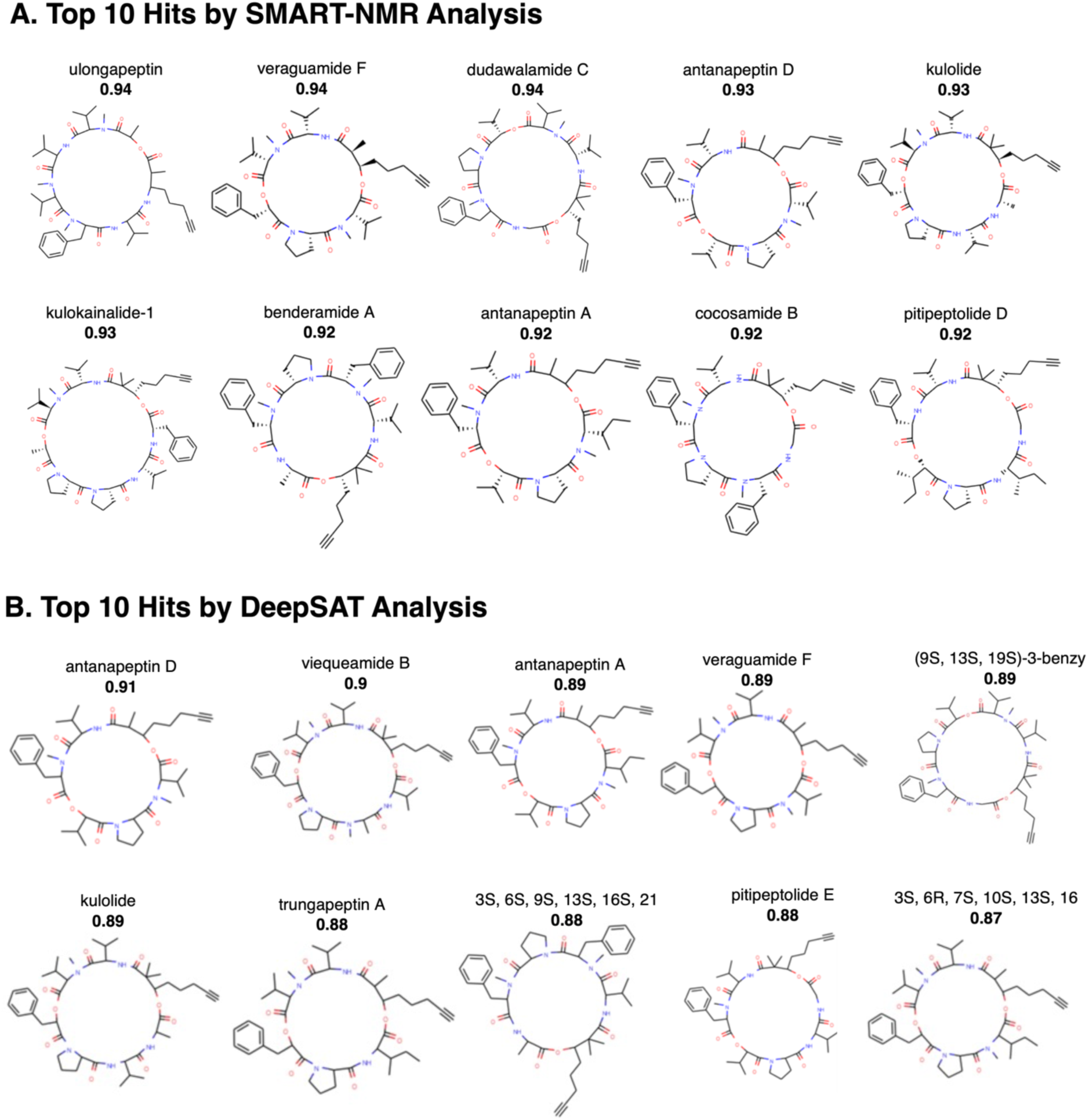
Top 10 hits by SMART-NMR 2.1 analysis (A) and DeepSAT analysis (B) of boavistamide A (**1**) in CDCl_3_. When DeepSAT was used, normal HSQC and multiplicity-edited HSQC provided equivalent results.

Further interpretation of the 2D NMR spectra confirmed the macrocyclic depsipeptide structure of boavistamide A (**1**). COSY, HMBC and ROESY correlations (Table S4) allowed for the sequential construction of amino acid and hydroxy acid units, including the sequence AMOYA–hydroxyisovaleric acid (HIVA)–*N*-Me-Val–*N*-Me-Phe–Val. COSY correlations between H-9/H-2/H-3/3-NH/H-4/H-5/H-6, together with HMBC correlations H-5/C-7 and H-6/C-8, established the AMOYA residue. COSY correlations between H-32/H-33/H-34/H-35, combined with HMBC correlations H-32/C-1 and H-2/C-1, connected the HIVA moiety to the AMOYA residue through an ester linkage. The COSY correlations between H-26/H-27/H-28/H-29 defined the *N*-Me-Val residue. HMBC correlations H-30/C-31 and H-30/C-26 placed the *N*-methyl group and established the connection between *N*-Me-valine and HIVA. A ROESY correlation between H-30/H-32 further supported this linkage. COSY correlations between H-16/H-17, together with HMBC correlations H-19/C-21, H-17/C-19, and H-24/C-16, defined the *N*-Me-phenylalanine residue. Its attachment to *N*-Me-Val was supported by HMBC correlations H-24/C-25 and H-26/C-25, as well as a ROESY correlation between H-26/H-16. HMBC correlations from the two sharp methyl singlets (δ_H_ 2.85, 3.21) to amide carbonyl carbons, together with the molecular formula and comparison with structurally related depsipeptides such as ulongapeptin, supported the assignment of two tertiary *N*-methylamide groups. COSY correlation between H-11/H-12/H-13/H-14, along with HMBC correlations 11-NH/C-11 and H-11/C-10, established the valine residue. HMBC correlations 11-NH/C-15 connected the valine to *N*-Me-phenylalanine, while an HMBC correlation 3-NH/C-10 linked the valine to the AMOYA residue, completing the macrocyclic framework. Taken together, these data allowed elucidation of the complete planar structure of boavistamide A (**1**), revealing a novel cyclic depsipeptide scaffold featuring the rare AMOYA lipid moiety, three amino acid residues, two of which were *N*-methylated, and one hydroxy acid (Figure 3).

**Figure 3.**
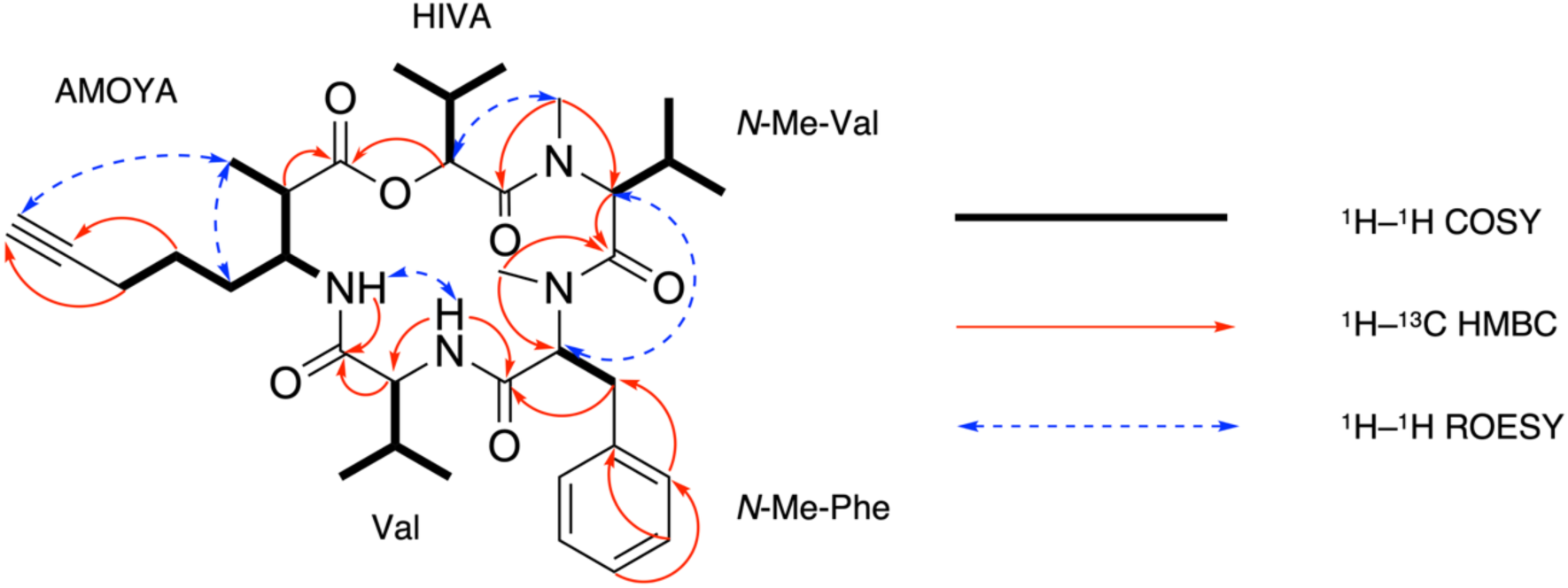
Key 2D NMR correlations of boavistamide A (**1**)

The planar structures of boavistamides B (**2**) and C (**3**) were elucidated using the same strategy as for boavistamide A (**1**), combining HRMS analysis (Figures S20 and S26) with comprehensive 1D and 2D NMR experiments (HSQC, COSY, HMBC, and ROESY) (Tables 1, S5 and S6, Figures S21–S25 and S27–S31). A comparative analysis of the ^1^H NMR spectra of compounds (**1–3**) revealed a conserved cyclic depsipeptide scaffold with a key variation localized within the AMOYA residue. Specifically, boavistamide B (**2**) displays signals consistent with a terminal alkene in place of the terminal alkyne found in (**1**), while boavistamide C (**3**) features a fully reduced alkyl chain. These progressive reductions are evident in the chemical shift changes of the olefinic and aliphatic proton regions (Figure S32). Similar trends have been reported in other cyanobacterial cyclic depsipeptides including the antanapetins, veraguamides and kohamamides.^24,25,26^

### D) Absolute Configuration of Boavistamides A–C (1–3)

To establish the absolute configuration of boavistamide A (**1**), a combination of advanced Marfey’s analysis, L-Phe-OMe derivatization, and NMR experiments was employed (Figure 4). Following acid hydrolysis, Marfey’s derivatization and LC-MS analysis against authentic standards identified the amino acid residues as *N*-Me-L-Val, *N*-Me-L-Phe, and L-Val. The hydroxy acid residue, determined as HIVA, was derivatized using L-phenylalanine methyl ester (L-Phe-OMe), and comparison with authentic D- and L-HIVA derivatives confirmed the (*R)*-HIVA configuration (Figures S33–S35). In contrast, the configuration of the 3-amino-2-methyl-7-octynoic acid (AMOYA) residue in boavistamide A (**1**) was investigated using NMR analysis. The H-3 resonance is consistent with a dddd spin system arising from coupling to the NH proton, vicinal H-2, and the diastereotopic H-4a/H-4b protons. A vicinal H-2/H-3 coupling constant of approximately 7 Hz indicates an anti relationship between H-2 and H-3. This arrangement can be accommodated by both *erythro* and *threo* configurations in idealized Newman projections. ROESY correlations between H-3/CH_3_–9 and H-2/H-4 are likewise consistent with both possibilities in such representations (Figures S36–S37). However, considering the conformational constraints imposed by the macrocyclic scaffold, which limits conformational flexibility, ROESY correlations become diagnostically meaningful. Under these conditions, only the *erythro* configuration is consistent with the observed spatial proximities, supporting an *erythro* relationship at C-2/C-3. Inter-residue ROESY correlations further place the AMOYA unit in defined spatial proximity to adjacent residues of known configuration (Figures S38–S40). To determine the absolute configuration, two full macrocyclic 3D models were generated in which the configurations of the adjacent residues, (*R*)-HIVA and L-Val, were fixed, while the AMOYA C-2/C-3 configuration was set as either (2*S*,3*R*) or (2*R*,3*S*). Only the (2*S*,3*R*) model satisfied the observed intra- and inter-residue ROESY correlations, whereas the (2*R*,3*S*) model failed to reproduce these spatial proximities with the adjacent residues (Figure S40–S41). Further evaluation of all possible C-2/C-3 stereochemical configurations confirmed that only the (2*S*,3*R*) model satisfies the observed ROESY-derived spatial constraints, supporting assignment of the AMOYA residue as (2*S*,3*R*). This assignment is also consistent with related AMOYA-containing peptides such as companeramide A (H-2/H-3 *J* = 7.2 Hz), which displays a similar large vicinal coupling, and contrasts with *threo*-configured analogues including portobelamide B (H-2/H-3 *J* = 2.4 Hz) and ulongapeptin (H-2/H-3 *J* = 3.6 Hz), where significantly smaller coupling constants are observed (Table S7).^11,8,7^ The same analytical strategy was applied to analogs **2** and **3**. The configurations of the amino acid and hydroxy acid residues were confirmed by chemical derivatization, and comparative NMR analysis indicated that the corresponding C-2 and C-3 stereocenters are the same as those of the AMOYA unit in boavistamide A (Figures S42–S46).

**Figure 4.**
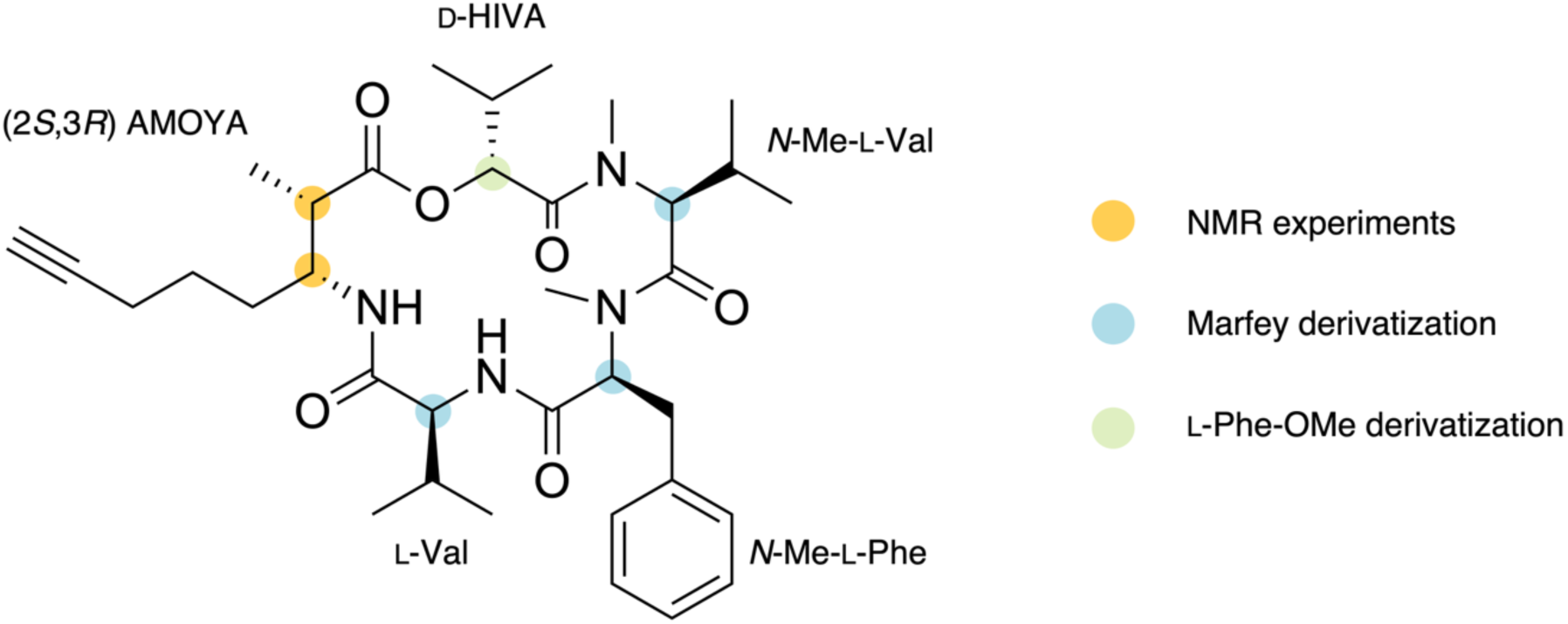
Absolute configuration of boavistamide A (**1**)

### E) Comparative Analysis of AMOYA-containing Depsipeptides

SMART-NMR and DeepSAT results for boavistamide A (**1**) provided matches to cyclic depsipeptides containing terminal alkyne functions, including several belonging to the kulolide superfamily.^27^ The kulolide superfamily is defined by the presence of a β-hydroxyoctynoic acid derivative, traditionally subdivided into two structural subgroups based on the lipid tail: DHOYA (2,2-dimethyl-3-hydroxy-7-octynoic acid) and HMOYA (3-hydroxy-2-methyl-7-octynoic acid).^28,29,30^ These lipophilic units with a terminal alkyne are embedded within a macrocyclic scaffold composed of amino and hydroxy acid residues. Members of this superfamily are characteristically produced by filamentous marine cyanobacteria, exhibit antiparasitic or cancer cell cytotoxic activities, and based on their structural features, are presumed to be assembled via hybrid NRPS–PKS biosynthetic pathways. ^27^

The AMOYA (3-amino-2-methyl-7-octynoic acid) residue, originally identified as a component of the mollusk-derived peptide onchidin in 1994, has since been reported in eight additional cyclic depsipeptides from marine cyanobacteria: malevamide C, portobelamides A and B, companeramides A and B, ulongapeptin, guineamide C, along with the newly discovered boavistamide A (**1**) (Figure 5). ^31,7,8,9,10,11^ These AMOYA-containing peptides exhibit strong parallels with the kulolide superfamily: a cyanobacterial origin, the presence of a terminal alkyne-containing lipid moiety structurally analogous to DHOYA or HMOYA, comparable biological activities (notably cytotoxic and/or antiparasitic), and biosynthetic logic consistent with a PKS–NRPS hybrid assembly. Notably, in all reported AMOYA-containing cyclic depsipeptides except onchidin, the AMOYA unit is directly linked to a hydroxy acid residue, suggesting a conserved structural motif that may reflect shared biosynthetic organization. However, the stereochemistry of these hydroxy acid residues is not uniformly conserved across this metabolite class. While boavistamides incorporate (*R*)-HIVA, related metabolites such as malevamide C and companeramides contain the opposite enantiomer, (*S*)-HIVA.^10,8^ This indicates that, despite architectural conservation, stereochemical control may vary between pathways, potentially involving ketoreductase-mediated steps, although any enzymatic basis remains speculative in the absence of direct genetic evidence. To date, no biosynthetic gene cluster (BGC) has been identified for any DHOYA-, HMOYA-, nor AMOYA-containing cyclic depsipeptide, preventing a definitive phylogenetic classification and reinforcing the value of a structure-based comparative approach. AntiSMASH (v7.0) analysis of the entire metagenomics dataset from the Cabo Verde cyanobacterial sample did not identify a contiguous biosynthetic gene cluster that could be clearly associated with boavistamide production, possibly due to fragmentation of the metagenome assembly (Figure S52). Previous kulolide superfamily members and AMOYA-containing cyclic peptides have predominantly been reported from Microcoleaceae, including genera such as *Okeania* and *Moorea* (formerly classified as *Lyngbya).*^7,9,29,32^ *T*ogether with the morphological observation, taxonomic assignment, and relative abundance evidence, this supports *Okeania* sp. as the putative producer of boavistamides A–C (**1–3**).

**Figure 5.**
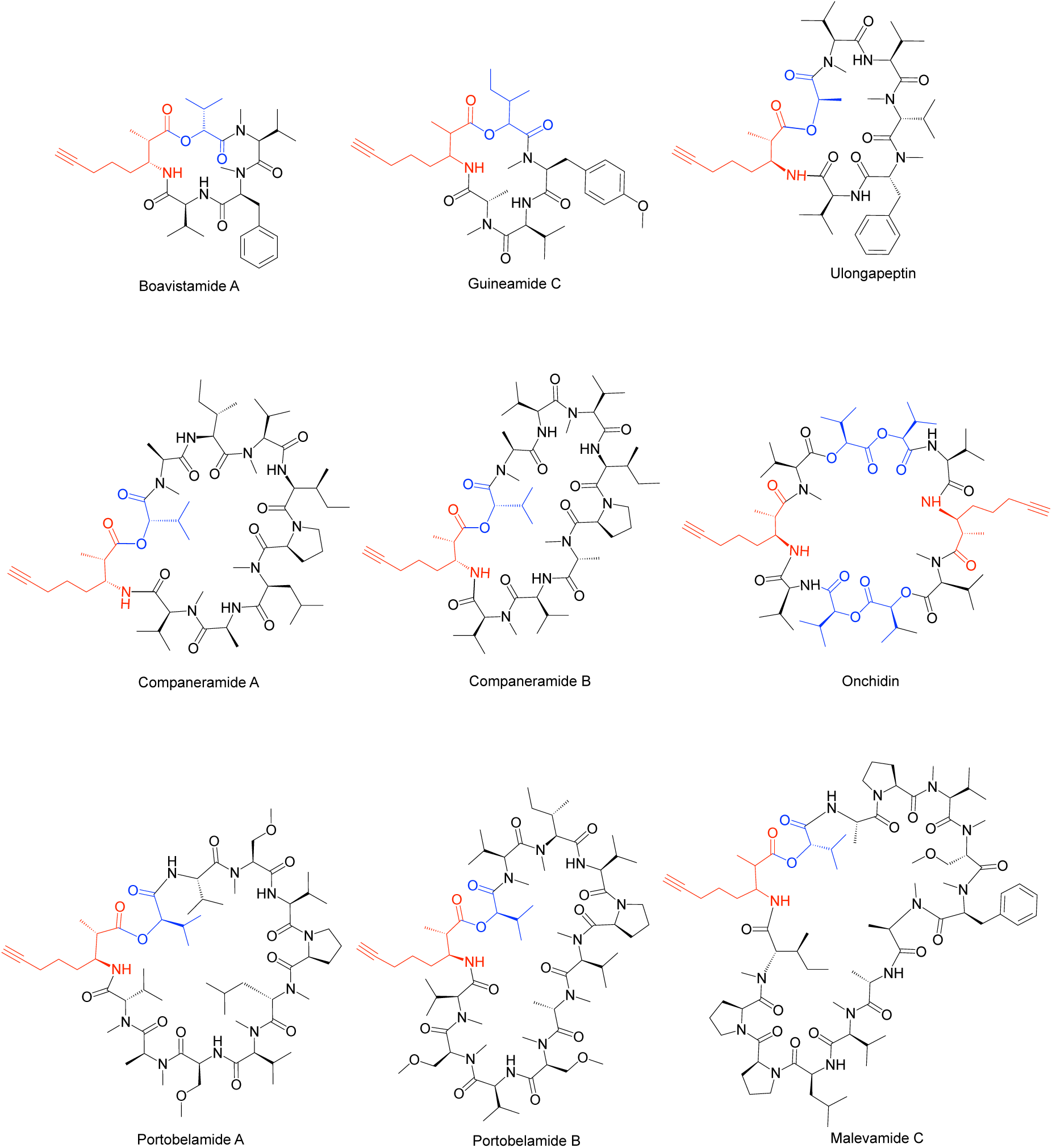
Structural diversity of known AMOYA-containing cyclic depsipeptides. The AMOYA moiety is highlighted in red, and hydroxy acid residues in blue.

In this context, we performed a comparative analysis of all known AMOYA-containing cyclic depsipeptides, based on a comprehensive survey conducted using SciFinder (https://scifinder-n.cas.org/) combined with manual inspection of the literature, to identify both conserved structural features and distinguishing variations (Tables S8–S10, Figures 5 and S47). Despite similarities to the kulolide scaffold, AMOYA-bearing peptides exhibit significantly greater architectural diversity. While DHOYA- and HMOYA-containing peptides are structurally conserved, respectively composed of 6 or 7 residues, AMOYA analogs range from 5 to 14 amino or hydroxy acid residues and display a broader range of structural components. This variability complicates direct sequence alignment and positional comparison with canonical kulolide structures. To date, only nine AMOYA-bearing cyclic depsipeptides have been reported, in contrast to more than 20 examples each for the DHOYA and HMOYA subgroups.^10,11,31,8,7,9^ This limited prevalence, coupled with their broader structural diversity, supports the classification of AMOYA-containing peptides as a chemically related but distinct group, rather than a formal third subgroup of the kulolide superfamily. Boavistamide A, isolated from marine cyanobacteria collected in Cabo Verde, represents not only the first AMOYA-containing compound reported from the African continent, but also matches guineamide C as the smallest macrocycle in this class of NPs.

### F) Cytotoxicity and Antiparasitic Assays

Initial biological screening of the crude extract and its fractions revealed moderate antiplasmodial and antitrypanosomal activity along with weak cytotoxicity toward NCI-H460 cells; antiplasmodial activity was used to prioritize the fractions for the isolation of boavistamides (Figures S5–S8). The purified boavistamides were subsequently evaluated for their antiplasmodial activity against *P. falciparum* Dd2 and for cytotoxicity toward the HEK293T and HepG2 mammalian cell lines. Boavistamides A and B (**1** and **2**) exhibited moderate antiplasmodial activity, with EC₅₀ values of 2.2 and 2.3 μM, respectively, and no detectable cytotoxicity at the maximum concentration tested (11.9 μM), resulting in selectivity indices > 5 and meeting the hit criteria of the screening platform Calibr. Boavistamide C (**3**) displayed slightly lower antiplasmodial activity (EC_50_ = 2.9 μM) and moderate cytotoxicity toward HEK293T cells (CC_50_ = 6.7 μM; SI = 2.3), and, therefore, did not meet the hit criteria (Figures S48–S50, Table S11). Additional screening data, including assays against *T. brucei* and SF188 glioblastoma cells, are provided in the Supporting Information (Figure S51, Table S12). In brief, **1** showed no activity against *T. brucei* at 4 μM and no cytotoxicity was detected toward glioblastoma cells up to 12 μM with 80% cell survival observed at 36 μM.

## CONCLUSIONS

In summary, we report here the discovery of three new cyclic depsipeptides, boavistamides A–C (**1**–**3**), from the marine, filamentous cyanobacterium, *Okeania* sp. that was collected near Boa Vista Island on Ilhéu de Sal Rei, Cabo Verde. These compounds were prioritized for isolation and characterization based on antiparasitic activity and GNPS molecular networking, which highlighted a distinctive molecular family within the Cabo Verde extract. Pure boavistamides A and B (**1** and **2**) displayed moderate antiplasmodial activity with little to no cytotoxicity to three mammalian cell lines. The structures of compounds **1**–**3** were elucidated through an AI-accelerated workflow combining HRMS, comprehensive NMR analyses, and annotation support from SMART-NMR and DeepSAT, enabling rapid and rigorous structural assignment. The absolute configurations of boavistamides A–C (**1**–**3**) were established through a combination of acid hydrolysis, advanced Marfey’s analysis, L-Phe-OMe derivatization, and detailed ^1^H NMR coupling constant and ROESY analysis. Boavistamide A (**1**) also expands the small and structurally diverse family of AMOYA-containing cyclic depsipeptides, a chemically related yet distinct group from the DHOYA/HMOYA-based kulolide superfamily. This study represents the first report of secondary metabolites from a marine cyanobacterium collected in Cabo Verde and highlights the potential of integrating metabolomics and AI-assisted tools to accelerate the discovery and characterization of new bioactive cyanobacterial NPs.

## EXPERIMENTAL SECTION

### A) General Experimental Procedures

Optical rotation values were measured on a JASCO P-2000 polarimeter using a 1 cm microcell. UV spectra were acquired on a Beckman Coulter DU-800 UV–vis spectrophotometer, and IR data were collected on a Nicolet iS50 FT-IR spectrometer (Thermo Fisher Scientific). NMR experiments (^1^H, ^13^C, HSQC, HMBC, and ROESY) were recorded at ambient temperature on two instruments: a Bruker Avance III DRX-600 MHz spectrometer equipped with a 1.7 mm dual-tune TCI cryoprobe, and a JEOL ECZ 500 MHz spectrometer fitted with a 3 mm inverse-detection probe. Chemical shifts were referenced to residual CDCl_3_ signals (δ_H_ 7.26, δ_C_ 77.16). All spectra were processed and analyzed using MestReNova (v14.3.0-30573, Mestrelab Research). LC-MS/MS analyses were performed on a Thermo LTQ XL Finnigan system comprising a Surveyor Autosampler, LC Pump Plus, and PDA Plus detector coupled to an Advantage Max mass spectrometer. Chromatographic separations used a Kinetex 5 μm C18 column (100 Å, 100 × 4.6 mm; Phenomenex). Data were examined using Xcalibur Qual Browser (v1.4 SR1). Analytical and semipreparative HPLC purifications were carried out on a Thermo Scientific Dionex UltiMate 3000 LC system equipped with a diode-array detector and operated through Chromeleon software. Depending on the purification stage, either a Kinetex 5 μm C18 analytical column (100 × 4.6 mm; Phenomenex) or a semipreparative C18 column (150 × 10 mm; Phenomenex) was used. High-resolution mass spectrometric data were collected at the UCSD Chemistry and Biochemistry Mass Spectrometry Facility on an Agilent 6230 TOF mass spectrometer equipped with a Jet Stream ESI source operating in positive-ion mode. The following parameters were applied: capillary voltage, 3500 V; fragmentor, 160 V; nozzle voltage, 500 V; drying and sheath gas temperatures, 325 °C; gas flow rates of 7.0 and 10 L/min, respectively; and nebulizer pressure, 40 psi. All solvents used for extraction, chromatography, and LC-MS/MS analyses were HPLC- or LC-MS-grade (Fisher Chemical). Deuterated solvents for NMR were purchased from Cambridge Isotope Laboratories.

### B) Biological Material, Metagenomic Sequencing, and Taxonomic Identification

Environmental sample 2370 was collected on May 19, 2023, from a rocky intertidal platform on Ilhéu de Sal Rei, Boa Vista Island, Cabo Verde (GPS coordinates: 16°09’50.8” N, 22°55’29.1”W; 16.164114, -22.924750). The material consisted of a dark green, filamentous biofilm attached to the rock surface (Table S1, Figure S1). A subsample was preserved in RNAlater for genomic analysis, while the remaining biomass was used for chemical extraction and compound isolation. Genomic DNA was extracted from RNAlater-preserved material using the DNeasy PowerSoil Kit (Qiagen). Shotgun metagenomic sequencing was performed at BGI using DNBSEQ™ paired-end technology (150 bp), yielding ∼10 Gb of data per sample. Paired-end reads were quality trimmed with BBDuk (Q20 filter; removal of reads <45 bp), and assemblies were generated using metaSPAdes (v3.15.5). Read mapping was performed using Bowtie2 (v2.5.1) and samtools (v1.13).^33,34,35^ Assembly processing, gene calling, functional annotation, profiling, and bin refinement were conducted with Anvi’o (v7.1) following the standard metagenomics workflow.^36^ Binning was carried out with CONCOCT (v1.1.0), followed by manual refinement in Anvi’o.^37^ Refined bins were taxonomically classified using GTDB-Tk (v2.4.0).^38^ Genome completeness and contamination estimates were computed with Anvi’o and CheckM (v1.2.2).^39^ Bins were retained as MAGs if they met standard quality criteria (>90% completeness and <5% contamination for high-quality; >50% completeness and <10% contamination for medium-quality), while bins below these thresholds were excluded from downstream analyses. BGC prediction was performed using antiSMASH (v7.0).^40^

### C) LC-MS Analysis and GNPS2 Classical Molecular Networking

The raw LC-MS/MS data were converted to *.mzXML files and a classical molecular network was created using the online workflow (https://ccms-ucsd.github.io/GNPSDocumentation/) on the GNPS website (http://gnps.ucsd.edu).^20^ The data were filtered by removing all MS/MS fragment ions within +/- 17 Da of the precursor *m/* prior to molecular networking analysis. MS/MS spectra were window filtered by choosing only the top 6 fragment ions in the +/- 50 Da window throughout the spectrum. The precursor ion mass tolerance was set to 2.0 Da with a MS/MS fragment ion tolerance of 0.5 Da. A network was then created where edges were filtered to have a cosine score above 0.6 and more than 6 matched peaks. Further, edges between two nodes were kept in the network if and only if each of the nodes appeared in each other’s respective top 10 most similar nodes. The spectra in the network were then searched against GNPS’ spectral libraries. The library spectra were filtered in the same manner as the input data. All matches kept between network spectra and library spectra were required to have a score above 0.7 and at least 6 matched peaks.

#### Boavistamide A

(**1**). White amorphous solid; [α]_D_^25^ – 45 (*c* 1.0, MeOH); UV (MeOH) λ_max_ (log E) 203 (4.22) nm; IR (film) v_max_ 3314, 2964, 2933, 2873, 1728, 1642, 1497, 1469, 1411, 1388, 1369, 1243, 1178, 1086, 700 cm^-1^; ^1^H and ^13^C NMR data, see Table 1; HRESIMS obs. *m/z* 647.3774 [M + Na]^+^ (calcd for C_35_H_52_N_4_O_6_Na, 647.3785, mass accuracy –1.63 ppm), molecular formula C_35_H_52_N_4_O_6_, 12 degrees of unsaturation (Figures S13, S53 and S54).

**Table 1.**
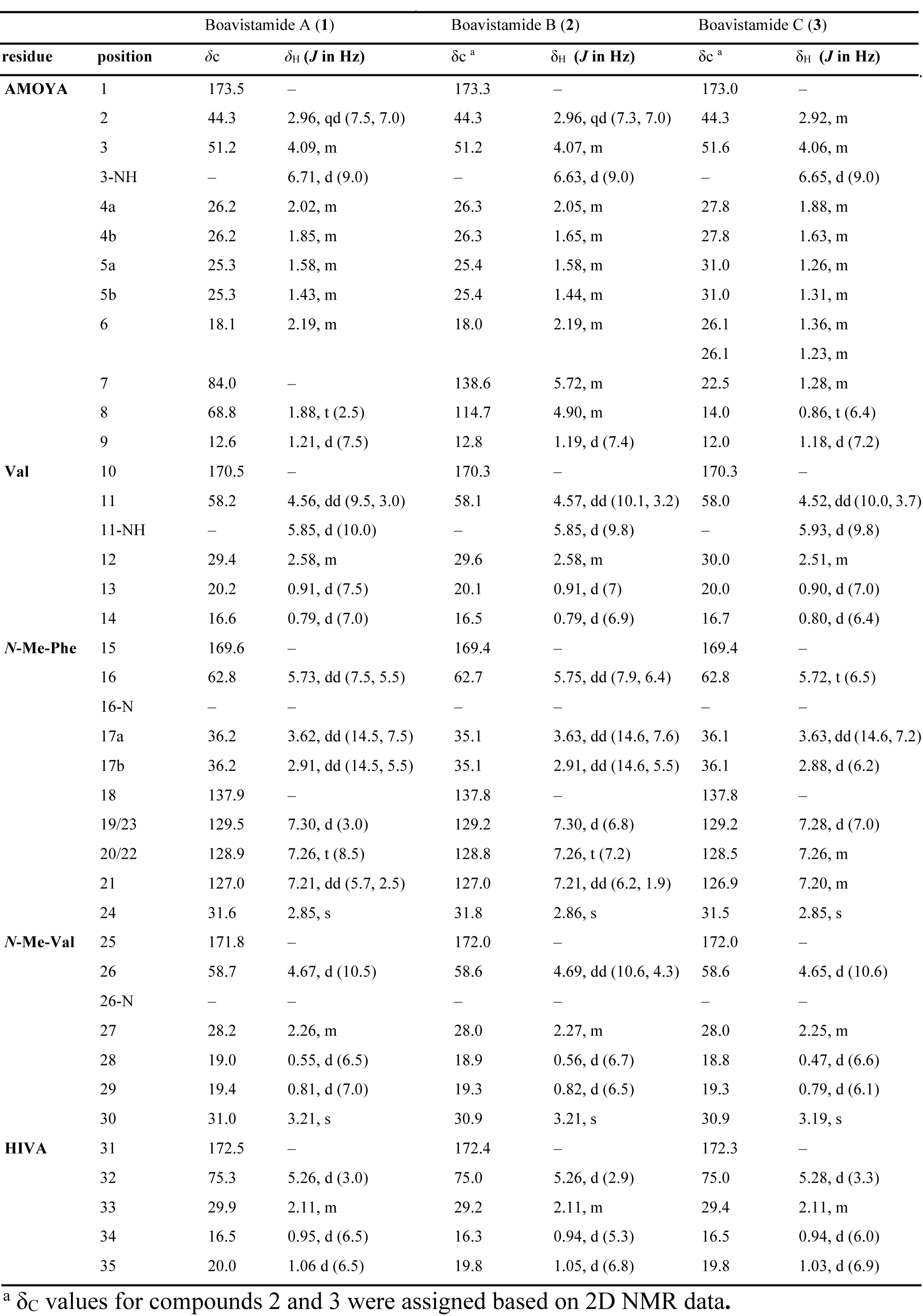
NMR Spectroscopic Data for Boavistamides A–C (1–3) (^¹^H 500 MHz for 1 and 2, 600 MHz for 3; ^¹³^C 125 MHz for 1; CDCl_₃_)

#### Boavistamide B

(**2**). White amorphous solid; [α]_D_^28^ – 12 (*c* 1.0, MeOH) ; UV (MeOH) λ_max_ (log E) 204 (4.11) nm ; IR (film) v_max_ 3420, 2935, 1617, 1498, 1403, 1356, 1255, 1143, 1057 cm^-1^; ^1^H and ^13^C NMR data, see Table 1; HRESIMS obs. *m/z* 649.3929 [M + Na]^+^ (calcd for C_35_H_54_N_4_O_6_Na, 649.394, mass accuracy –1.86 ppm), molecular formula C_35_H_54_N_4_O_6_, 11 degrees of unsaturation (Figures S20, S55 and S56).

#### Boavistamide C

(**3**). White amorphous solid; [α]_D_^26^ – 7 (*c* 1.0, MeOH) ; UV (MeOH) λ_max_ (log E) 205 (4.07) nm ; IR (film) v_max_ 3316, 2953, 2841, 1646, 1450, 1111, 1013 cm^-1^; ^1^H and ^13^C NMR data, see Table 1; HRESIMS obs. *m/z* 651.4092 [M + Na]^+^ (calcd for C_35_H_56_N_4_O_6_Na, 651.4098, mass accuracy –0.85 ppm), molecular formula C_35_H_56_N_4_O_6_, 10 degrees of unsaturation (Figures S26, S57 and 58).

### D) Total Hydrolysis and Advanced Derivatization Analysis

#### Acid Hydrolysis

Boavistamides A–C (**1**–**3**) (0.2 mg; 0.1 mg in each of two vials, one for Marfey’s derivatization and one for L-Phe-OMe derivatization) were hydrolyzed with 6 N HCl (200 μL) in sealed vials at 110 °C for 16 h.

#### Marfey’s Analysis

The hydrolyzed samples were dried and resuspended in H₂O (100 μL), followed by addition of a 1% acetone solution of *N*-α-(2,4-dinitro-5-fluorophenyl)-L-leucinamide (L-FDLA; 500 μL), 1 M NaHCO_3_ (100 μL), and DMSO (50 μL). Authentic standards, including D-Val, L-Val, *N*-Me-D-Phe, *N*-Me-L-Phe, *N*-Me-D-Val, and *N*-Me-L-Val, were prepared as 50 mM solutions (50 μL each in 5 mL vials). Each standard solution received a 1% acetone solution of L-FDLA (100 μL), 1 M NaHCO_3_ (20 μL), and DMSO (10 μL). All mixtures were stirred at 40 °C for 1 h, cooled to room temperature, and quenched with 2 M HCl (50 μL for the boavistamide A-derived sample, 10 μL for the standards). Samples were dried under a nitrogen stream and reconstituted in MeOH (1 mL for the sample, 10 mL for standards). After filtration through a 0.2 μm nylon membrane, samples were analyzed by LC-MS (Kinetex 5 μm C18 100 Å column, 100 × 4.6 mm,) using a linear gradient from 30:70 to 99:1 MeCN/H_2_O (0.1% formic acid) over 32 min at a flow rate of 0.75 mL/min in positive ion mode. Data acquisition was performed using Xcalibur software on a Thermo Scientific LTQ XL mass spectrometer. Retention times of derivatized amino acids from the sample were compared with those of the authentic standards to determine the absolute configurations.

#### L-Phe-OMe Derivatization

Authentic standards of D- and L-hydroxyisovaleric acid (HIVA, 11 mg, 0.093 mmol) were individually coupled with L-phenylalanine methyl ester hydrochloride (L-Phe-OMe, 45 mg, 0.21 mmol) using O-(benzotriazol-1-yl)-*N*,*N*,*N*′,*N*′-tetramethyluronium hexafluorophosphate (HBTU, 38 mg, 0.10 mmol) and *N*,*N*-diisopropylethylamine (DIPEA, 100 μL) in DMSO (1 mL). Reactions were stirred overnight at room temperature. Crude reaction mixtures were loaded onto preconditioned Sep-Pak C18 cartridges, washed with H_2_O to remove residual DMSO, and eluted with MeOH. The eluted products were dried under N_2_ and analyzed by LC-MS as described above. For the NPs, boavistamides A–C (0.1 mg each) were derivatized under identical conditions using L-Phe-OMe (4.5 mg, 0.021 mmol), HBTU (3.8 mg, 0.010 mmol), DIPEA (100 μL), and DMSO (100 μL). After overnight reaction, the mixtures were dried directly under nitrogen and reconstituted in MeOH. Samples were filtered through a 0.2 μm nylon membrane and analyzed by LC-MS under the same chromatographic conditions. Retention times and *m/z* values were compared to those of authentic derivatives to determine the absolute configuration of the HIVA moiety.

## DATA AVAILABILITY

The NMR data for boavistamides A–C (**1**–**3**) have been deposited in the Natural Products Magnetic Resonance Database under the accession numbers NP0354020, NP0354021 and NP0354022. Raw metagenomic sequencing data have been deposited in the European Nucleotide Archive (ENA) under project accession number PRJEB110160. The GNPS molecular networking analysis is publicly available at https://gnps2.org/status?task=b11800555aba447eb5be1e0f3ef1971a#

## SUPPORTING INFORMATION

Sample collection details; LC-MS chromatograms and GNPS-based molecular networking analysis; metagenomic binning and taxonomic analysis; ^1^H and ^13^C NMR, COSY, HSQC, HMBC, and ROESY spectra for compounds **1**–**3**; HRESIMS data; chiral derivatization and Marfey’s analysis data; conformational analysis of the AMOYA unit; biological activity and cytotoxicity assay data.

## NOTE

The authors declare no competing financial interest.

## ACKNOWLEDGMENTS

The authors thank B. Duggan (UC San Diego) for assistance with the acquisition of 600 MHz NMR data and Y. Su (UC San Diego) for assistance with HRMS data acquisition. We are grateful to the Dickinson Foundation for the purchase of the JEOL ECZ 500 MHz NMR spectrometer. Cyanobacterial sampling in Cabo Verde was conducted within the framework of the EMERTOX project, co-funded by the European Union’s Horizon 2020 research and innovation programme under the Marie Skłodowska-Curie grant agreement No. 778069. We gratefully acknowledge J. Morais, J. Almeida, and F. Oliveira for assistance with sample collection. K. Rodriguez (Calibr) assisted with preparation of assay ready plates for bioactivity screening efforts. We acknowledge ACD/Labs for providing access to the HSQC prediction tool, which was instrumental in building the datasets used to train SMART 2.1 and DeepSAT. We thank Dr. J. Diaz (UC San Diego) for the use of SpectraMax M3 microplate reader (Molecular Devices). This work was supported by a PhD scholarship (N° 2024.03345.BD) from Fundação para a Ciência e a Tecnologia (FCT, Portugal) to M.C. This project received funding to P.N.L. from the European Union (ERC, GREASEDLIGHTNING, grant agreement N° 101088806), and from the European Union’s Horizon 2020 research and innovation programme under grant agreement N° 952374 (BlueBio4Future) and the Bill and Melinda Gates Foundation (INV-037899)(WHG). This work was also supported by Fundação para a Ciência e a Tecnologia (FCT, Portugal) to A.R., under the scope of the project 2023.15022.PEX (https://doi.org/10.54499/2023.15022.PEX), including computational resources at Deucalion supercomputer (2023.15022.PEX.F1). Research reported in this publication was also supported in part by the National Center for Complementary and Integrative Health of the NIH under award number F32AT011475 to N.E.A and R21AI171824 to C.R.C.

