## Supporting Information for "AI-Accelerated Structure Elucidation of Boavistamides A–C, Cyclic Depsipeptides from a Marine Filamentous Cyanobacterium Collected in Cabo Verde"

**Table S2.** Bin 81 relative abundance, completeness, contamination, GC and genome size .. . ..........................................................................................................................9

### Figures

**Figure S1.** Geographic origin of environmental cyanobacterial sample collected in Insular West Africa....... ........................................................................................................20

**Figure S33.** Scheme for absolute configuration determination of HIVA, Valine, *N*-Me-phenylalanine and *N*-Me-Valine of boavistamides A-C (**1**–**3**) by derivatization methods. 52

**Method S1.** Preliminary bioactivity screening of extract and VLC fractions.

Preliminary *in vitro* antiparasitic, antimicrobial, and cytotoxicity screenings were conducted on crude extract and vacuum liquid chromatography (VLC) fractions from cyanobacterial sample 2370 to prioritize fractions for subsequent isolation. The initial bioactivity panel included *Plasmodium falciparum* growth inhibition assays (Plouffe et al., 2008), *Trypanosoma brucei brucei* growth inhibition assays (Chirwa et al., 2024), and antimicrobial assays against *Staphylococcus aureus*, *Escherichia coli*, *Bacillus subtilis*, *Salmonella typhimurium*, and *Candida albicans* using an agar-based disk diffusion method (Girão et al., 2019) and cytotoxicity assays against mammalian cell lines, including NCI-H460 human lung carcinoma cells (Avalon et al., 2024) and HEK293 human embryonic kidney cells using a resazurin-based viability assay (O'Brien et al., 2000). VLC fractions 2370-D and 2370-E showed the most relevant bioactivity profiles and were therefore selected for further purification (Figures S5–S8 and Table S3).

**Method S2.** Biological evaluation of boavistamides A–C (**1**–**3**).

*S2.1 In Vitro Antiparasitic Assays*

*Anti-plasmodium assay*

*Plasmodium falciparum* Dd2 parasites were cultured in RPMI1640 (Gibco) supplemented with L-glutamine, 0.5% Albumax, 25 mM HEPES, and 50 mg/L hypoxanthine at 2% hematocrit in human O+ erythrocytes. Cultures were maintained at 37 °C in a malaria gas mixture of 90% $N_2$,5% $CO_2$, 5% $O_2$. *In vitro* antimalarial activity of boavistamides A–C (**1**–**3**) was evaluated in a dose-response assay using a three-fold serial dilution with 12 concentrations in a 1536-well plate. The assay plate was incubated for 72 hours. Parasite proliferation was measured using 2x SYBR Green I DNA/RNA staining diluted in a cell lysis buffer (20 mM Tris-HCl, 5 mM EDTA, 0.08% (v/v) saponin, 0.8% (v/v) Triton X-100) (Plouffe et al., 2008). The plate was incubated overnight in the dark to complete lysis and staining, and fluorescence was measured at 485/530 nm. Data were normalized to DMSO and 3.125 µM artesunate controls. (Figures S48–S50). The assays were performed in one biological replicate that had two technical replicates.

*Anti-trypanosomal assay*

*Trypanosoma brucei brucei* (Lister 427) was cultured at 37 °C under a humidified 5% $CO^2$ atmosphere in HMI-9 modified medium (Hirumi and Hirumi, 1989) supplemented with 10% heat-inactivated fetal bovine serum (FBS) (Monti et al., 2023). The anti-trypanosomal assay was performed as described (Chirwa et al., 2024). Briefly, boavistamide A (**1**) was diluted in DMSO and added to 96-well polystyrene assay plates to give a final assay concentration of 4 µM for boavistamide A (1 µL; 0.5% total DMSO). Fresh HMI-9 medium (99 µL/well) was added to the assay plate. Parasites in the exponential phase were suspended at $2 \times 10^5$ parasites/mL in HMI-9 medium and added to each well (100 µL) to a total density of $2 \times 10^4$ trypanosomes/well. Assay plates were incubated at 37 °C and 5% $CO_2$ for 70 h, followed by the addition of 20 µL/well of 0.5 mM resazurin. The plates were incubated for an additional 2 h, and fluorescence was measured at 535 nm and 590 nm excitation and emission wavelengths, respectively, using a 2104 EnVision® multilabel plate reader (PerkinElmer, Waltham, MA). The viability of the parasites was normalized to positive and negative controls in each assay plate. The screening was performed in technical quadruplicate, and pentamidine at a fixed concentration of 4 μM was employed as positive drug control. (Table S12)

### S2.2 In Vitro Mammalian Cell Cytotoxicity Assays

#### *HepG2 and HEK293T mammalian cell lines*

HepG2 (ATCC) and HEK293T (ATCC) mammalian cell lines were maintained in Dulbecco's Modified Eagle Medium (DMEM, Gibco) with 10% heat-inactivated HyClone FBS (GE Healthcare Life Sciences), 100 IU penicillin, and 100 mg/mL streptomycin (Gibco) at 37°C with 5% $CO_2$ in a humidified tissue culture incubator. To assay mammalian toxicity of boavistamides A–C (**1**–**3**), 250 HepG2 or 375 HEK293T cells/well were seeded in assay media (DMEM, 2 % FBS, 100 IU penicillin, and 100 mg/mL streptomycin) in 1536-well, white, tissue culture-treated, solid bottom plates (Corning, 9006BC) that contained acoustically transferred compounds in a three-fold serial dilution starting at 12 μM. After 72-h of incubation, CellTiter-Glo Luminescent Cell Viability Assay (Promega) was used to quantify cell viability as per manufacturer instructions. Luminescence signal was read on the CLARIOstar Plus plate reader (BMG Labtech). Data were normalized to DMSO and 10 μM puromycin controls. (Figure S48–S50)

#### *SF188 human glioblastoma and NCI-H460 human lung carcinoma cell lines*

SF188 human glioblastoma cells (Sigma, Cat. #SCC282) and NCI-H460 human lung carcinoma cells (ATCC, NCI-H460, HTB-177 ™) were cultured as monolayers in Minimum Essential Medium (MEM) or Eagle's Minimum Essential Medium (EMEM, 30-2002) supplemented with 10% fetal bovine serum (Sigma, ES-009-B or (ATCC, 30-2020), 2 mM L-glutamine (TMS-002-C) (SF188), and 1% penicillin–streptomycin (Hyclone, SV-300-10) until approximately 90% confluence. Cells were then detached using accutase (SF188) or 0.25% trypsin (NCI-H460) solutions and seeded into 96 wp at 6,000 cells/well. 24 hours later either extracts (two concentrations of 10 and 1 μg/mL) or boavistamide (ten half-log concentrations, starting from 36 μM) were added to the cells. Quisinostat was used as positive control. 48-hour later cells were stained with MTT ((3-(4,5-dimethylthiazol-2-yl)-2,5-diphenyl tetrazolium bromide, 0.83 mg/mL) for either 60 or 25 min and analyzed in comparison with the negative control (100% complete medium set to represent 100% cell viability) at 570 nm on SpectraMax M3 microplate reader (Molecular Devices). OD values at 630 nm were subtracted as the background. (Avalon et al., 2024). (Figures S5 and S51)

**Table S1.** Sampling information and field observations for the cyanobacterial biofilm collected on Ilhéu de Sal Rei, Boa Vista Island, Cabo Verde.

| Sample Code | Sample ID (Alias) | Collection Date | Collection Site | GPS Coordinates | Site Description | Field Macroscopic Observation | Field Microscopic Observation | Metagenomics Sample |
|---|---|---|---|---|---|---|---|---|
| BV043 | 2370 | May 19, 2023 | Ilhéu de Sal Rei, Boavista Island, Cabo Verde | 16°09'50.8"N 22°55'29.1"W \| 16.164114, -22.924750 | Rocky platform (intertidal zone) | Dark green biofilm attached to the rock | Mixed cyanobacterial community (Oscillatoriales-dominant, Nostocales present) | Yes |

**Table S2.** Bin 81 relative abundance, completeness, contamination, GC and genome size.

| Bin | Taxonomic classification (GTDB-Tk) | Relative abundance (Anvi'o, %) | Completeness (Check m, %) | Contamination (Check m, %) | GC (Check m, %) | GC std (Check m, %) | Genome size (Check m, bp) |
|---|---|---|---|---|---|---|---|
| 81 | d__Bacteria;p__Cyanobacteriota;c__Cyanobacteriia;o__Cyanobacteriales;f__Microcoleaceae;g__Okeania;s | 4.46 | 94.65 | 3.93 | 36 | 2 | 7,628,354 |

**Table S3.** Antimicrobial activity of the VLC fractions (A–I) from cyanobacterial sample 2370, evaluated by agar disc diffusion assay against a panel of reference microorganisms: *Staphylococcus aureus* (ATCC 29213), *Bacillus subtilis* (ATCC 6633), *Escherichia coli* (ATCC 25922), *Salmonella typhimurium* (ATCC 25241), and *Candida albicans* (ATCC 10231) at a test quantity of 15 µg/disc. A dash indicates no zone of inhibition.

| **Strains** | **2370 fractions** | | | | | | | | | |
|---|---|---|---|---|---|---|---|---|---|---|
| | A | B | C | D | E | F | G | H | Hx | I |
| *Candida albicans* | — | — | — | — | — | — | — | — | — | — |
| *Bacillus subtilis* | 2mm | — | — | — | — | — | — | — | — | — |
| *Salmonella typhimurium* | — | — | — | — | — | — | — | — | — | — |
| *Staphylococcus aureus* | — | — | — | — | — | — | — | — | — | — |
| *Escherichia coli* | — | 2mm | — | — | — | — | — | — | — | — |

**Table S4.** NMR Spectroscopic Data of Boavistamide A (**1**) ($^1$H 500 MHz and $^{13}$C 125 MHz, $CDCl_3$)

| residue | position | $\delta_C$ | $\delta_H$ (*J* in Hz) | COSY | HMBC | ROESY |
|---|---|---|---|---|---|---|
| **AMOYA** | 1 | 173.5 | – | – | 2, 3, 9, 32 | – |
| | 2 | 44.3 | 2.96, qd (7.5, 7.0) | 3, 9 | 1, 3, 4, 9 | 3, 4a, 9, 14 |
| | 3 | 51.2 | 4.09, m | 2, 3-NH, 4 | 1 | 2, 4b, 5b, 6, 9 |
| | 3-NH | – | 6.71, d (9.0) | 3 | – | 4a, 11, 11-NH |
| | 4a | 26.2 | 2.02, m | 3, 5 | 3, 5 | 3, 9, 34 |
| | 4b | 26.2 | 1.85, m | 3, 5 | 5, 6, 7 | 3, 9 |
| | 5a | 25.3 | 1.58, m | 4, 6 | 8 | 4a |
| | 5b | 25.3 | 1.43, m | 4, 6 | 3, 4, 6, 7 | 4a, 4b |
| | 6 | 18.1 | 2.19, m | 5 | 4, 5, 7, 8 | 5b |
| | 7 | 84.0 | – | – | 5a, 5b, 6, 8 | – |
| | 8 | 68.8 | 1.88, t (2.5) | – | 5, 6, 7 | 3, 9 |
| | 9 | 12.6 | 1.21, d (7.5) | 2 | 1, 2, 3 | 2, 3, 4b |
| **Val** | 10 | 170.5 | – | – | 3-NH | – |
| | 11 | 58.2 | 4.56, dd (9.5, 3.0) | 12 | 10, 13, 14 | 13 |
| | 11-NH | – | 5.85, d (10.0) | – | 11, 15 | 14, 3-NH |
| | 12 | 29.4 | 2.58, m | 11, 13, 14 | 13, 14 | 11, 13, 14 |
| | 13 | 20.2 | 0.91, d (7.5) | 12 | 11, 12, 14 | 11, 12, 24 |
| | 14 | 16.6 | 0.79, d (7.0) | 12 | 11, 12, 13 | 11-NH, 12, 24 |
| ***N*-Me-Phe** | 15 | 169.6 | – | – | 11-NH, 16, 17a, 17b | – |
| | 16 | 62.8 | 5.73, dd (7.5, 5.5) | 17 | 15, 17, 18, 24, 26 | 11-NH |
| | 16-N | – | – | – | – | – |
| | 17a | 36.2 | 3.62, dd (14.5, 7.5) | 16 | 16, 15, 18, 19 | 16, 17b |
| | 17b | 36.2 | 2.91, dd (14.5, 5.5) | 16 | 16, 15, 18, 19 | 17a |
| | 18 | 137.9 | – | – | 16, 17, 19/23, 20/22 | – |
| | 19/23 | 129.5 | 7.30, d (3.0) | – | 20/22, 21 | 16, 17a/b, 26, 28 |
| | 20/22 | 128.9 | 7.26, t (8.5) | – | 18, 19 | 12 |
| | 21 | 127.0 | 7.21, dd (5.7, 2.5) | – | – | – |
| | 24 | 31.6 | 2.85, s | – | 16, 25 | 28, 29 |
| ***N*-Me-Val** | 25 | 171.8 | – | – | 16, 24, 26 | – |
| | 26 | 58.7 | 4.67, d (10.5) | 27 | 25, 27, 30 | 16, 27, 28, 29 |
| | 26-N | – | – | – | – | – |
| | 27 | 28.2 | 2.26, m | 26, 28, 29 | 26, 28 | 26, 28, 29, 30 |
| | 28 | 19.0 | 0.55, d (6.5) | 27 | 26, 27, 29 | 26, 27 |
| | 29 | 19.4 | 0.81, d (7.0) | 27 | 26, 27, 28 | 26, 29, 30, 31 |
| | 30 | 31.0 | 3.21, s | – | 26, 31 | 27,29, 32, 33 |
| **HIVA** | 31 | 172.5 | – | – | 30 | – |
| | 32 | 75.3 | 5.26, d (3.0) | 33 | 1, 33, 34, 35 | 30, 33, 35 |
| | 33 | 29.9 | 2.11, m | 32, 34, 35 | 34, 35 | 31, 32, 34, 35 |
| | 34 | 16.5 | 0.95, d (6.5) | 33 | 32, 33, 35 | 33 |
| | 35 | 20.0 | 1.06 d (6.5) | 33 | 32, 33, 34 | 32, 33 |

**Table S5.** NMR Spectroscopic Data of Boavistamide B (**2**) ($^1$H 500 MHz, $CDCl_3$)

| residue | position | δc [a] | δH (*J* in Hz) | COSY | HMBC | ROESY |
|---|---|---|---|---|---|---|
| AMOYA | 1 | 173.3 | – | – | 2, 9 | – |
| | 2 | 44.3 | 2.96, qd (7.3, 7.0) | 9 | 1, 3, 9 | 8, 9 |
| | 3 | 51.2 | 4.07, m | 2, 3-NH, 4 | 2, 9 | 2, 9 |
| | 3-NH | – | 6.63, d (9.0) | 3 | 10 | – |
| | 4a | 26.3 | 2.05, m | 3, 4 | 7, 8 | – |
| | 4b | 26.3 | 1.65, m | 3, 4 | – | – |
| | 5a | 25.4 | 1.58, m | – | – | – |
| | 5b | 25.4 | 1.44, m | – | – | – |
| | 6 | 18.0 | 2.19, m | 5 | 5 | – |
| | 7 | 138.6 | 5.72, m | 8 | – | – |
| | 8 | 114.7 | 4.9, m | 7 | – | – |
| | 9 | 12.8 | 1.19, d (7.4) | 2 | 1, 2, 3 | – |
| Val | 10 | 170.3 | – | – | 3-NH, 11 | – |
| | 11 | 58.1 | 4.57, dd (10.1, 3.2) | 11-NH | 10, 12, 14 | 12, 13, 14 |
| | 11-NH | – | 5.85, d (9.8) | – | 11, 15 | 13 |
| | 12 | 29.6 | 2.58, m | 13, 14 | 11, 13, 14 | 13, 14 |
| | 13 | 20.1 | 0.91, d (7.0) | 12 | 11, 12, 14 | – |
| | 14 | 16.5 | 0.79, d (6.9) | 12 | 11, 12, 13 | – |
| N-Me-Phe | 15 | 169.4 | – | – | 16, 17a | – |
| | 16 | 62.7 | 5.75, dd (7.9, 6.4) | 17a, 17b | 15, 18, 24, 25 | 26, 19/23 |
| | 16-N | – | – | – | – | – |
| | 17a | 35.1 | 3.63, dd (14.6, 7.6) | 16, 17b | 15, 16, 17b, 19/23, 18 | – |
| | 17b | 35.1 | 2.91, dd (14.6, 5.5) | 16, 17a | 15, 16, 19/23 | – |
| | 18 | 137.8 | – | – | 16, 19/23 | – |
| | 19/23 | 129.2 | 7.30, d (6.8) | – | 20/22 | – |
| | 20/22 | 128.8 | 7.26, t (7.2) | – | 18 | – |
| | 21 | 127.0 | 7.21, dd (6.2, 1.9) | – | 19/23 | – |
| | 24 | 31.8 | 2.86, s | – | 25 | – |
| N-Me-Val | 25 | 172.0 | – | – | 26 | – |
| | 26 | 58.6 | 4.69, dd (10.6, 4.3) | 27, 28, 29 | 25, 27, 28, 30 | 16, 29 |
| | 26-N | – | – | – | – | – |
| | 27 | 28.0 | 2.27, m | 26, 29 | 26, 28 | 16, 29 |
| | 28 | 18.9 | 0.56, d (6.7) | 27 | 26, 27, 29 | – |
| | 29 | 19.3 | 0.82, d (6.5) | 27 | 27, 28 | – |
| | 30 | 30.9 | 3.21, s | – | 31 | – |
| HIVA | 31 | 172.4 | – | – | 30 | – |
| | 32 | 75.0 | 5.26, d (2.9) | 33 | 31, 33, 34 | 30, 33, 35 |
| | 33 | 29.2 | 2.11, m | 34, 35 | 34, 35 | 30, 34, 35 |
| | 34 | 16.3 | 0.94, d (5.3) | 33 | 32, 33, 35 | – |
| | 35 | 19.8 | 1.05, d (6.8) | 33 | 32, 33, 34 | – |

[a]assigned by 2D NMR analysis

**Table S6**. NMR Spectroscopic Data of Boavistamide C (**3**) ($^1$H 600 MHz, $CDCl_3$)

| residue | position | δc [a] | δH (*J* in Hz) | COSY | HMBC | ROESY |
|---|---|---|---|---|---|---|
| **AMOYA** | 1 | 173.0 | – | – | 2 | – |
| | 2 | 44.3 | 2.92, m | 9 | 2, 9 | – |
| | 3 | 51.6 | 4.06, m | 3-NH, 5 | – | 2, 9 |
| | 3-NH | – | 6.65, d (9.0) | 3, 5 | 10 | – |
| | 4a | 27.8 | 1.88, m | 3 | 2 | – |
| | 4b | 27.8 | 1.63, m | 3 | – | – |
| | 5a | 31.0 | 1.26, m | – | 8 | – |
| | 5b | 31.0 | 1.31, m | – | – | – |
| | 6a | 26.1 | 1.36, m | – | – | – |
| | 6b | 26.1 | 1.23, m | – | – | – |
| | 7 | 22.5 | 1.28, m | 8 | 8 | – |
| | 8 | 14.0 | 0.86, t (6.4) | 7 | – | – |
| | 9 | 12.0 | 1.18, d (7.2) | 8 | 1, 2, 3 | – |
| **Val** | 10 | 170.3 | – | – | 3-NH, 11 | – |
| | 11 | 58.0 | 4.52, dd (10.0, 3.7) | 11-NH | 10, 12 | 12, 13 |
| | 11-NH | – | 5.93, d (9.8) | 11 | 15 | – |
| | 12 | 30.0 | 2.51, m | 13, 14 | – | 13, 14 |
| | 13 | 20.0 | 0.90, d (7.0) | 12 | 11, 14 | – |
| | 14 | 16.7 | 0.80, d (6.4) | 12 | 11, 12 | – |
| **N-Me-Phe** | 15 | 169.4 | – | – | 16, 17a, 17b | – |
| | 16 | 62.8 | 5.72, t (6.5) | 17a, 17b | 15, 18, 17 | 26 |
| | 16-N | – | – | – | – | – |
| | 17a | 36.1 | 3.63, dd (14.6, 7.2) | 16 | 15, 16, 18, 19/23 | – |
| | 17b | 36.1 | 2.88, d (6.2) | 16, 17a | 15, 16, 18, 19/23 | – |
| | 18 | 137.8 | – | – | 16, 21, 17a, 17b | – |
| | 19/23 | 129.2 | 7.28, d (7.0) | – | – | – |
| | 20/22 | 128.5 | 7.26, m | – | 17, 21 | – |
| | 21 | 126.9 | 7.20, m | – | 19/23 | – |
| | 24 | 31.5 | 2.85, s | – | 16, 25 | – |
| **N-Me-Val** | 25 | 172.0 | – | – | – | – |
| | 26 | 58.6 | 4.65, d (10.6) | 27, 28, 29 | 28, 30 | – |
| | 26-N | – | – | – | – | – |
| | 27 | 28.0 | 2.25, m | 26, 28, 29 | 26, 29 | 29, 30 |
| | 28 | 18.8 | 0.47, d (6.6) | 27 | 26, 27, 29 | – |
| | 29 | 19.3 | 0.79, d (6.1) | 27 | 26, 27, 28 | – |
| | 30 | 30.9 | 3.19, s | – | 26, 31 | – |
| **HIVA** | 31 | 172.3 | – | – | 26, 30 | – |
| | 32 | 75.0 | 5.28, d (3.3) | 33 | 31 | 30, 33, 35 |
| | 33 | 29.4 | 2.11, m | 34, 35 | 34, 35 | 30, 32, 34 |
| | 34 | 16.5 | 0.94, d (6.0) | 33 | 32, 33, 35 | – |
| | 35 | 19.8 | 1.03, d (6.9) | 33 | 32, 33, 34 | – |

[a]assigned by 2D NMR analysis

**Table S7.** Comparison of 1D NMR data and ROESY correlations of the AMOYA unit

| | Boavistamide A | Companeramide A | Portobelamide B | Ulongapeptin |
|---|---|---|---|---|
| δH-2 / δC-2 | 2.96 / 44.3 | 2.51 / 46.4 | 2.65 / 41.7 | 2.82 / 41.8 |
| δH-9 / δC-9 | 1.21 / 12.6 | 1.31 / 14.2 | 0.84 / 7.7 | 1.13 / 14.7 |
| δH-3 / δC-3 | 4.09 / 51.2 | 3.97 / 51.2 | 4.43 / 49.8 | 4.27 / 49.4 |
| $J_{2-3}$ (Hz) | 7 | 7.2 | 2.4 | 3.6 |
| Configuration | *erythro* (2*S*,3*R*) | *erythro* (2*S*,3*R*) | *threo* (2*S*,3*S*) | *threo* (2*S*,3*S*) |
| Method | J + ROESY | Marfey's | J + ROESY | Marfey's |
| H2 ↔ H3 (ROESY) | Yes | Yes | Yes | Yes |
| H2 ↔ $CH_3$–9 (ROESY) | Yes | Yes | Yes | Yes |
| NH ↔ $CH_3$–9 (ROESY) | No | No | Yes | Yes |
| H3↔ H4a/H4b (ROESY) | Yes | Yes | Yes | Yes |
| H3 ↔ 5b, 6 (ROESY) | Yes | Yes | No | No |
| NH ↔ H11 (ROESY) | Yes | Yes | Yes | Yes |

**Table S8.** Comparative Structural Features of DHOYA-, HMOYA-, and AMOYA-Containing Cyclic Depsipeptides

| | **DHOYA** | **HMOYA** | **AMOYA** |
|---|---|---|---|
| Number of known compounds | 21 | 25 | 9 |
| Residue count | 7 | 6 | 5 to 14 |
| Macrocycle type | Conserved | Conserved | Variable |
| BGC identified | No | No | No |

**Table S9**. Comparison of residues of AMOYA-containing cyclic depsipeptides.

| Compound name | Residue 1 | Residue 2 | Residue 3 | Residue 4 | Residue 5 | Residue 6 | Residue 7 | Residue 8 | Residue 9 | Residue 10 | Residue 11 | Residue 12 | Residue 13 | Residue 14 |
|---|---|---|---|---|---|---|---|---|---|---|---|---|---|---|
| Malevamide C | AMOYA | HIVA | Ala | Pro | N-Me-Val | N,O-diMeSer | N-Me-Phe | N-Me-Ala | Ala | N-Me-Val | Leu | Pro | Pro | N-Me-Ile |
| Portobelamide B | AMOYA | HIVA | N-Me-Val | N-Me-Ile | Val | Pro | N-Me-Val | N-Me-Ala | N,O-diMeSer | Val | N,O-diMeSer | N-Me-Val | | |
| Portobelamide A | AMOYA | HIVA | Val | N,O-diMeSer | Val | Pro | N-Me-Leu | N-Me-Val | O-Me-Ser | N-Me-Ala | N-Me-Val | | | |
| Onchidin | AMOYA | N-Me-Val | HIVA | HIVA | Val | AMOYA | N-Me-Val | HIVA | HIVA | Val | | | | |
| Companeramide B | AMOYA | HIVA | N-Me-Ala | Val | N-Me-Val | Ile | Pro | N-Me-Ala | Val | N-Me-Val | | | | |
| Companeramide A | AMOYA | HIVA | N-Me-Ala | Ile | N-Me-Val | Ile | Pro | N-Me-Leu | Ala | N-Me-Val | | | | |
| Ulongapeptin | AMOYA | Lactic Acid | N-Me-Val | Val | N-Me-Val | N-Me-Phe | Val | | | | | | | |
| Guineamide C | AMOYA | HMPA | N,O-diMeTyr | Val | N-Me-Ala | | | | | | | | | |
| Boavistamide A | AMOYA | HIVA | N-Me-Val | N-Me-Phe | Val | | | | | | | | | |

**Table S10.** AMOYA-containing cyclic depsipeptides, source organisms, sampling locations and bioactivity.

| Compound name | Source organism | Sampling location | Bioactivity |
|---|---|---|---|
| Malevamide C | *Symploca laete-viridis* | Hawaii | Not cytotoxic against P-388, A-549, and HT-29 |
| Portobelamide B | *Caldora sp.* | Panama | Not cytotoxic against H-460 (97% survival at 0.9 μM) |
| Portobelamide A | *Caldora sp.* | Panama | Cytotoxicity against H-460 (33% survival at 0.9 μM) |
| Onchidin | *Onchidium sp.* | New Caledonia | Fraction cytotoxic against Kb cells (93% inhibition at 10 μg/mL) |
| Companeramide B | *Leptolyngbya sp.* | Panama | Antiplasmodial activity ($IC_{50}$ = 570 nM) |
| Companeramide A | *Leptolyngbya sp.* | Panama | Antiplasmodial activity ($IC_{50}$ = 570 nM) |
| Ulongapeptin | *Lyngbya sp.* | Palau | Cytotoxic against KB cells ($IC_{50}$ = 0.63 μM) |
| Guineamide C | *Lyngbya majuscula* | Papua New Guinea | Cytotoxic against neuro-2a mouse neuroblastoma cells ($IC_{50}$ = 16 μM) |
| Boavistamide A | *Okeania sp.* | Cabo Verde | Moderate antiplasmodial activity ($EC_{50}$ = 2.21 μM); non-cytotoxic against HEK293T ($CC_{50}$ > 11.88 μM), HepG2 ($CC_{50}$ > 11.88 μM) and SF188 glioblastoma cells ($IC_{50}$ > 36 μM) |

**Table S11**. Potency and cytotoxicity profiles of boavistamides A–C (**1**–**3**) against *Plasmodium falciparum* Dd2 and two mammalian cell lines (HEK293T and HepG2). Selectivity index (SI) = $CC_{50}$ (mammalian cells) / $EC_{50}$ (*P. falciparum* Dd2). The assays were performed in one biological replicate that had two technical replicates.

| | **Selectivity Index compared to *P. falciparum* Dd2** | | ***P. falciparum* Dd2** | | | | **HEK293T** | | | **HepG2** | | |
|---|---|---|---|---|---|---|---|---|---|---|---|---|
| | **HEK293T** | **HepG2** | **$EC_{50}$ (µM)** | **$EC_{90}$ (µM)** | **Maximal Conc. (µM)** | **Signal at Max. Conc. (%)** | **$CC_{50}$ (µM)** | **Max Conc. (µM)** | **Signal at Max. Conc. (%)** | **$CC_{50}$ (µM)** | **Maximal Conc. (µM)** | **Signal at Max. Conc. (%)** |
| **Boavistamide A (1)** | >5.39 | >5.39 | 2.21 | 7.37 | 6.23 | -86.7 | >11.88 | 11.88 | -1.6 | >11.88 | 11.88 | -19.2 |
| **Boavistamide B (2)** | >5.29 | >5.29 | 2.25 | 21.72 | 6.23 | -77.1 | >11.88 | 11.88 | -0.4 | >11.88 | 11.88 | -19.5 |
| **Boavistamide C (3)** | 2.28 | >4.04 | 2.94 | 10.51 | 6.23 | -83.2 | 6.70 | 11.88 | -56.7 | >11.88 | 11.88 | -38.2 |

**Table S12**. *In vitro* antitrypanosomal activity of boavistamide A (**1**) against *Trypanosoma brucei brucei* (Lister 427).

| Compound | Concentration tested | % Inhibition | Conclusion |
|---|---|---|---|
| Boavistamide A (**1**) | 4 µM | −6 ± 10 | inactive |
| Pentamidine | 4 µM | 100 ± 1 | positive control |
| DMSO | 0.5–1 % | 0 ± 10 | negative control |

The assay was performed in technical quadruplicate.

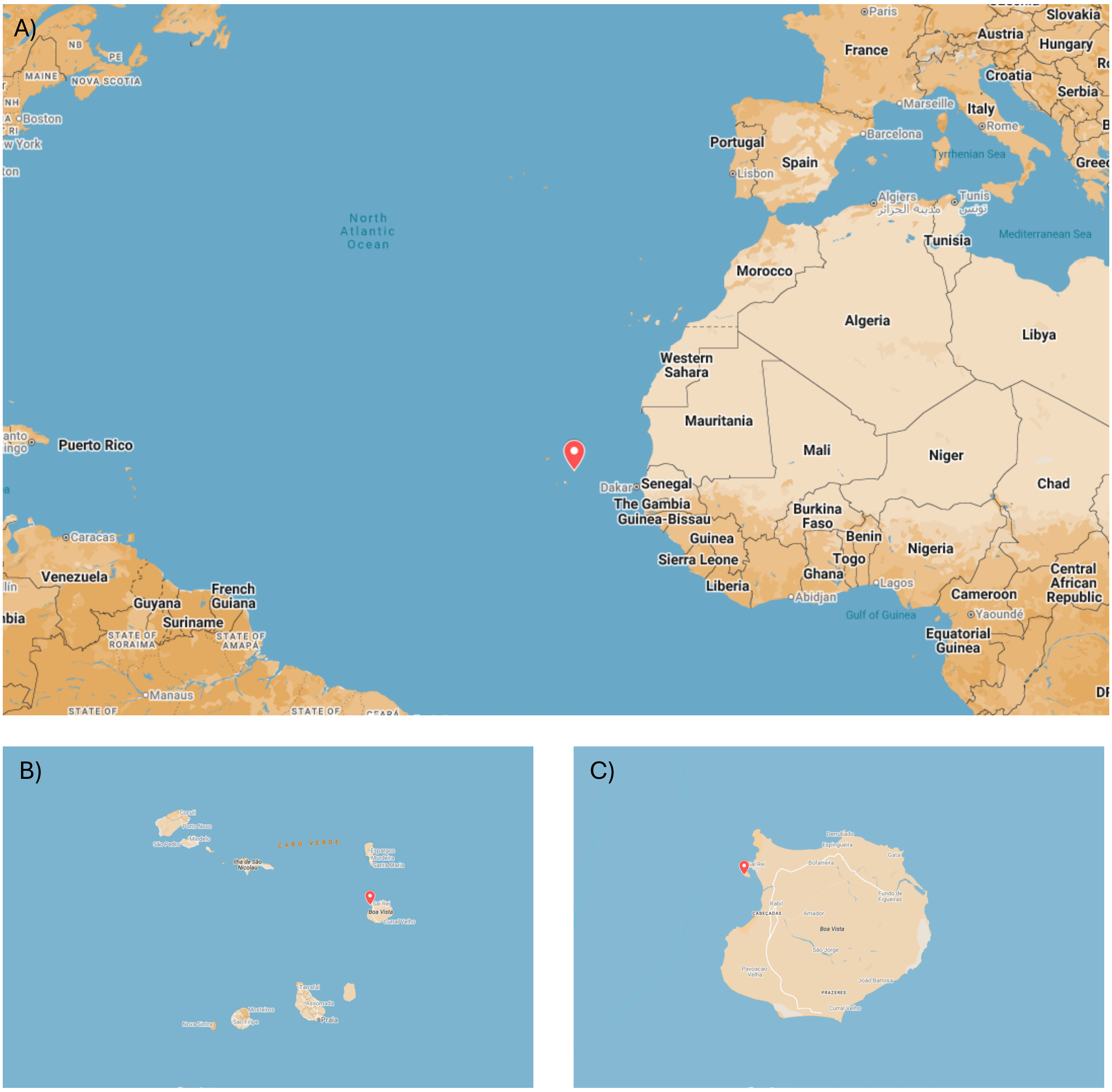

**Figure S1.** Geographic origin of environmental cyanobacterial sample collected in Insular West Africa. (A) Overview map showing collection site in Cabo Verde. (B) General map of the Cabo Verde archipelago. (C) Zoom-in on Boa Vista Island (Cabo Verde).

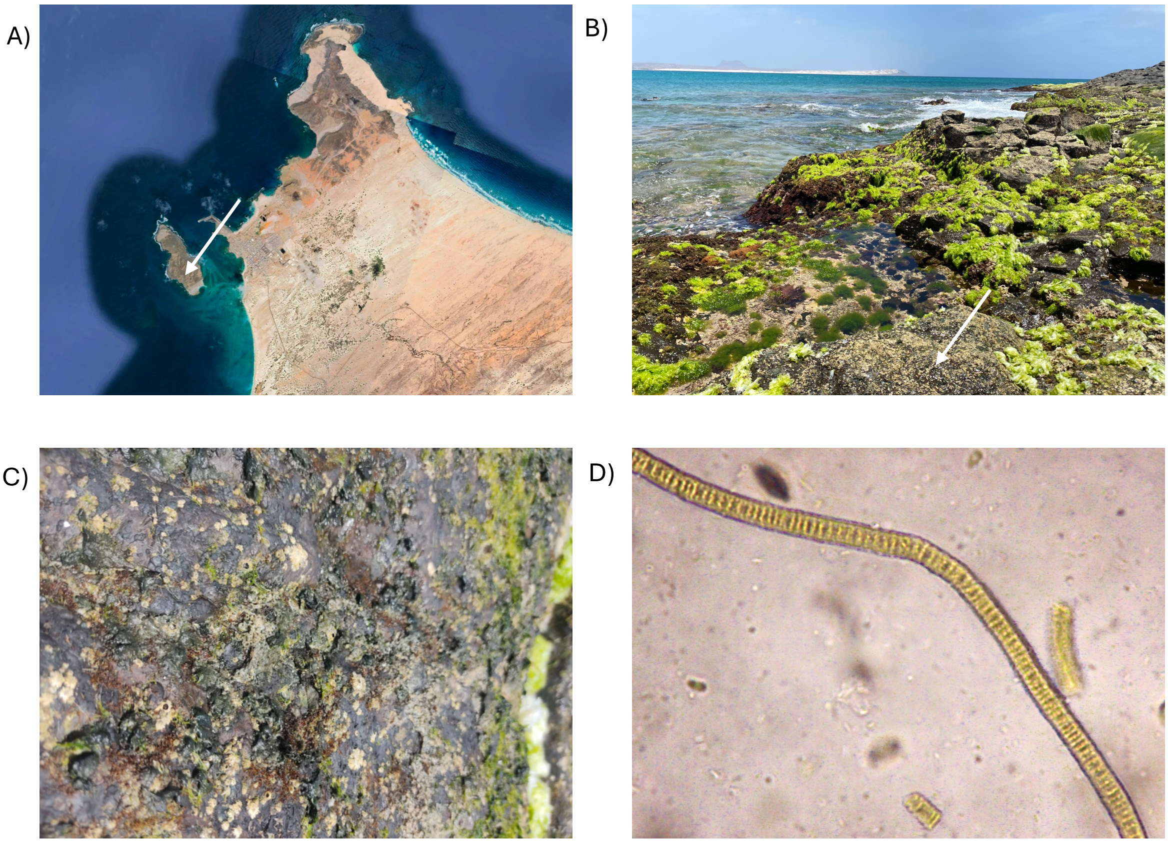

**Figure S2.** Collection site and morphological observations of sample 2370 from Ilhéu de Sal Rei (Boa Vista, Cabo Verde). (A) Satellite view showing the collection location at Ilhéu de Sal Rei. (B) Field photograph of the sampling area on the islet. (C) Close-up image of the cyanobacterial biofilm collected from the rock surface. (D) Microscopic image of the RNAlater-preserved material showing filamentous cyanobacteria with morphology consistent with *Okeania*-like taxa, representing the predominant morphotype observed.

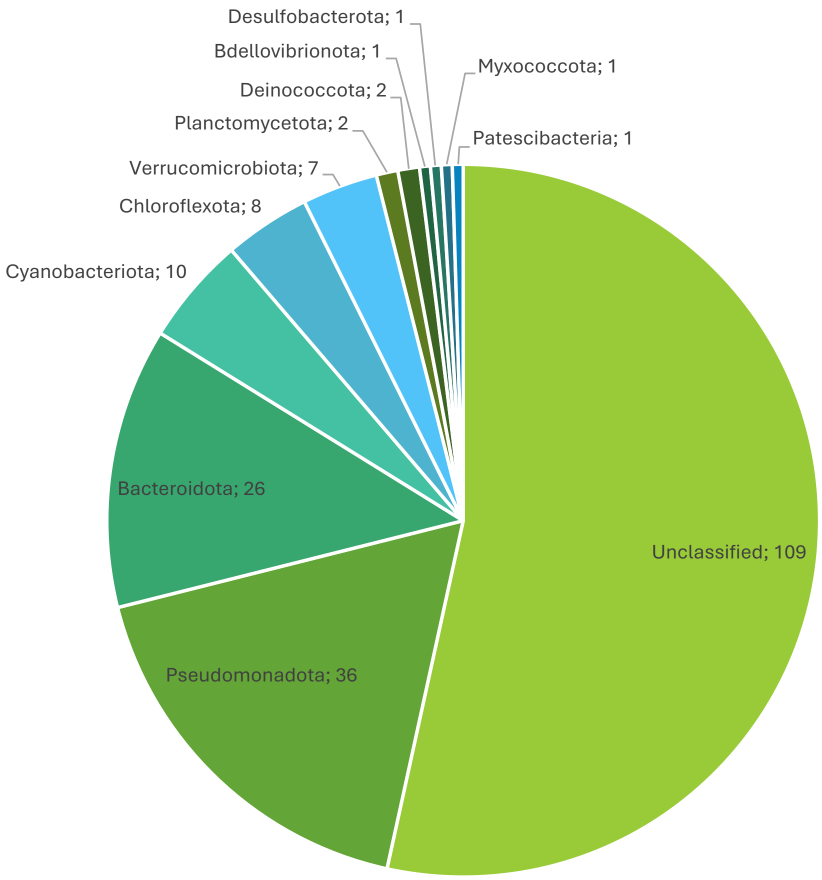

**Figure S3.** Taxonomic distribution of metagenomic bins (phylum level) in sample 2370 from Ilhéu de Sal Rei (Boa Vista, Cabo Verde), based on GTDB-Tk classification.

A)

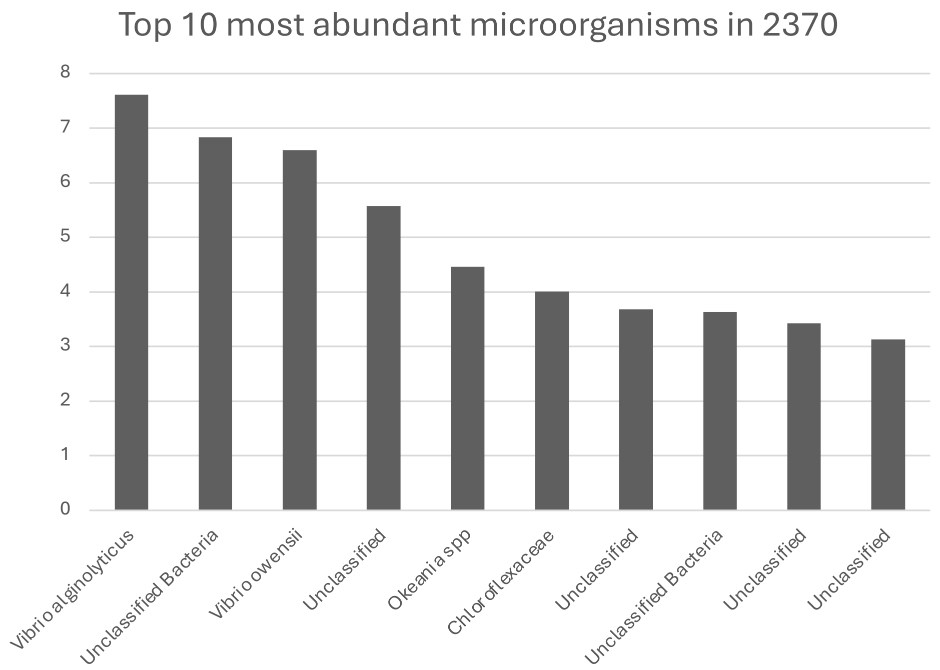

B)

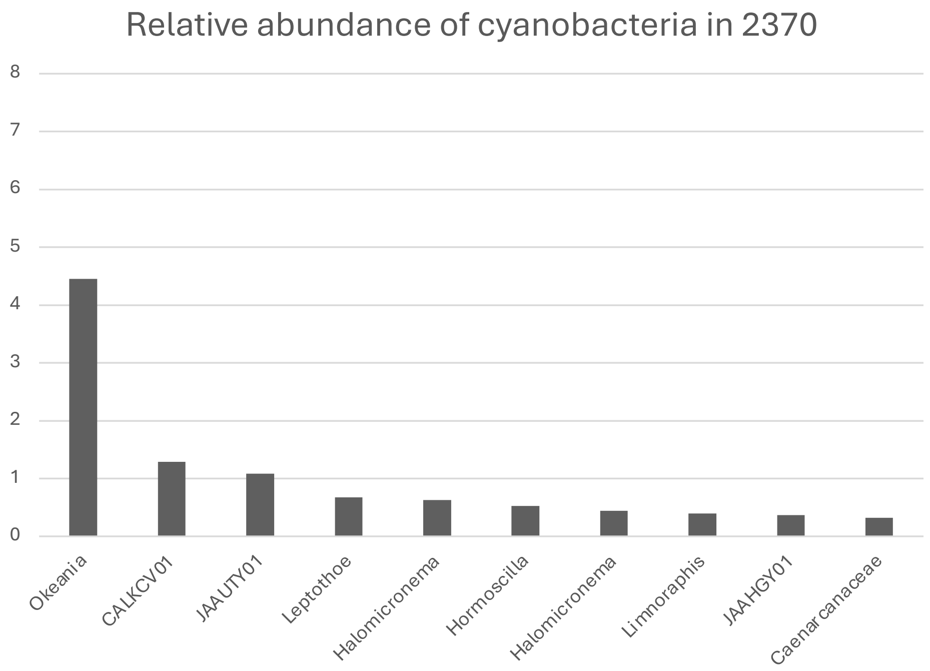

**Figure S4**. Anvi'o analysis. A) Top 10 most abundant microorganisms in 2370, B) Relative abundance of cyanobacteria in sample 2370 from Ilhéu de Sal Rei (Boa Vista, Cabo Verde).

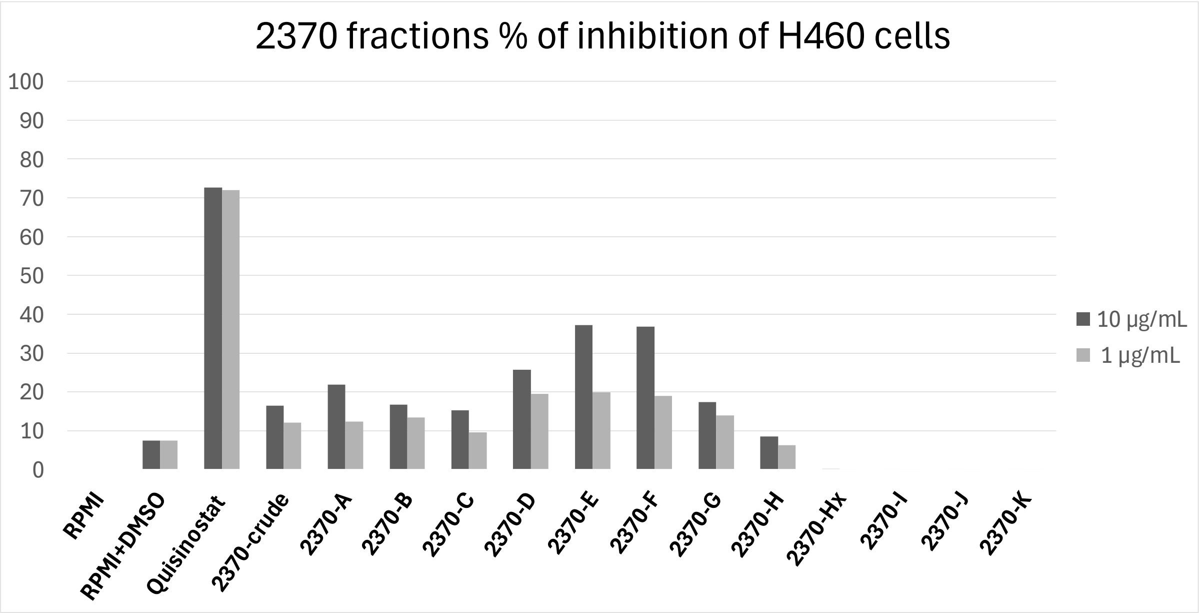

**Figure S5.** Percentage inhibition of H-460 human non-small cell lung carcinoma cell proliferation by VLC fractions (A–K) of the organic extract from environmental cyanobacterial sample 2370.

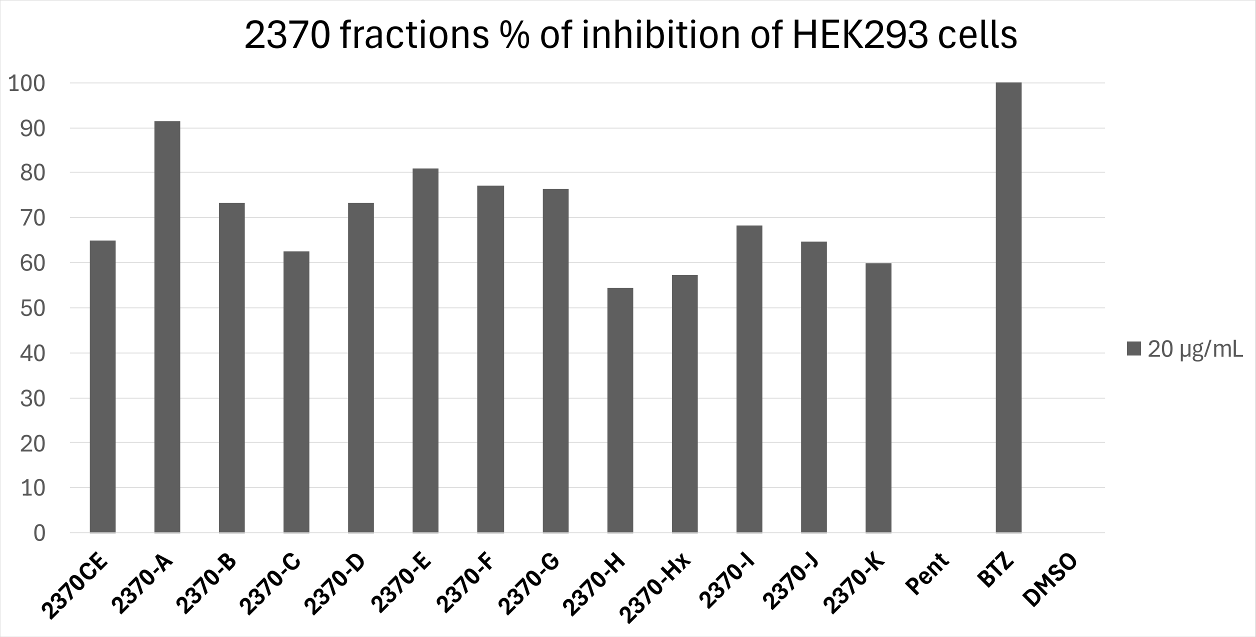

**Figure S6.** Percentage inhibition of HEK293 cell proliferation by VLC fractions (A–K) of the organic extract from environmental cyanobacterial sample 2370. The controls, pentamidine (Pent) and bortezomib (BTZ) were tested at 4 and 1 µM, respectively. The assay was performed in technical quadruplicate.

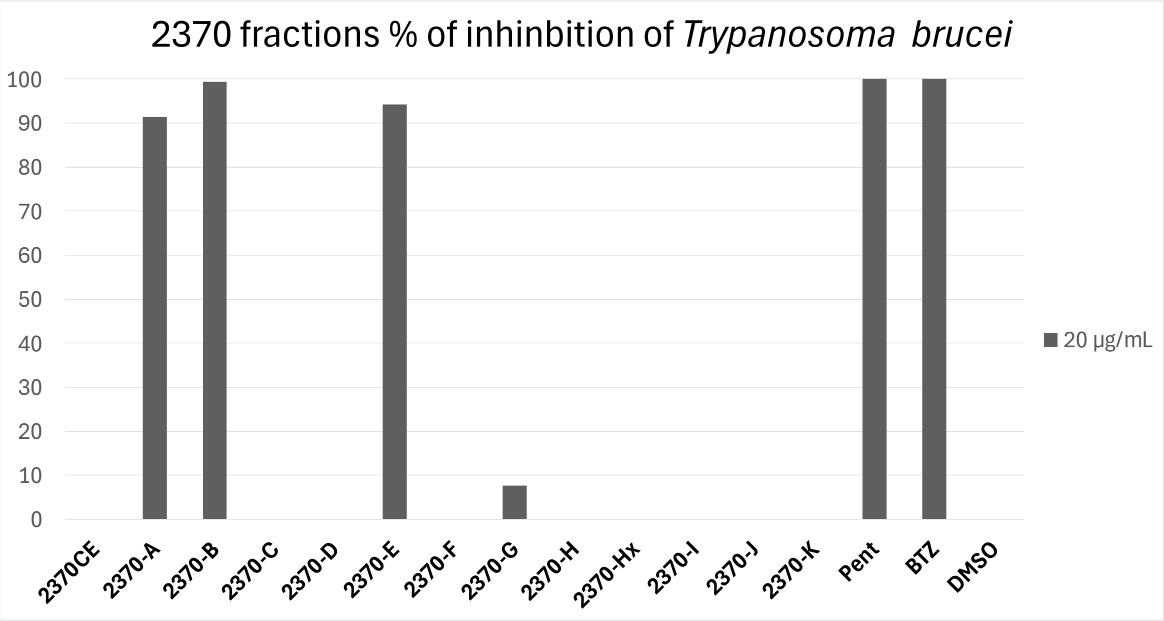

**Figure S7.** Percentage inhibition of *Trypanosoma brucei* proliferation by VLC fractions (A–K) obtained from the organic extract of environmental cyanobacterial sample 2370. The controls, pentamidine (Pent) and bortezomib (BTZ) were tested at 4 and 1 µM, respectively. The assay was performed in technical quadruplicate.

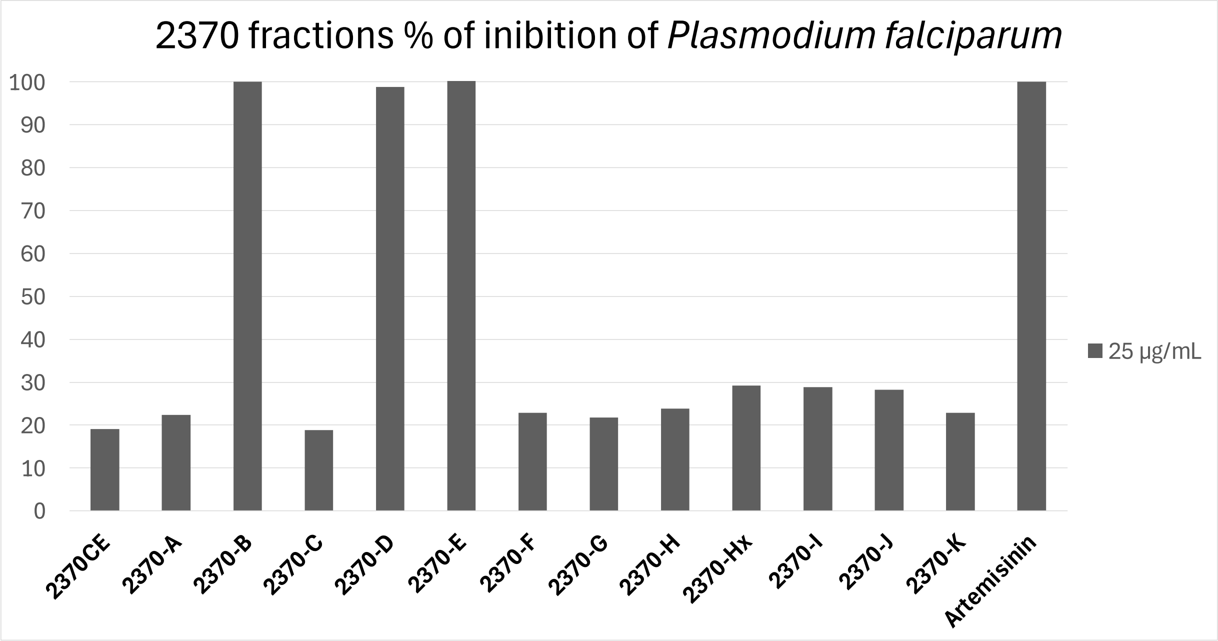

**Figure S8.** Percentage inhibition of *Plasmodium falciparum* proliferation by VLC fractions (A–K) from the organic extract of environmental cyanobacterial sample 2370. Artemisinin was used as the positive control. The assay was performed in technical quintuplicate.

MS1 boavistamide A **(1)**

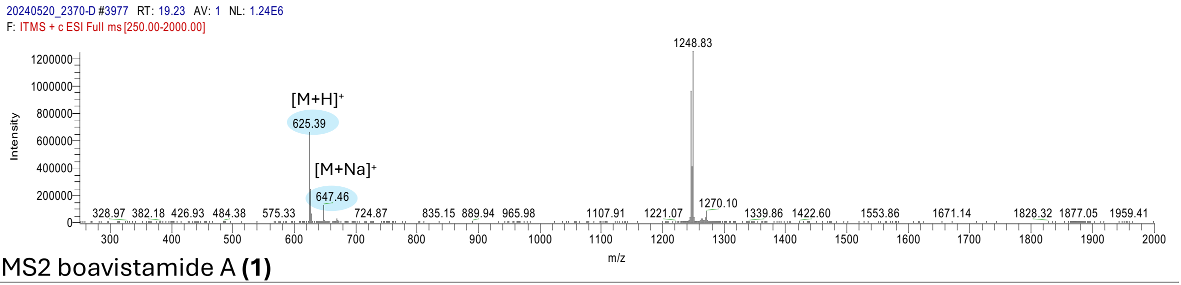

MS2 boavistamide A **(1)**

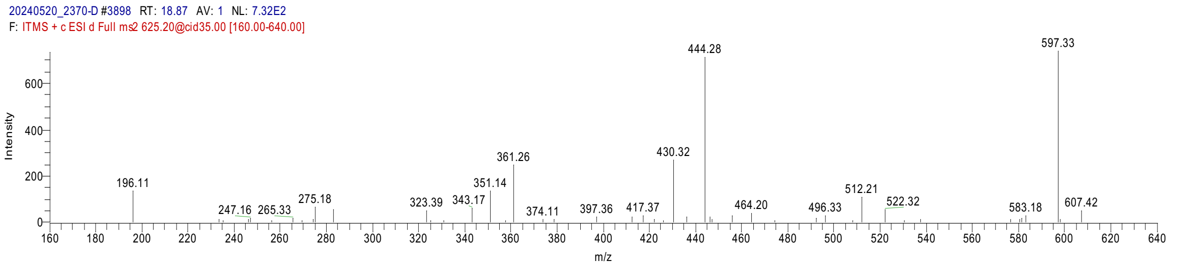

**Figure S9.** LC-MS1 and MS2 spectra of boavistamide A (**1**), showing the $[M+H]^+$ and $[M+Na]^+$ ions and corresponding fragmentation patterns.

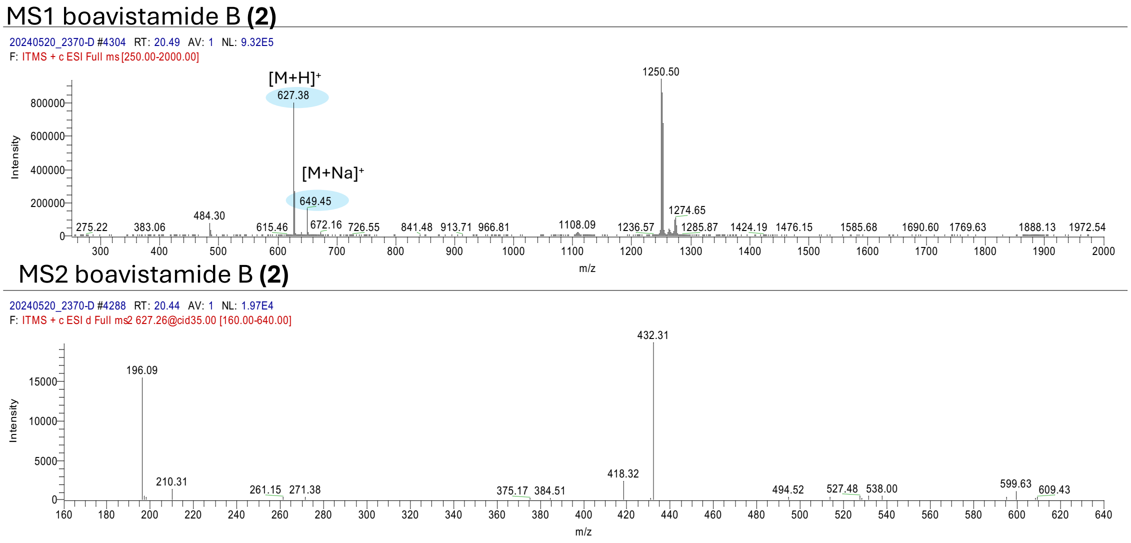

**Figure S10.** LC-MS1 and MS2 spectra of boavistamide B (**2**), showing the $[M+H]^+$ and $[M+Na]^+$ ions and corresponding fragmentation patterns.

MS1 Boavistamide C **(3)**

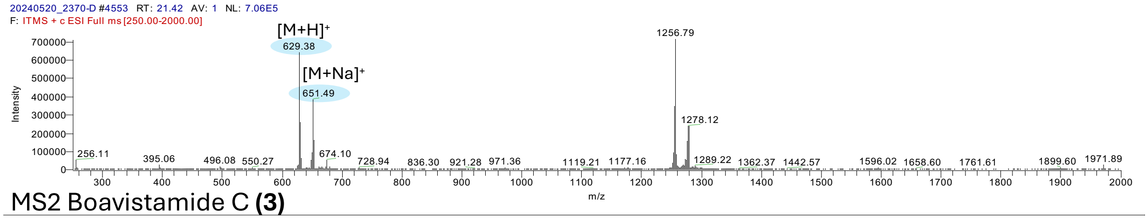

MS2 Boavistamide C **(3)**

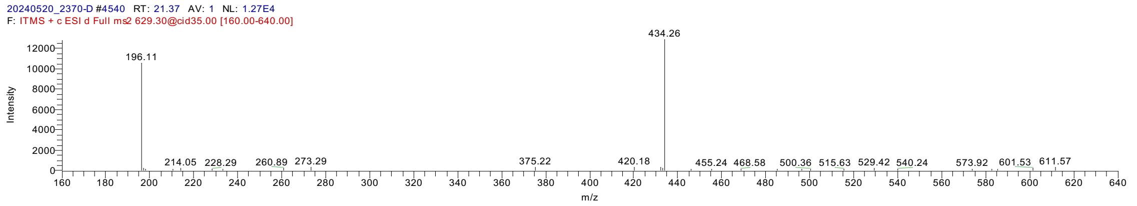

**Figure S11.** LC-MS1 and MS2 spectra of boavistamide C (**3**), showing the $[M+H]^+$ and $[M+Na]^+$ ions and corresponding fragmentation patterns.

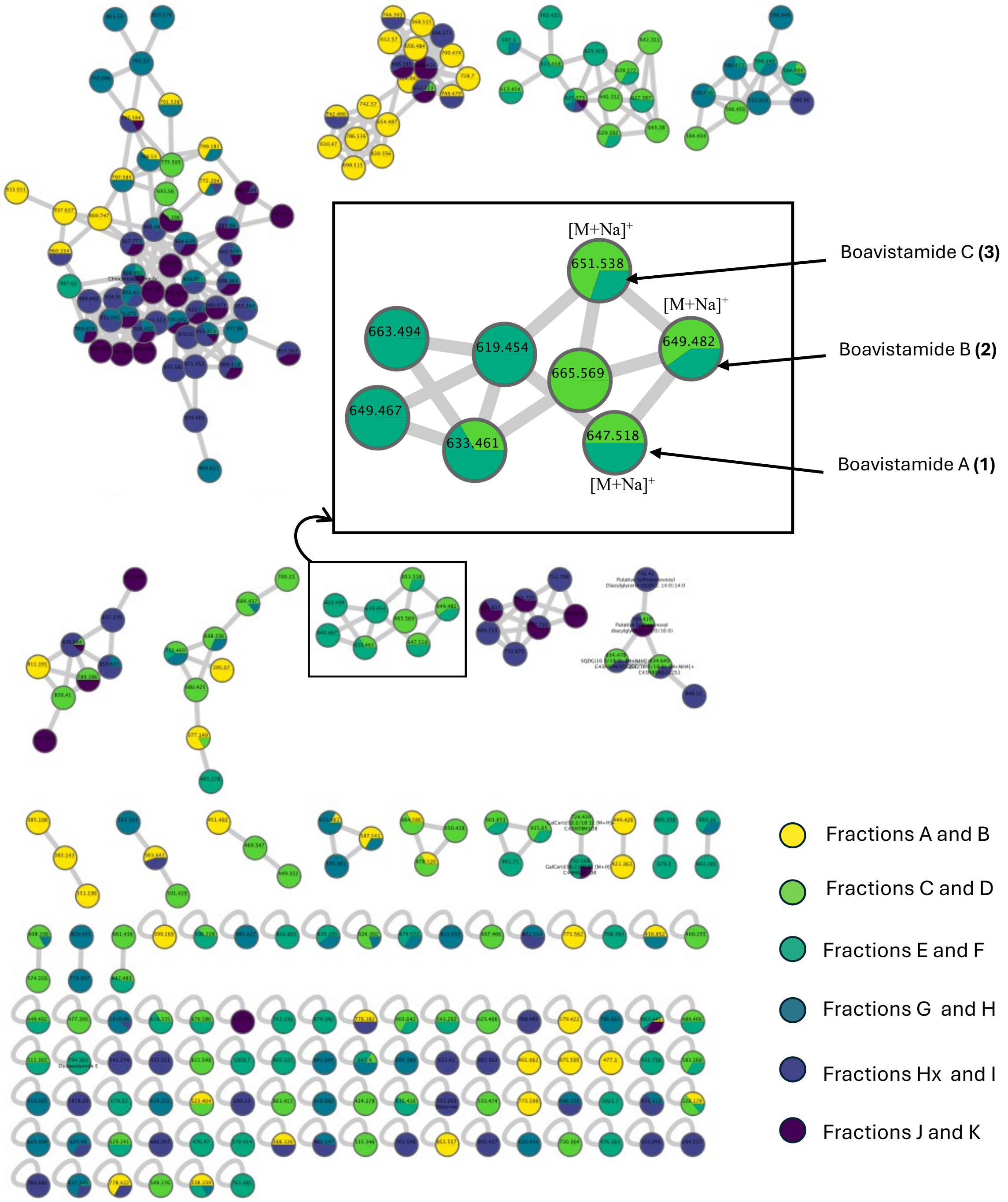

**Figure S12.** MS/MS molecular network derived from VLC fractions of cyanobacterial extract 2370 (Ilhéu de Sal Rei, Boa Vista Island, Cabo Verde), generated using the GNPS platform and visualized in Cytoscape. A unique and chemically unprecedented cluster corresponding to boavistamides A–C (**1**–**3**) is highlighted. This molecular family, absent from GNPS spectral libraries and the MarinLit database, underscores the novelty of these compounds. Node colors indicate the fraction of origin. Edges represent MS/MS spectral similarity (cosine score > 0.7).

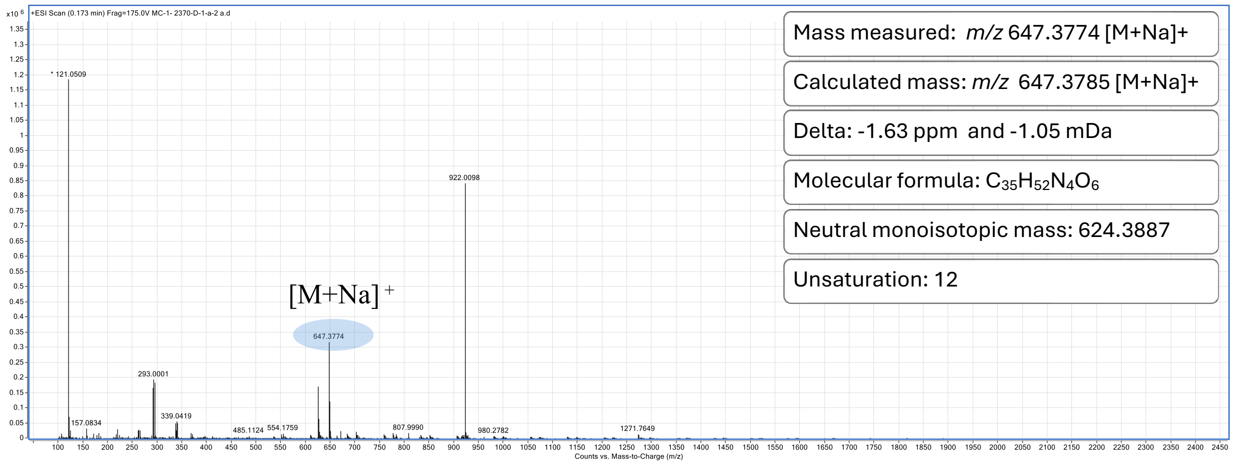

**Figure S13.** HRESI(+)MS spectrum of boavistamide A (**1**).

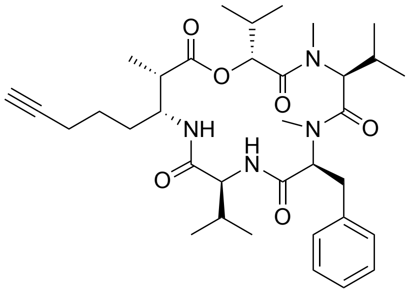

Boavistamide A (**1**)

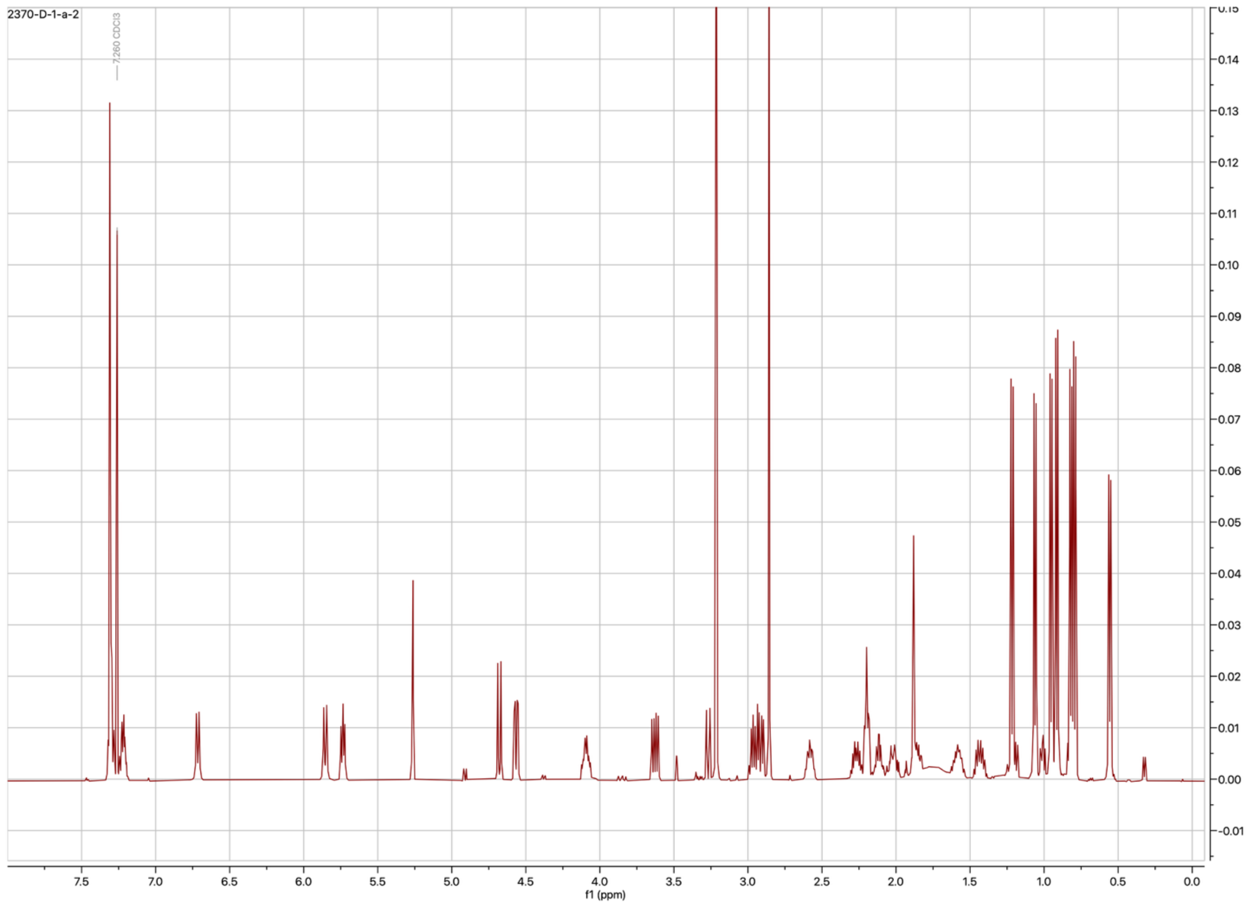

**Figure S14.** $^{1}$H NMR spectrum of boavistamide A (**1**) in $CDCl_3$ (500 MHz).

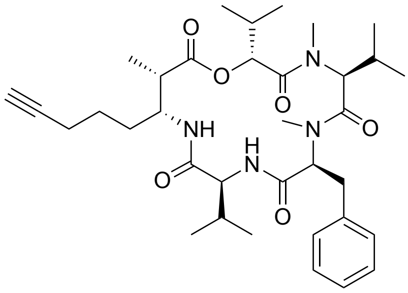

Boavistamide A (**1**)

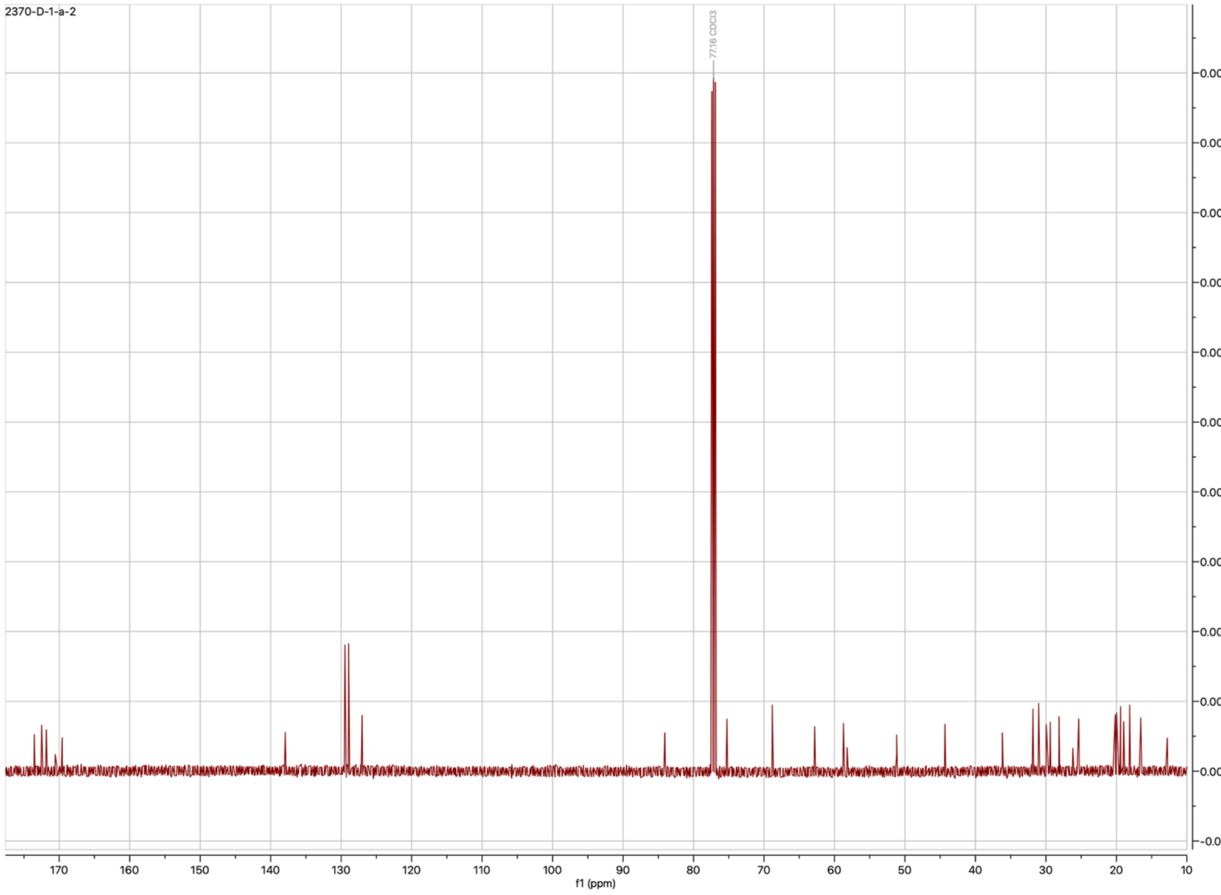

**Figure S15.** $^{13}$C NMR spectrum of boavistamide A (**1**) in $CDCl_3$ (125 MHz).

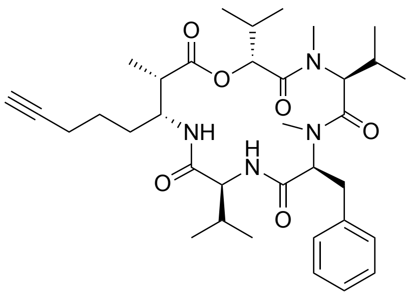

Boavistamide A (**1**)

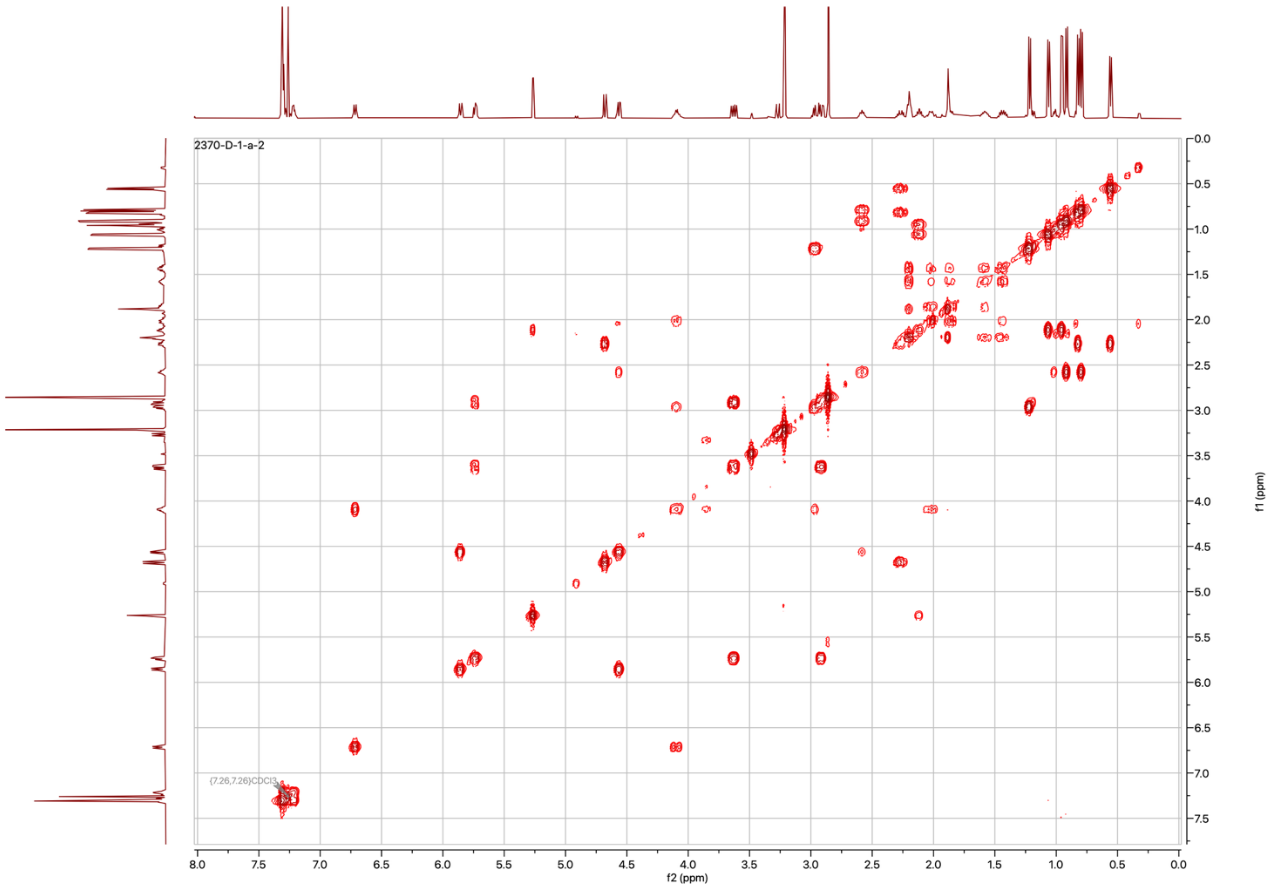

**Figure S16.** $^{1}$H-$^{1}$H COSY spectrum of boavistamide A (**1**) in $CDCl_3$ (500 MHz).

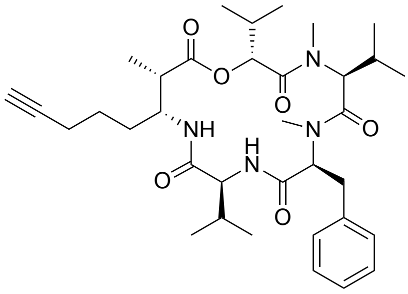

Boavistamide A (**1**)

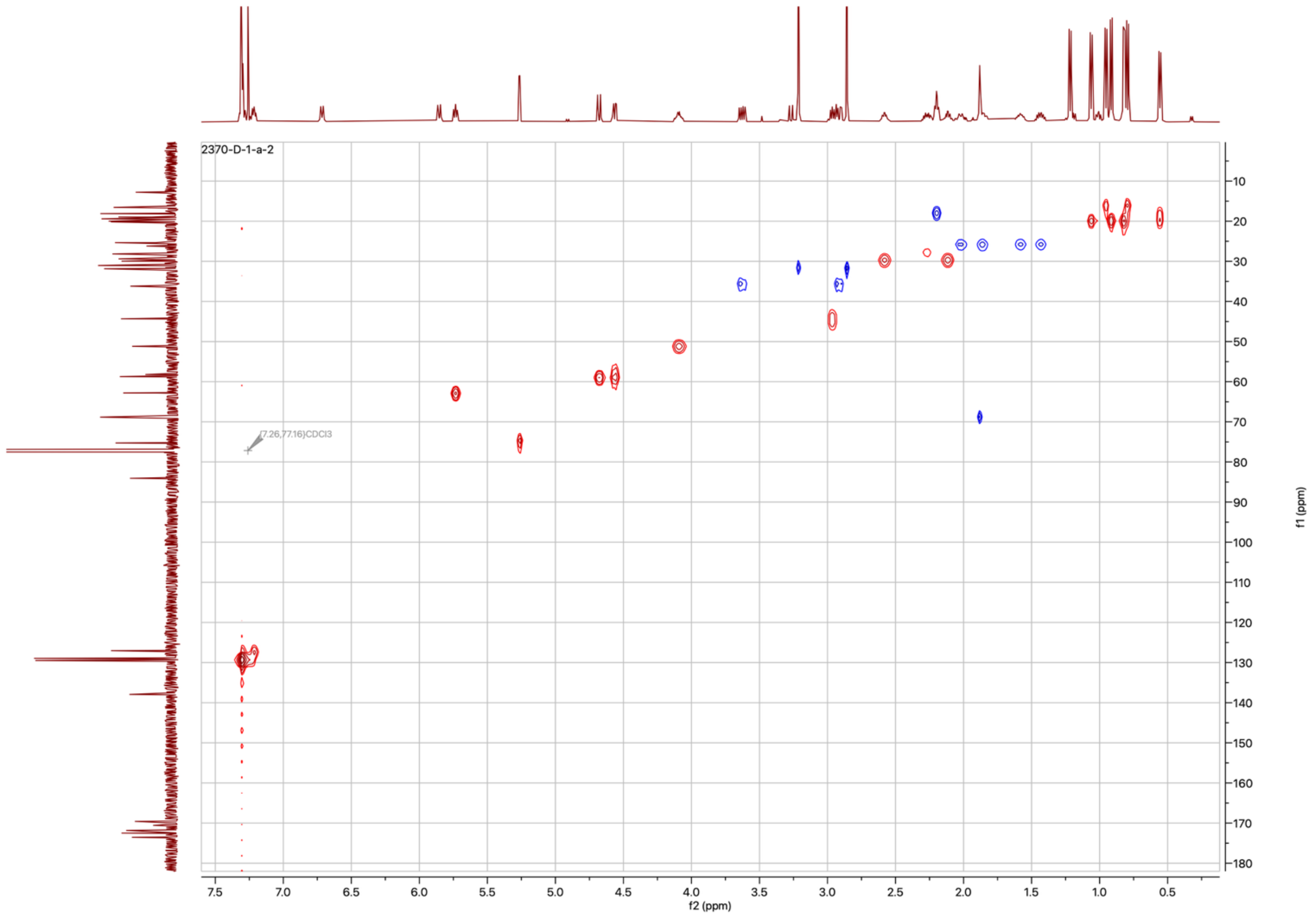

**Figure S17.** Multiplicity-edited HSQC spectrum of boavistamide A (**1**) in $CDCl_3$ (500 MHz).

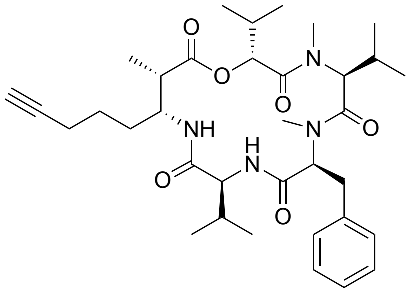

Boavistamide A **(1)**

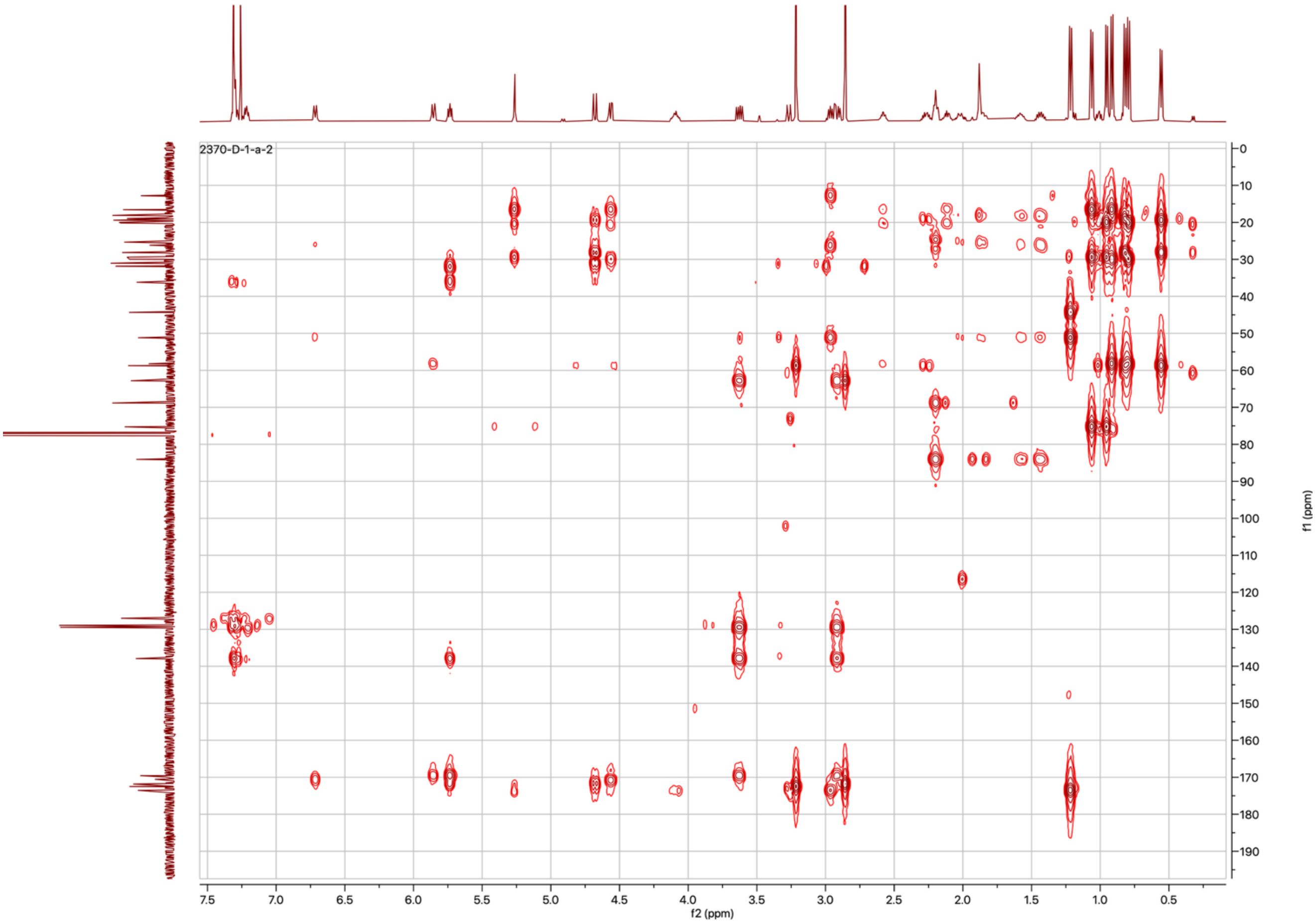

**Figure S18.** HMBC spectrum of boavistamide A (**1**) in $CDCl_3$ (500 MHz).

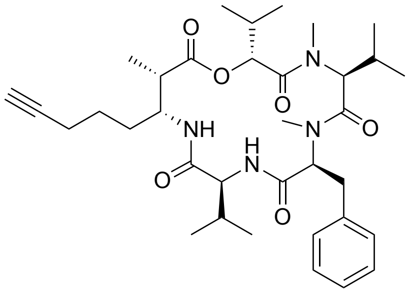

Boavistamide A **(1)**

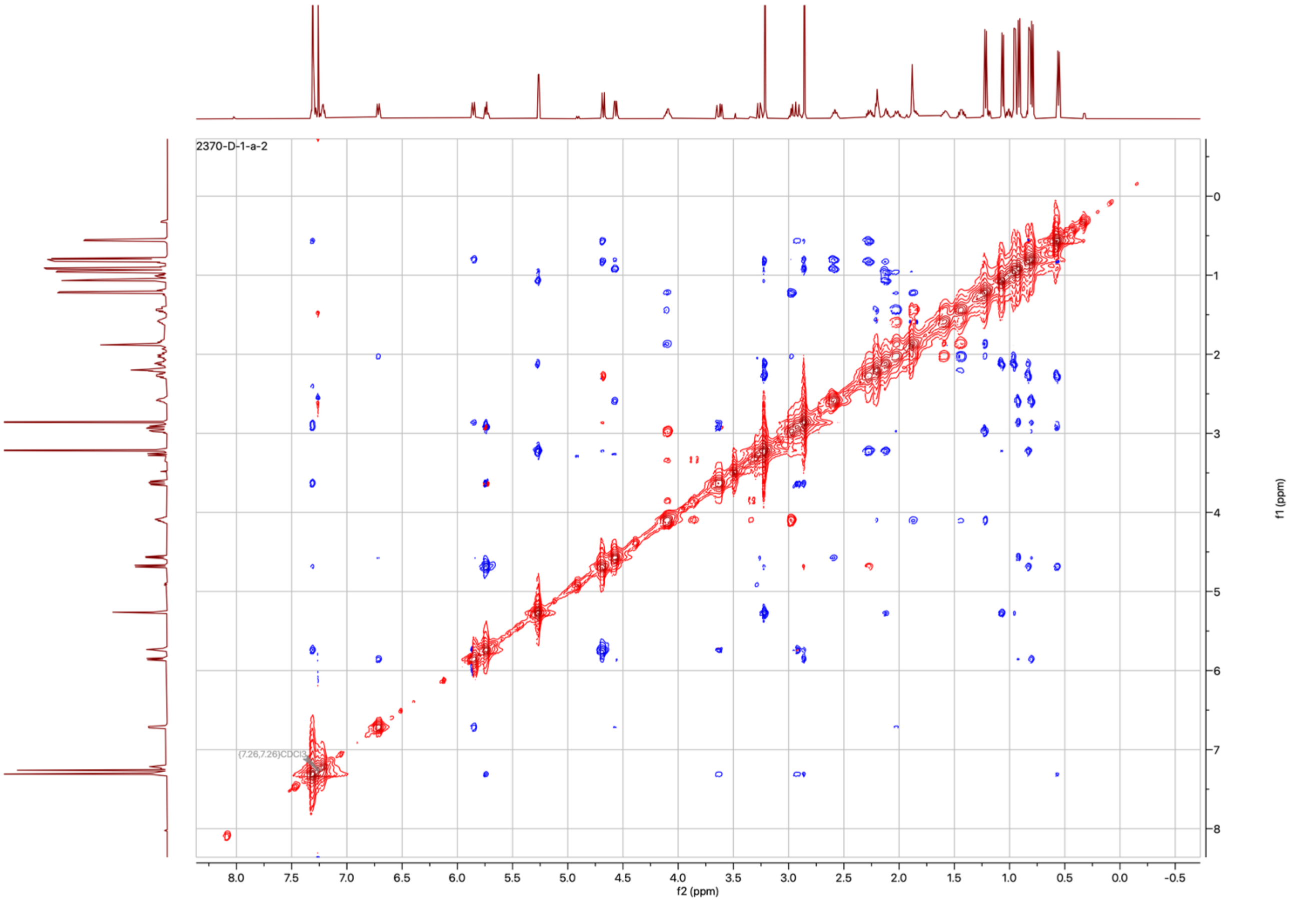

**Figure S19.** ROESY spectrum of boavistamide A (**1**) in $CDCl_3$ (500 MHz).

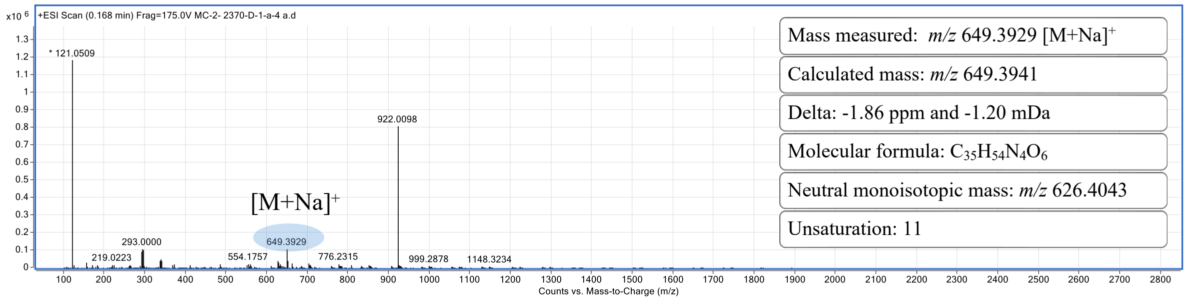

**Figure S20.** HRESI(+)MS spectrum of boavistamide B (**2**).

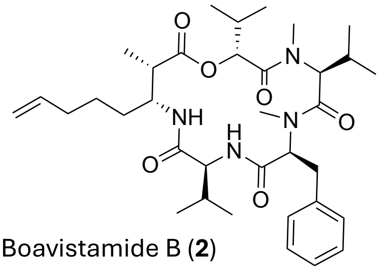

Boavistamide B (**2**)

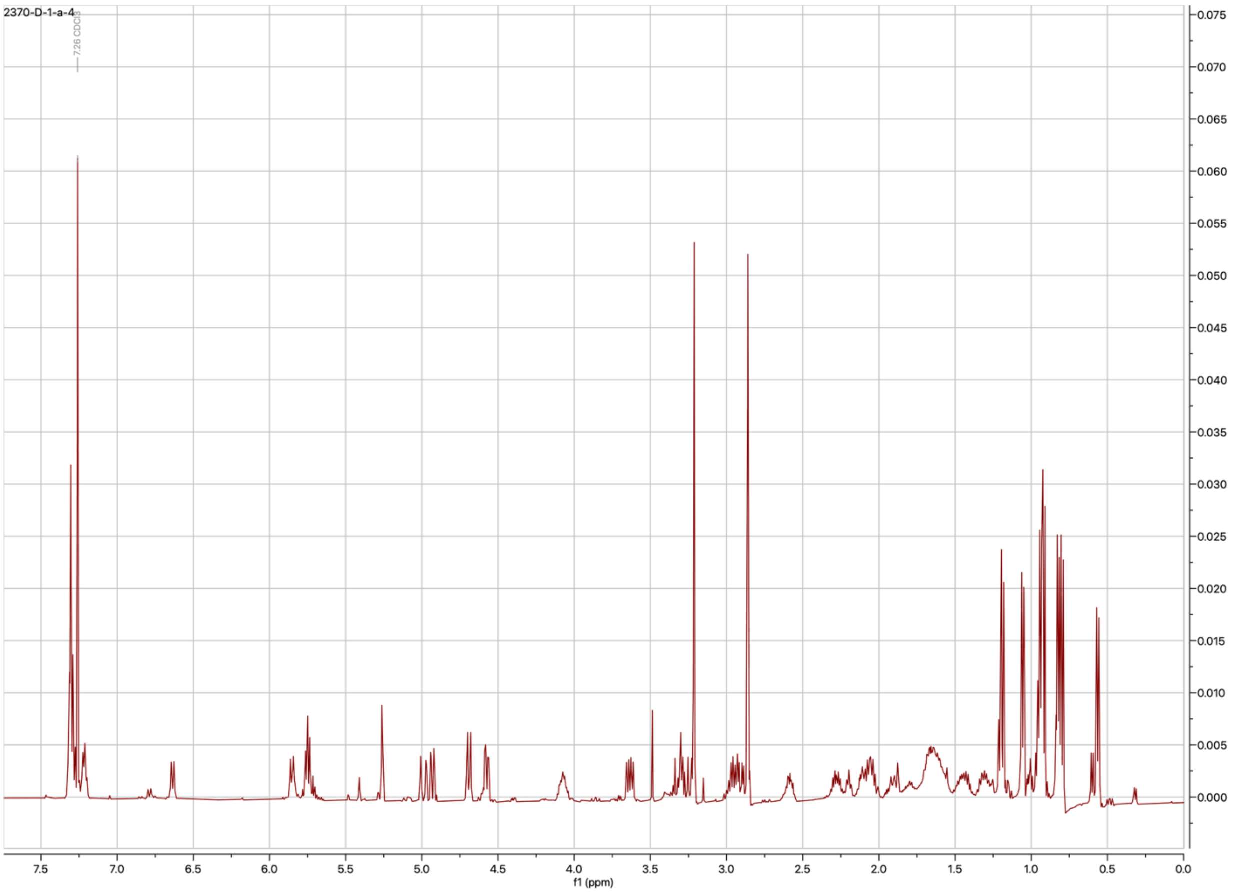

**Figure S21.** $^1$H NMR spectrum of boavistamide B (**2**) in $CDCl_3$ (500 MHz).

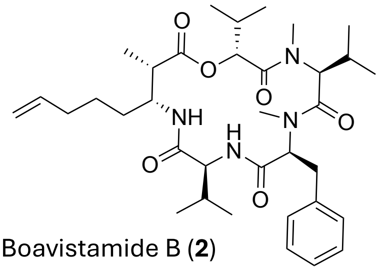

Boavistamide B (**2**)

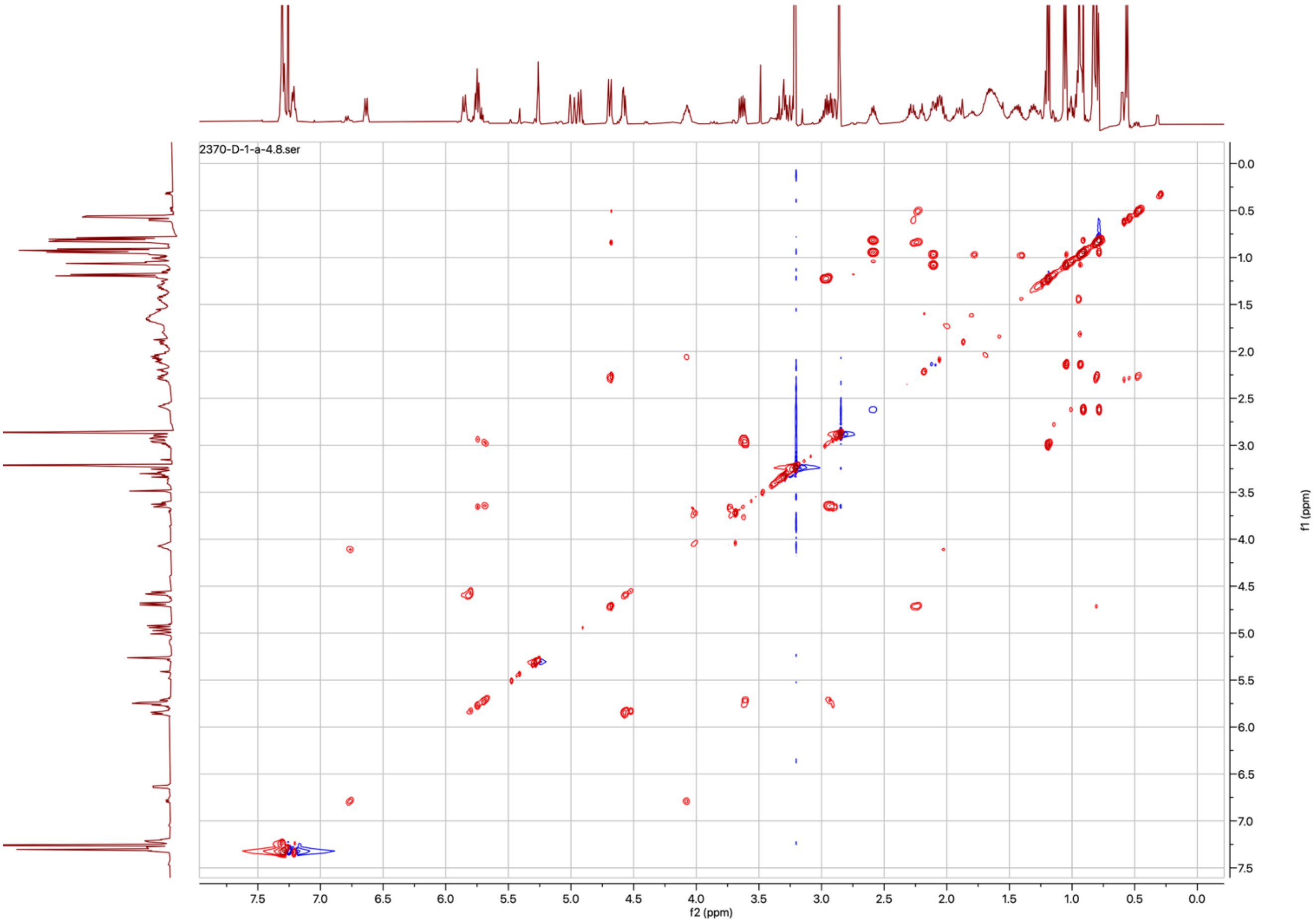

**Figure S22**. $^1$H-$^1$H COSY spectrum of boavistamide B (**2**) in $CDCl_3$ (600 MHz).

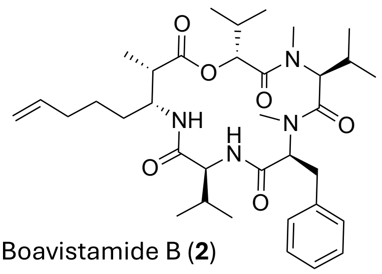

Boavistamide B (**2**)

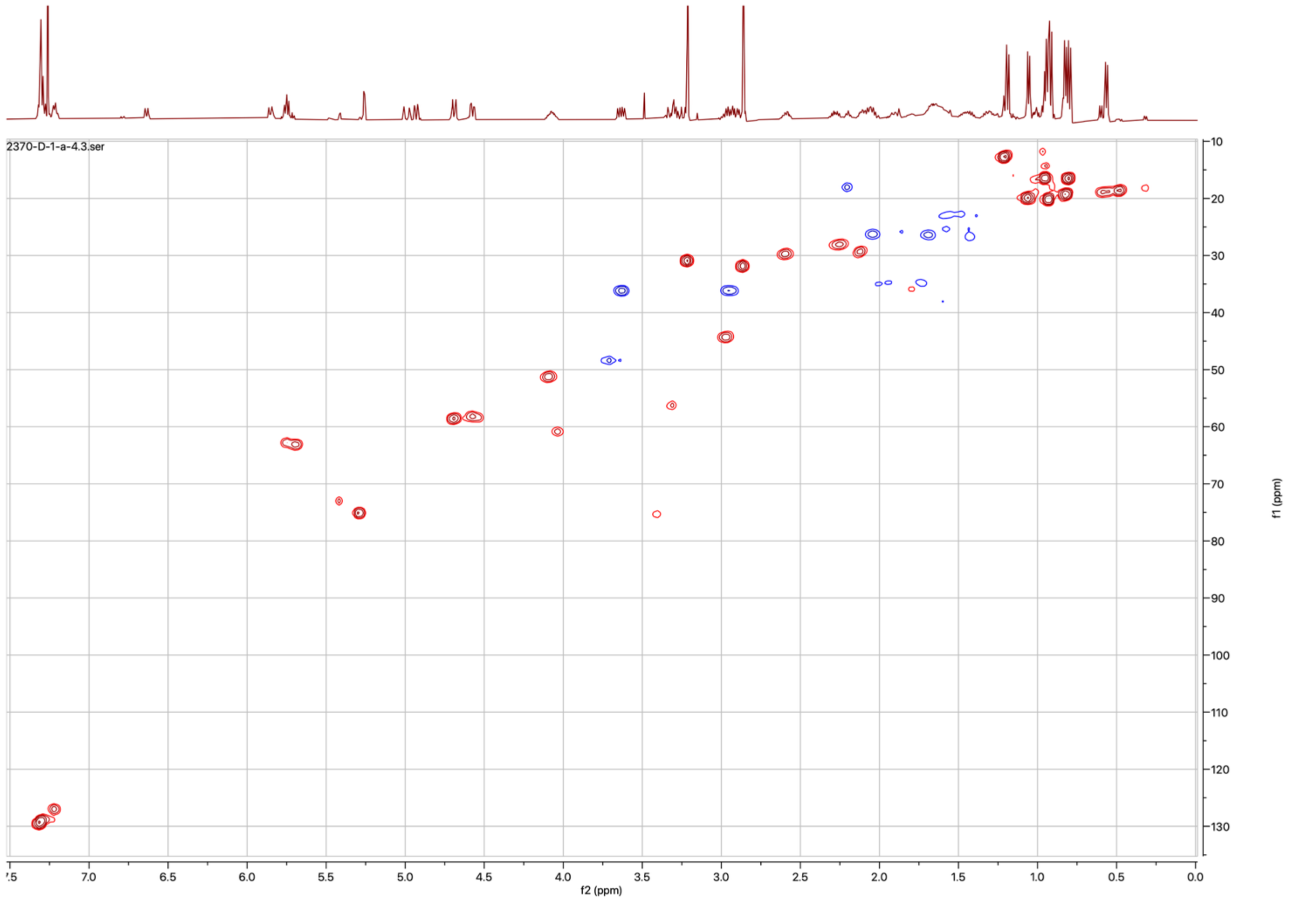

**Figure S23.** Multiplicity-edited HSQC spectrum of boavistamide B (**2**) in $CDCl_3$ (600 MHz).

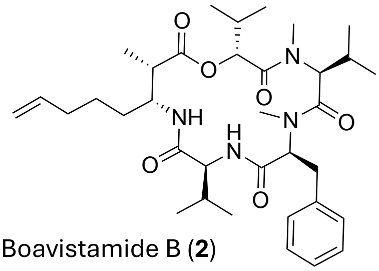

Boavistamide B (**2**)

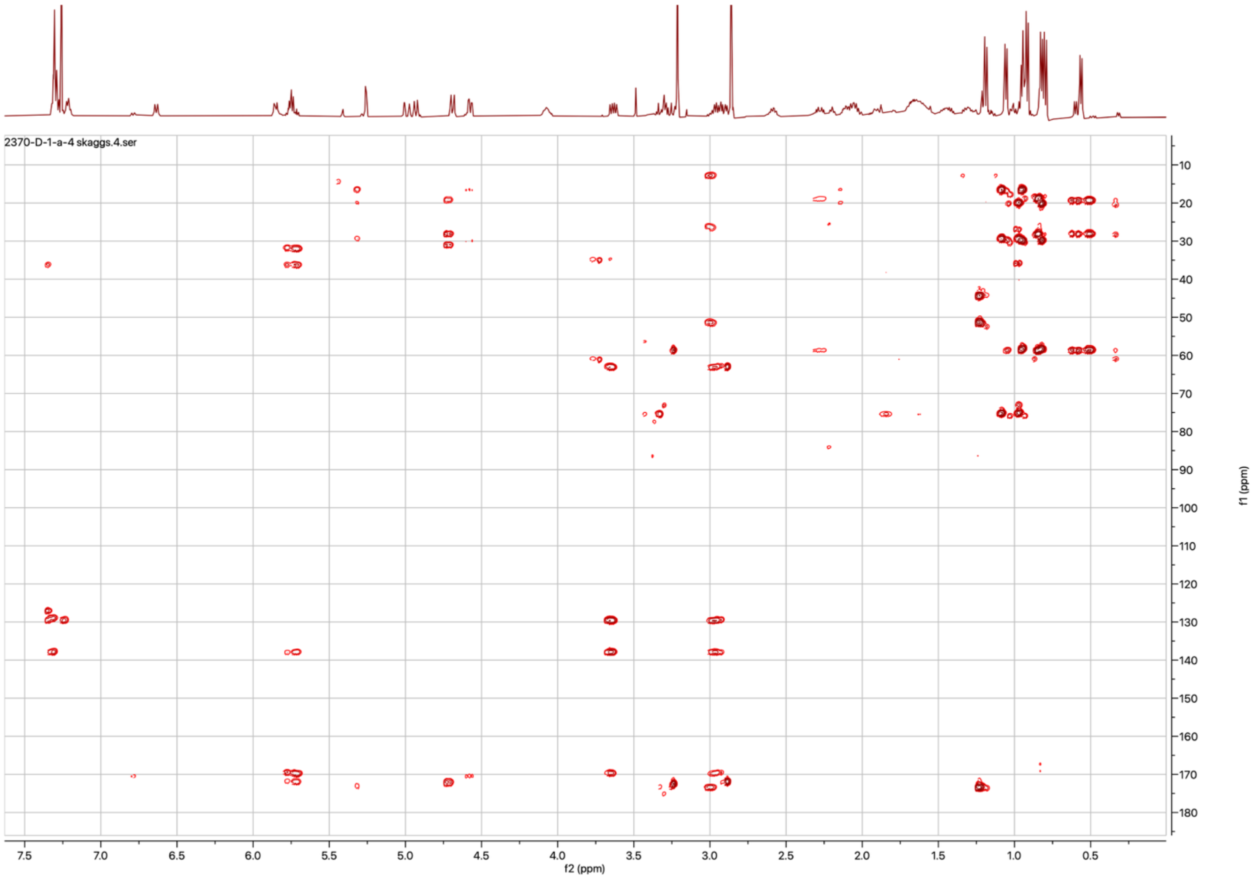

**Figure S24.** HMBC spectrum of boavistamide B (**2**) in $CDCl_3$ (600 MHz).

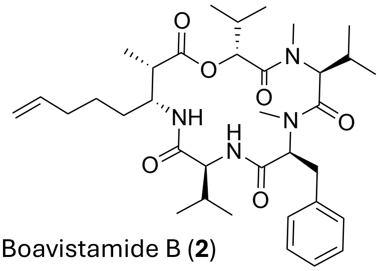

Boavistamide B (**2**)

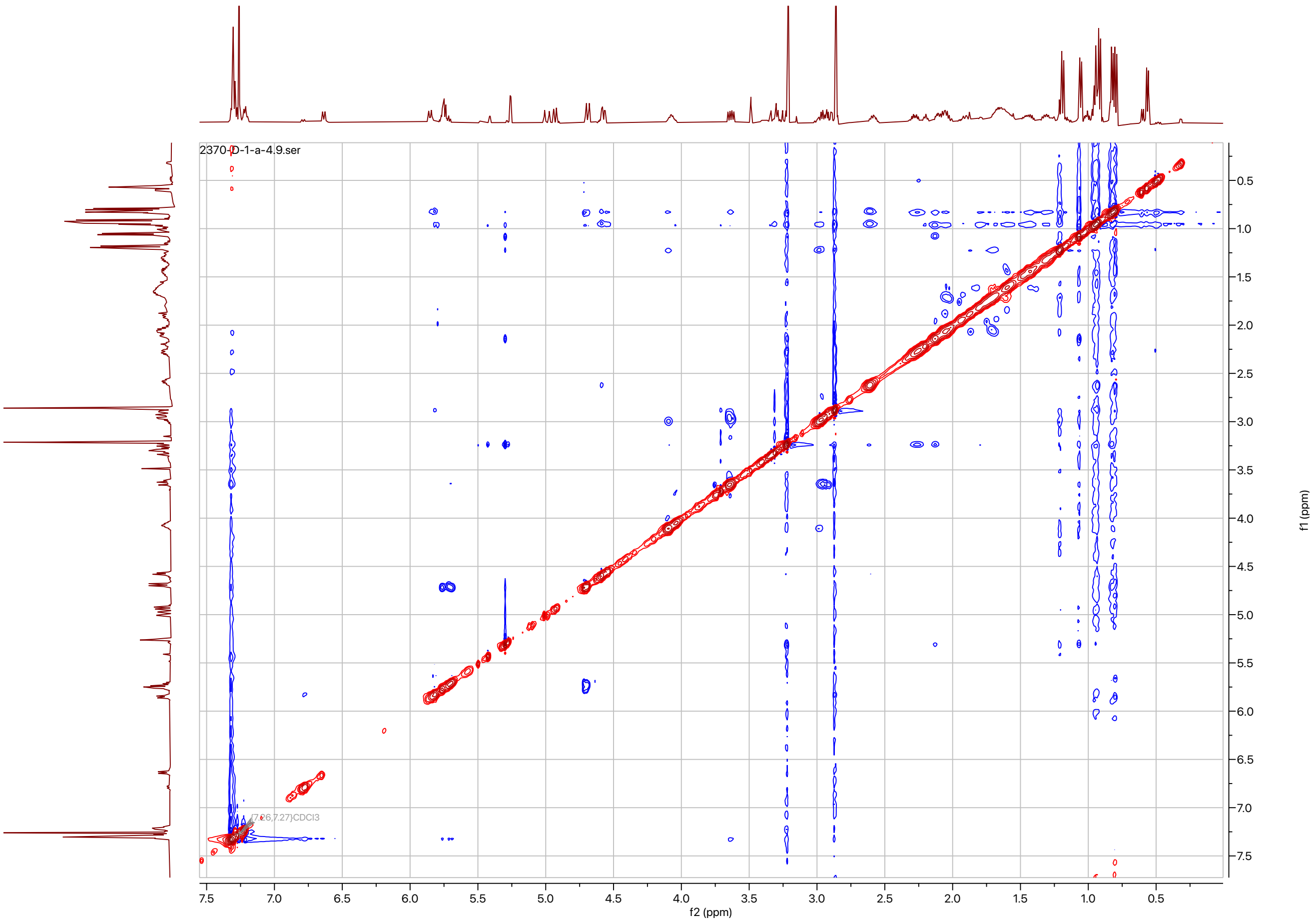

**Figure S25.** ROESY spectrum of boavistamide B (**2**) in $CDCl_3$ (600 MHz).

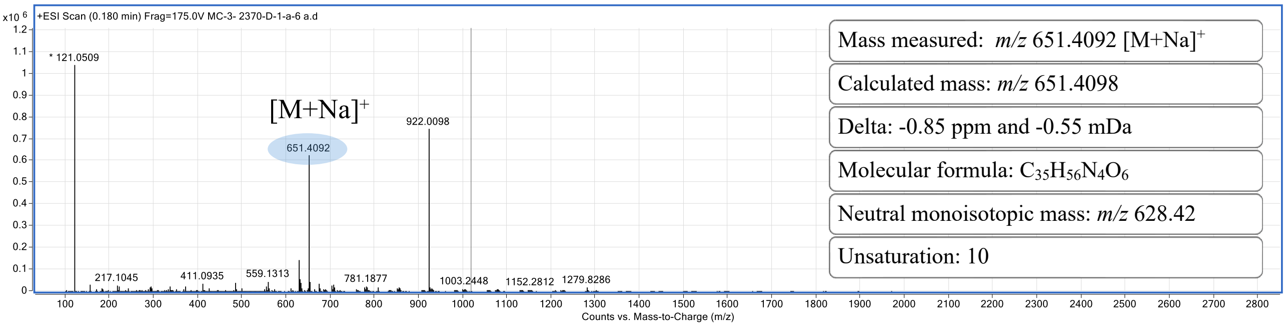

**Figure S26.** HRESI(+)MS spectrum of boavistamide C (**3**).

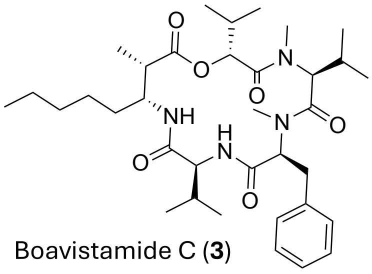

Boavistamide C (**3**)

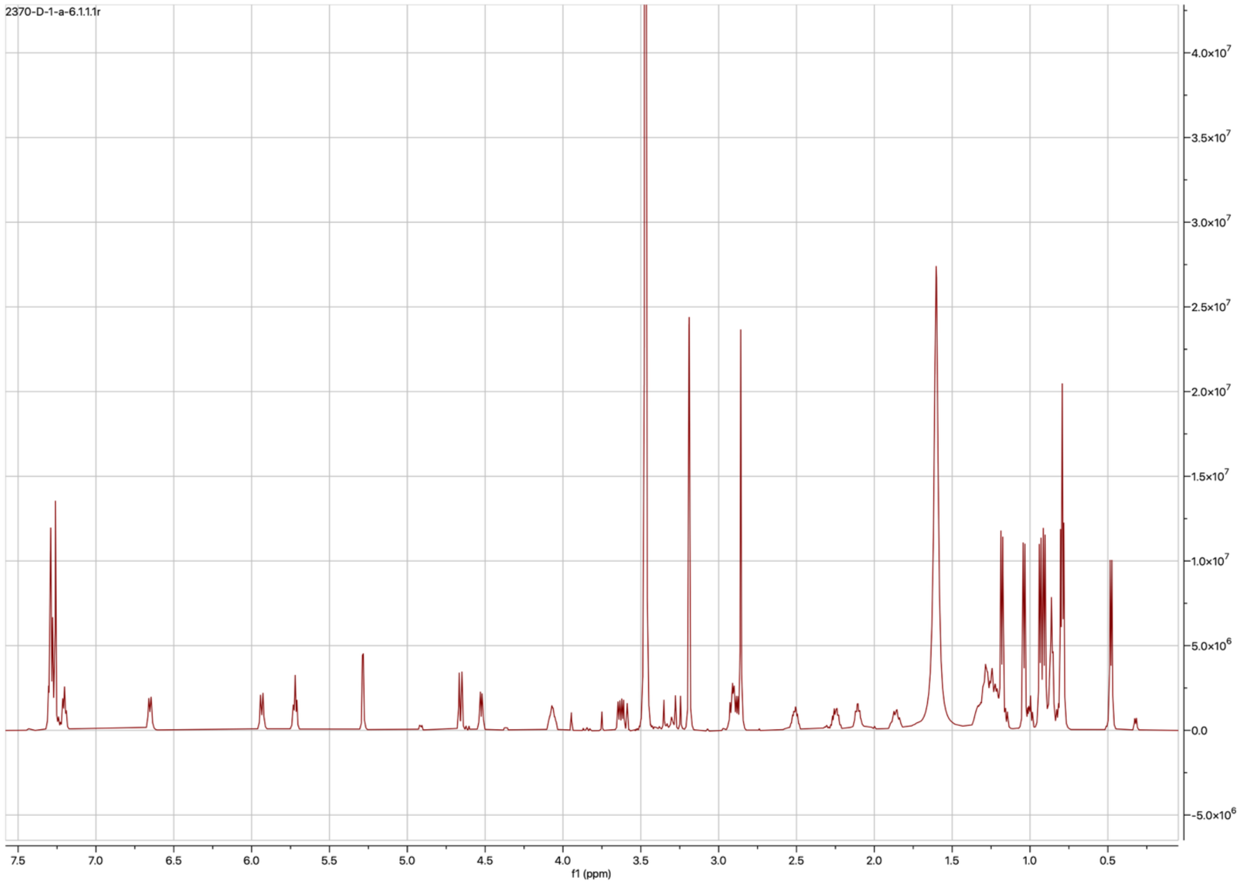

**Figure S27.** $^1$H NMR spectrum of boavistamide C (**3**) in $CDCl_3$ (600 MHz).

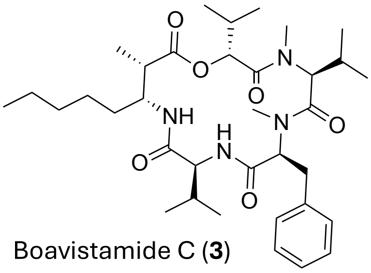

Boavistamide C (**3**)

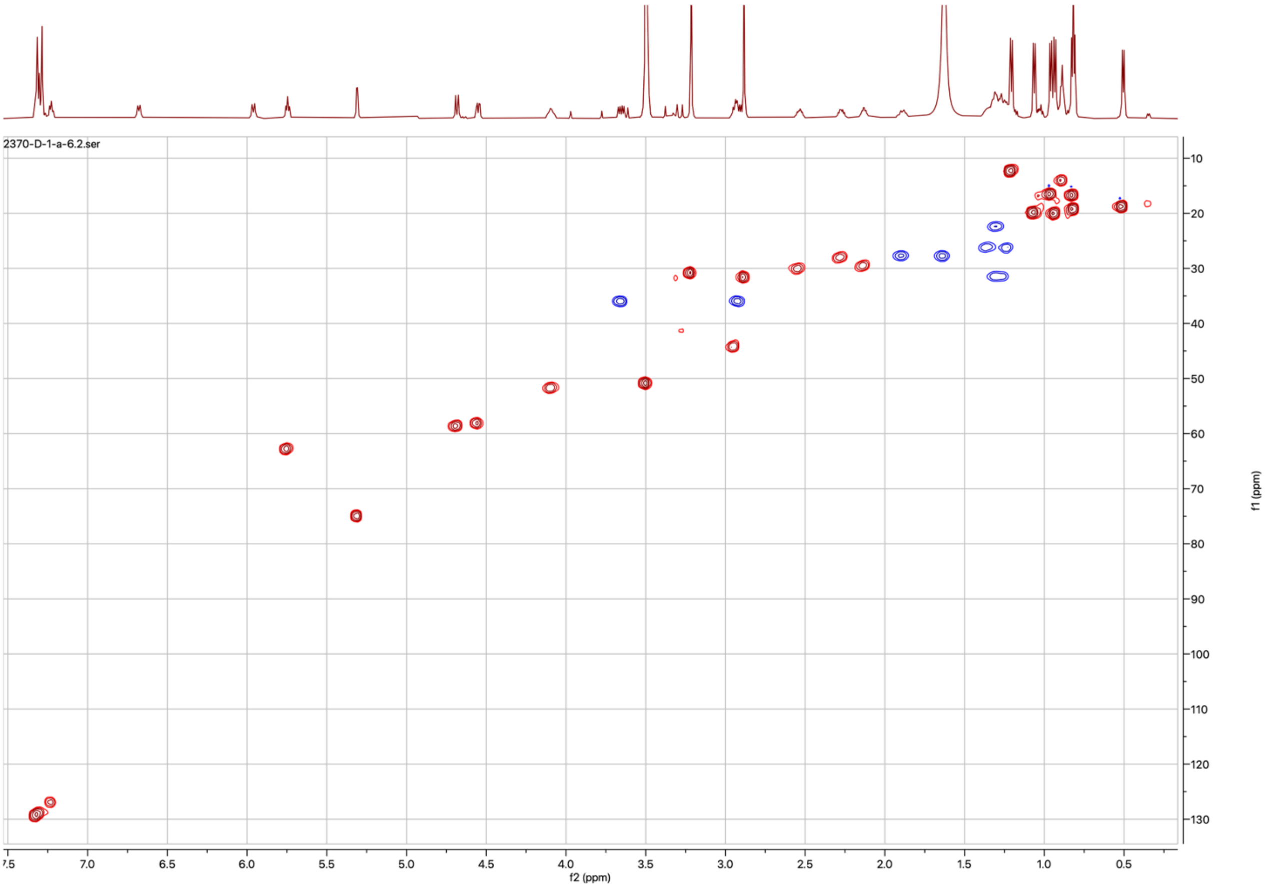

**Figure S28.** Multiplicity-edited HSQC spectrum of boavistamide C (**3**) in $CDCl_3$ (600 MHz).

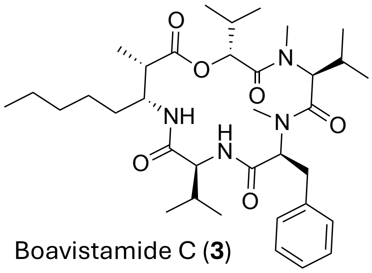

Boavistamide C (**3**)

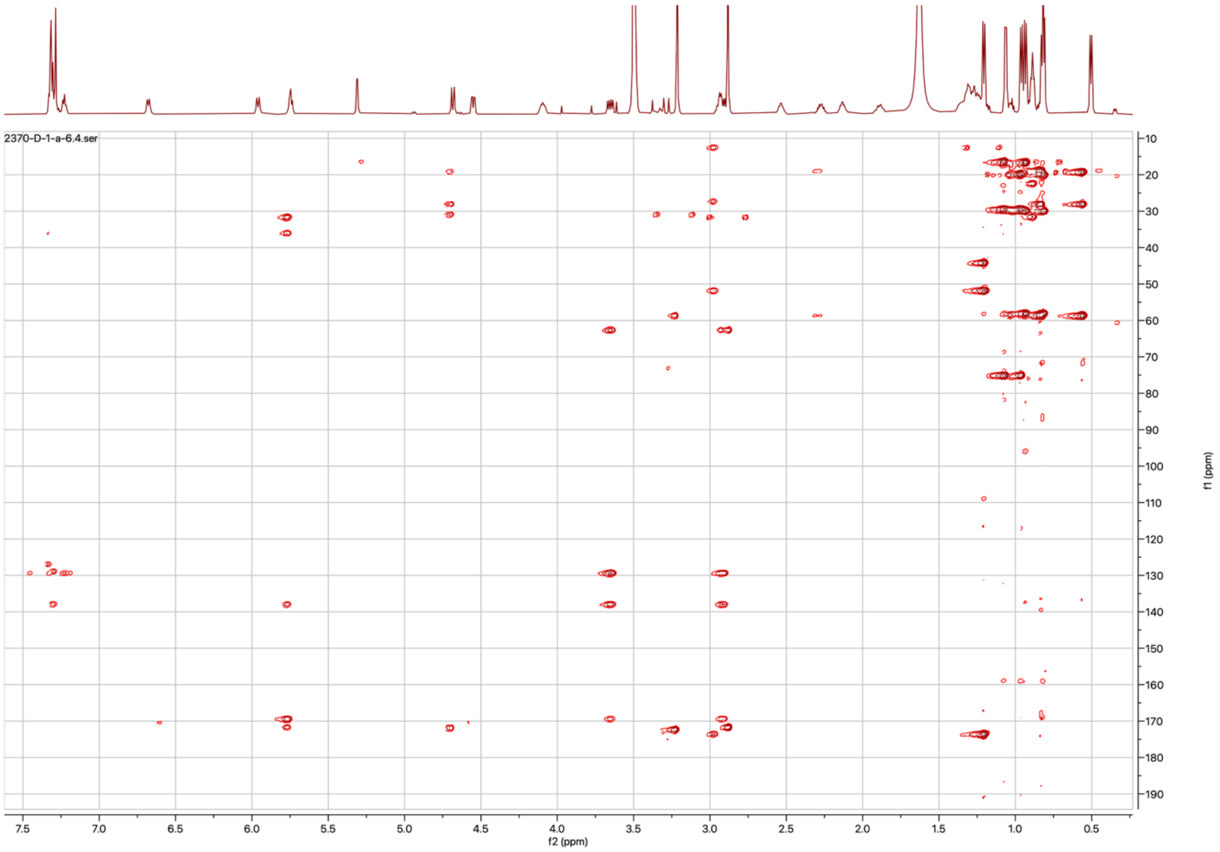

**Figure S29.** HMBC spectrum of boavistamide C (**3**) in $CDCl_3$ (600 MHz).

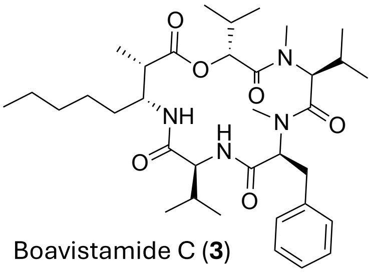

Boavistamide C (**3**)

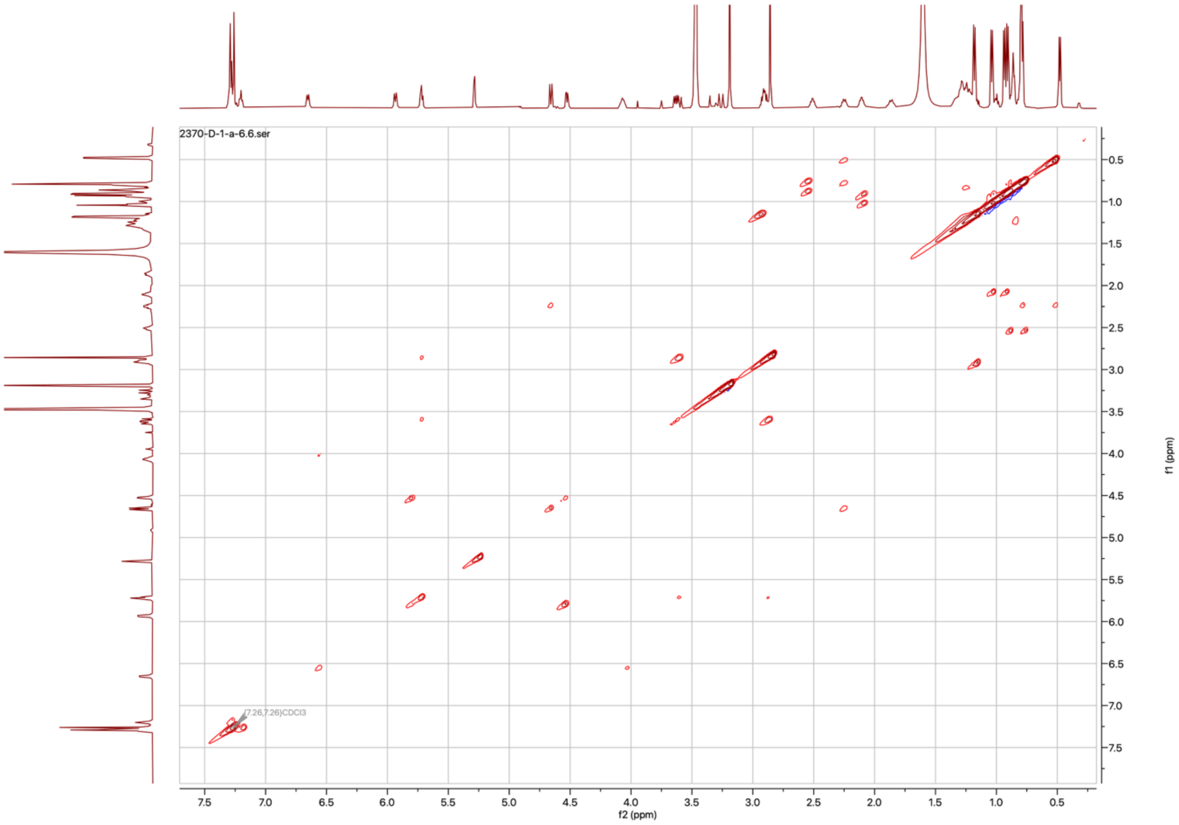

**Figure S30.** $^{1}H$-$^{1}H$ COSY spectrum of boavistamide C (**3**) in $CDCl_3$ (600 MHz).

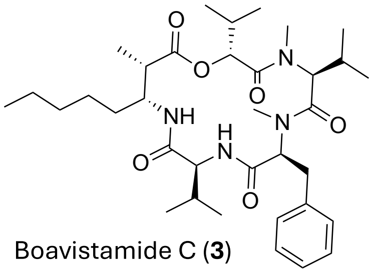

Boavistamide C (**3**)

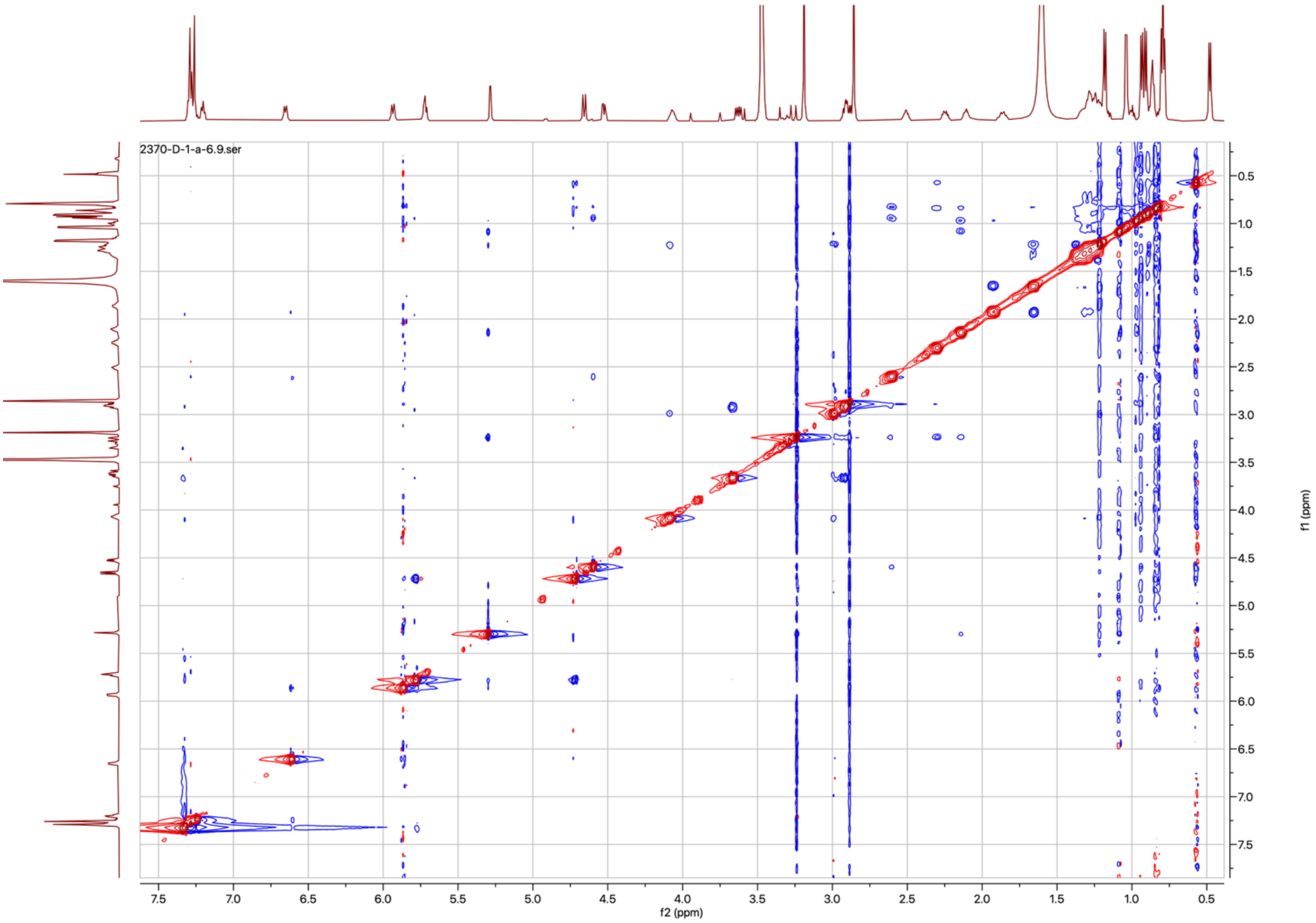

**Figure S31.** ROESY spectrum of boavistamide C (**3**) in $CDCl_3$ (600 MHz).

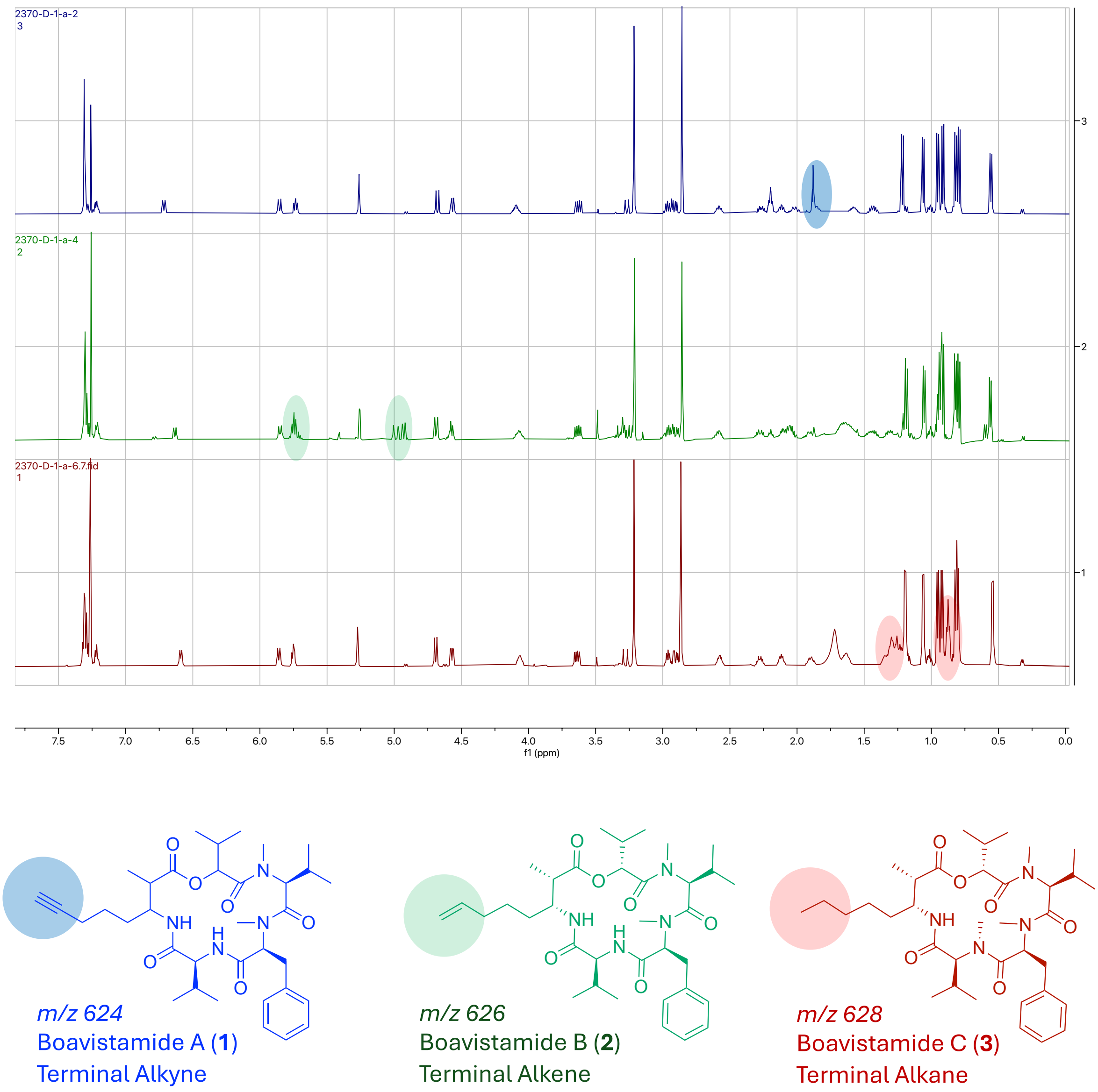

**Figure S32.** Overlay of the $^1$H NMR spectra of boavistamides A (**1**), B (**2**), and C (**3**) in $CDCl_3$, highlighting the structural differences within the AMOYA residue. The progressive reduction of the terminal alkyne (in **1**) to an alkene (in **2**), and then to an alkane (in **3**), is reflected by characteristic changes in the chemical shifts of the olefinic and aliphatic protons.

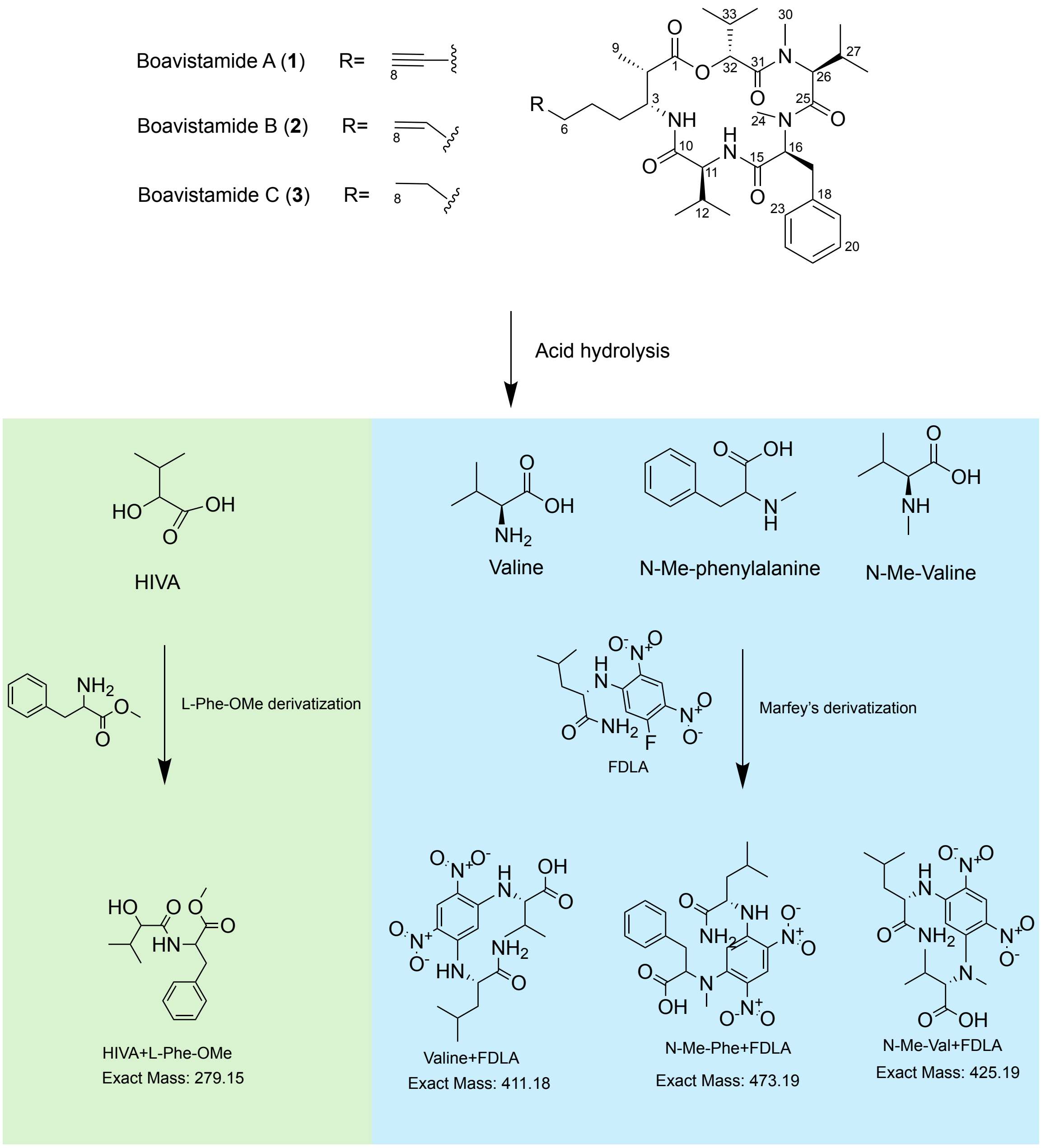

**Figure S33.** Scheme for absolute configuration determination of HIVA, valine, *N*-Me-phenylalanine and *N*-Me-valine residues of boavistamides A–C (**1**–**3**) by hydrolysis and derivatization methods.

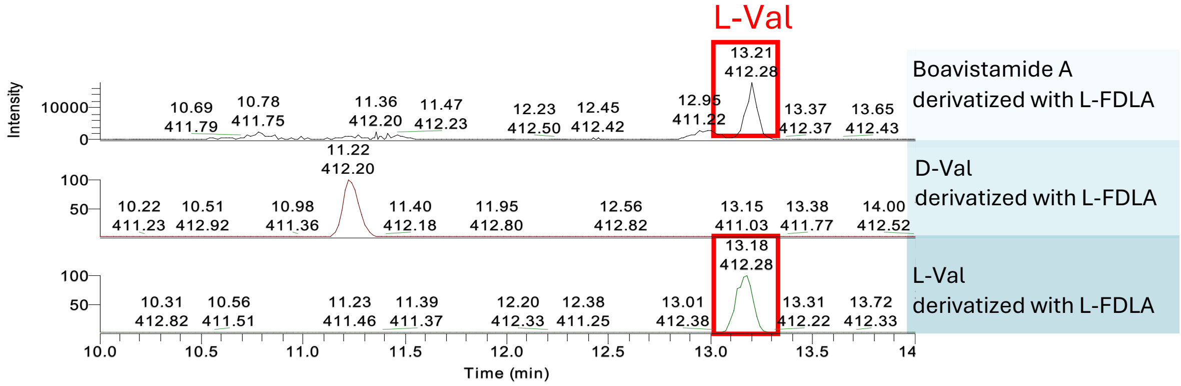

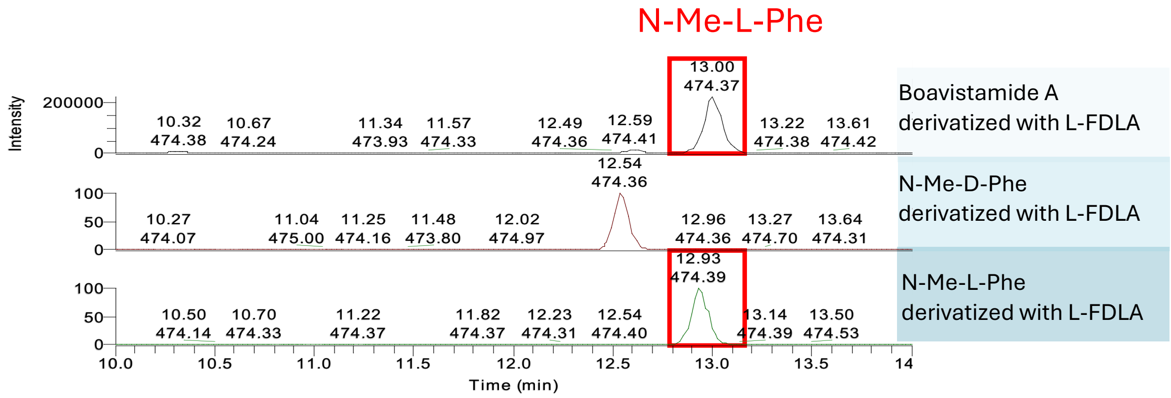

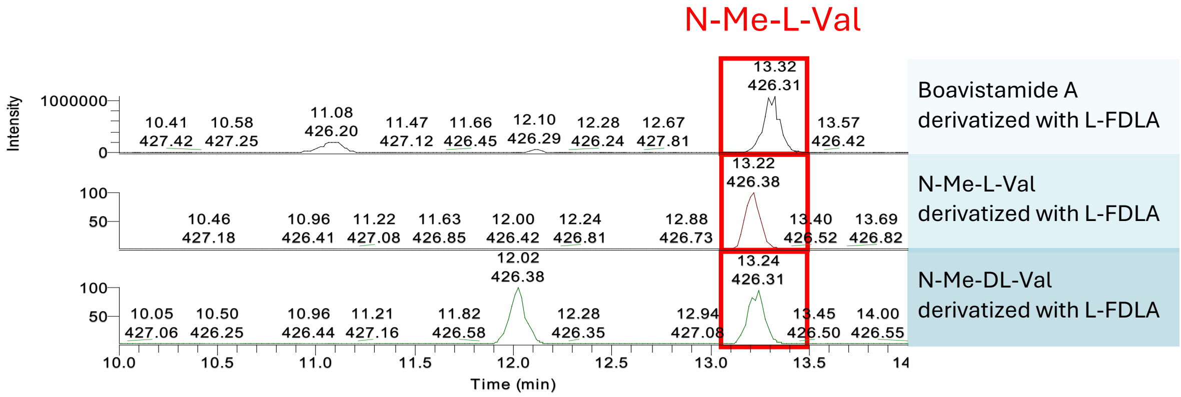

**Figure S34.** Advanced Marfey's analysis of boavistamide A (**1**). LC-MS chromatogram shows the presence of L-Val (m/z 412.28 $[M+H]^+$, RT = 13.21 min), *N*-Me-L-Phe (*m/z* 474.37 $[M+H]^+$, RT = 13.00 min), and *N*-Me-L-Val (*m/z* 426.31 $[M+H]^+$, RT = 13.32 min), identified by comparison with authentic amino acid standards derivatized with L-FDLA. These results support the assignment of L-configurations for all chiral centers in the corresponding residues of boavistamide A (**1**).

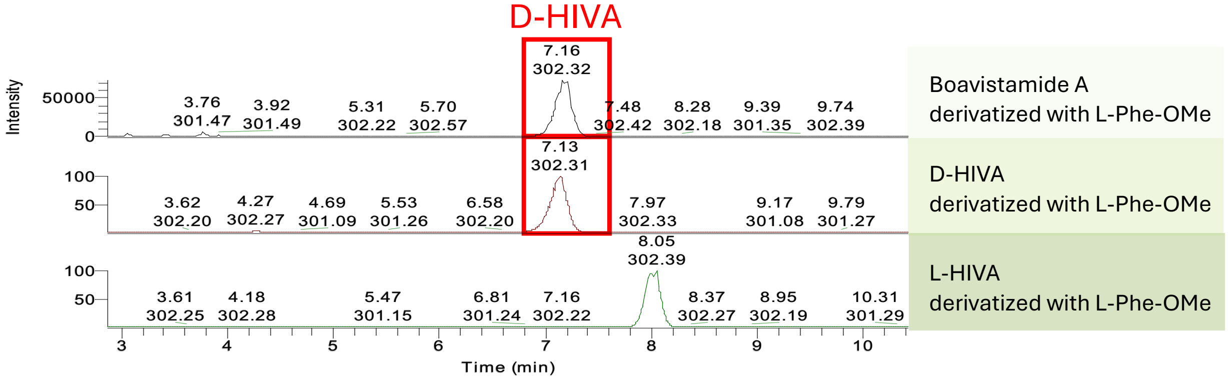

**Figure S35.** L-Phe-OMe derivatization of boavistamide A (**1**). LC-MS analysis reveals a peak at *m/z* 302.32 $[M+Na]^+$ with a retention time of 7.16 min, corresponding to the derivatized D-HIVA, confirming the D-configuration of the hydroxy acid residue.

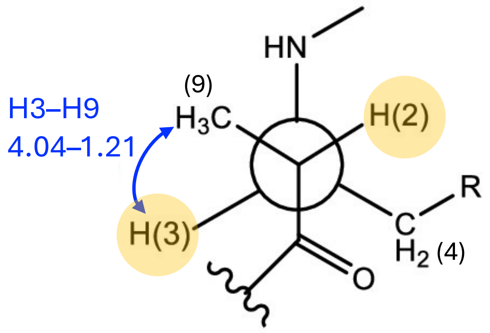

Erythro configuration at C2-C3
Large J H2-H3
(2*S*,3*R*)

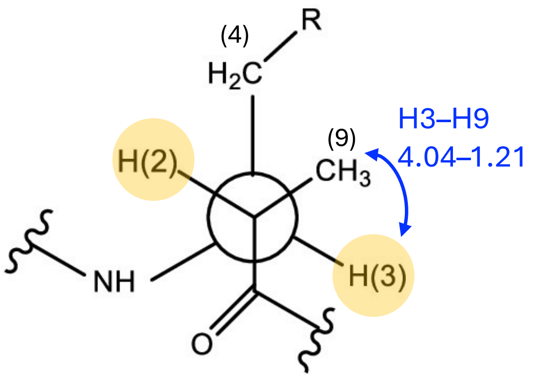

Threo configuration at C2-C3
Large J H2-H3
(2*S*,3*S*)

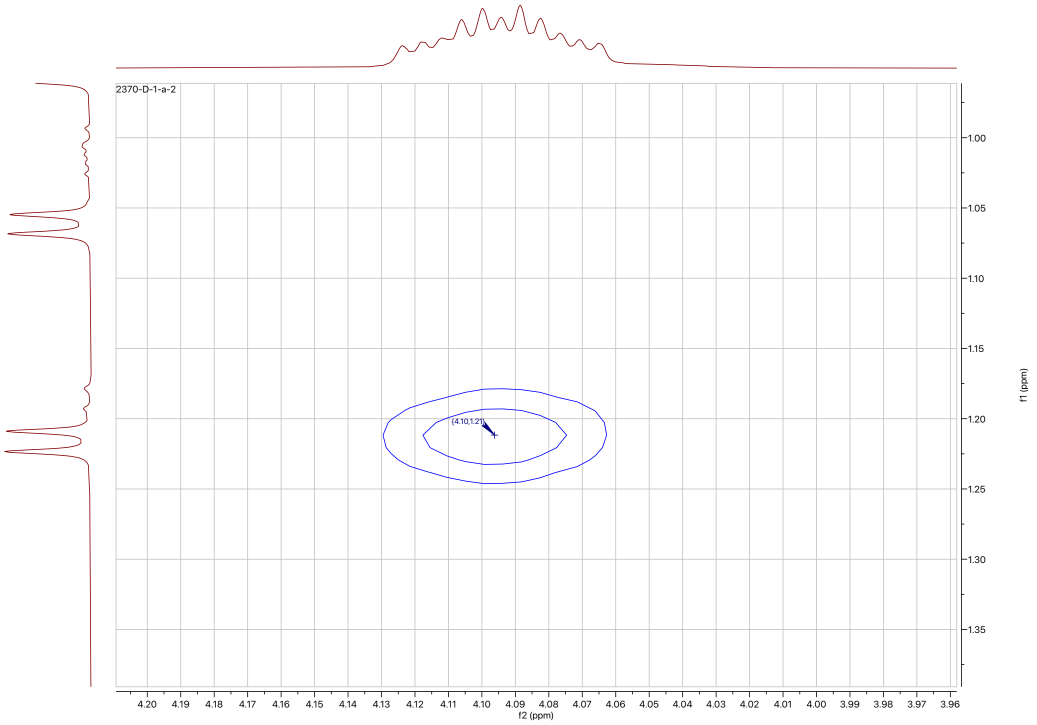

**Figure S36.** ROESY spectrum of boavistamide A (**1**) showing clear cross-peaks between H-3 and $CH_3$–9**,** with corresponding Newman projections illustrating erythro and threo configurations at C-2/C-3.

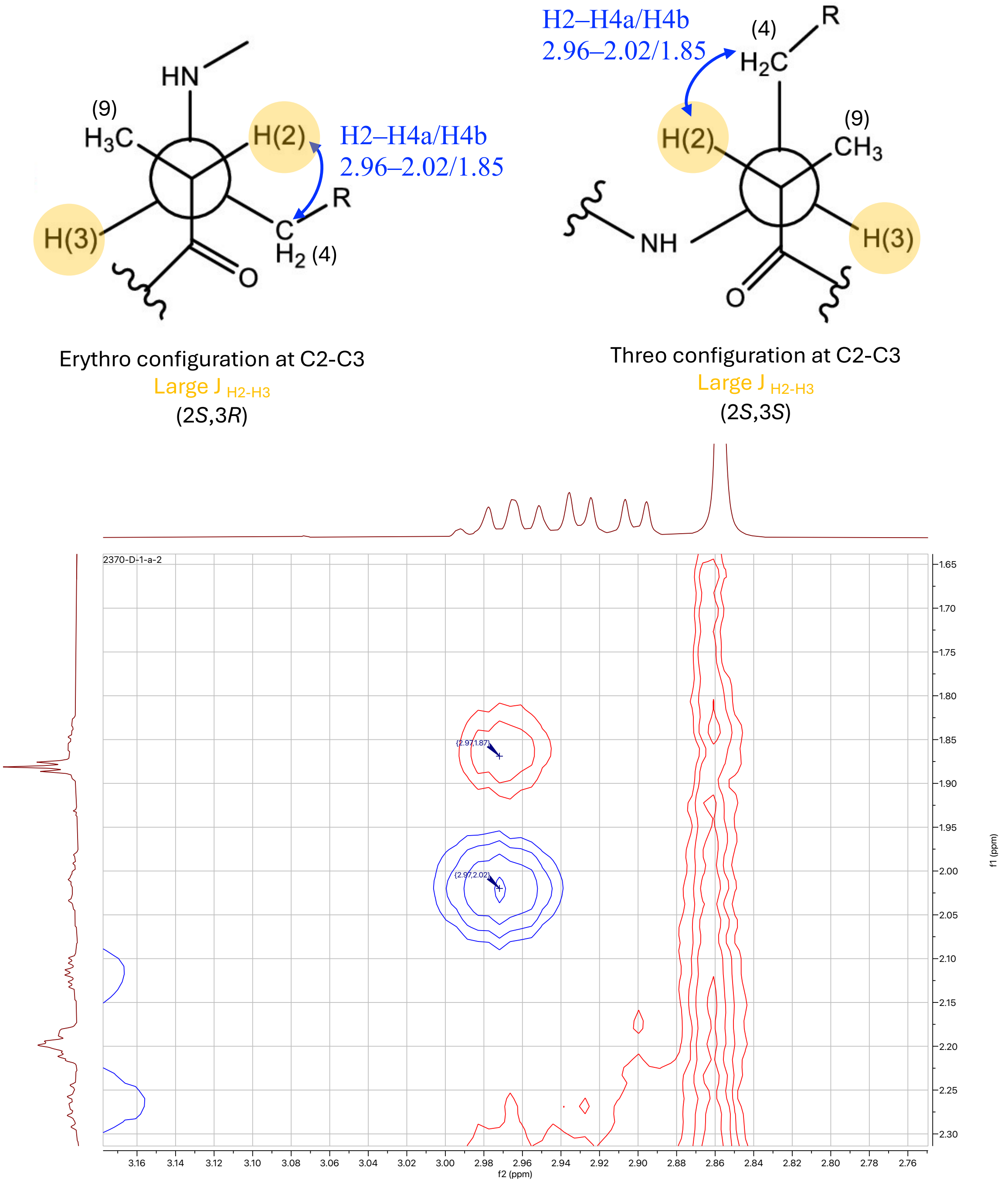

**Figure S37.** ROESY spectrum of boavistamide A (**1**) showing clear cross-peaks between H-2 and H-4**,** with corresponding Newman projections illustrating erythro and threo configurations at C-2/C-3.

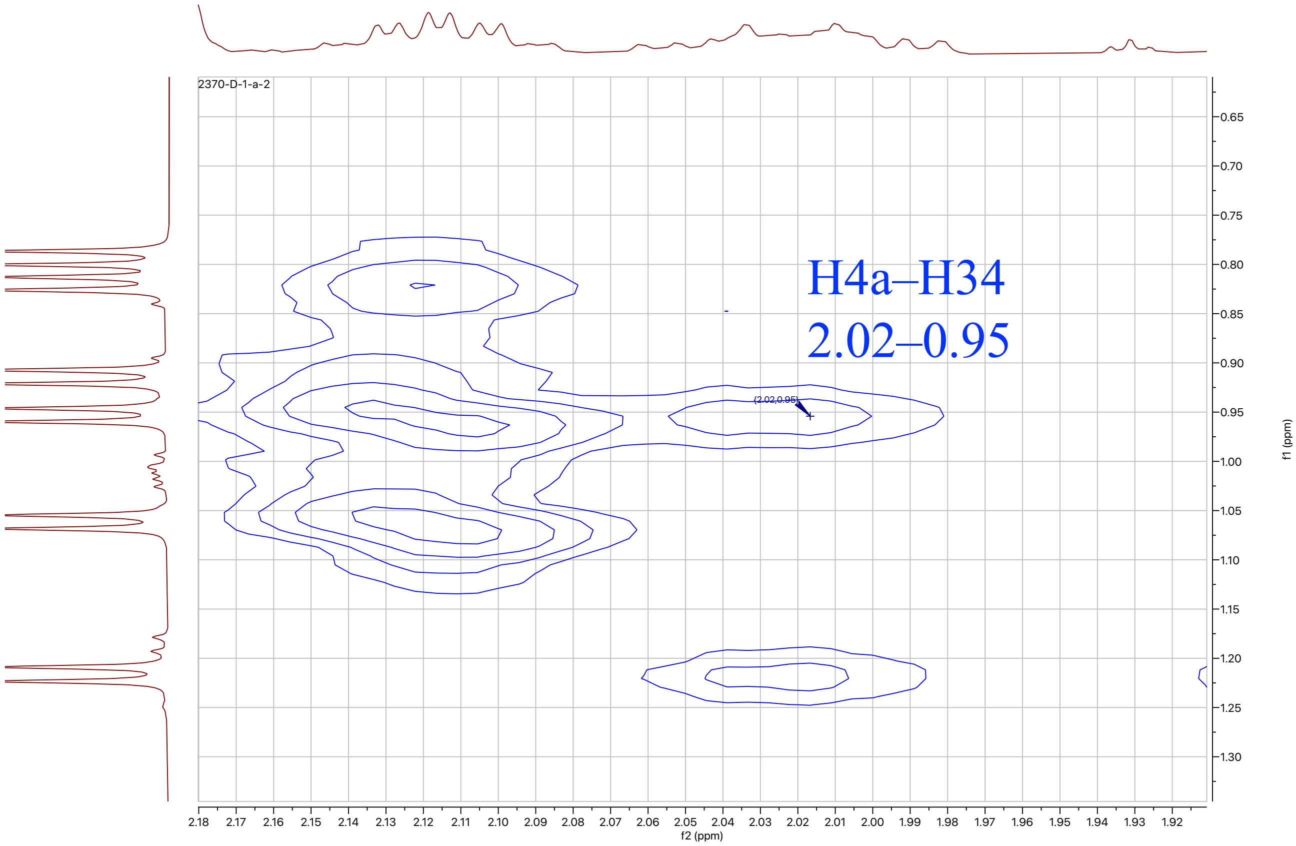

**Figure S38.** ROESY spectrum of boavistamide A (**1**) showing clear cross-peaks between H-4a and H-34**.**

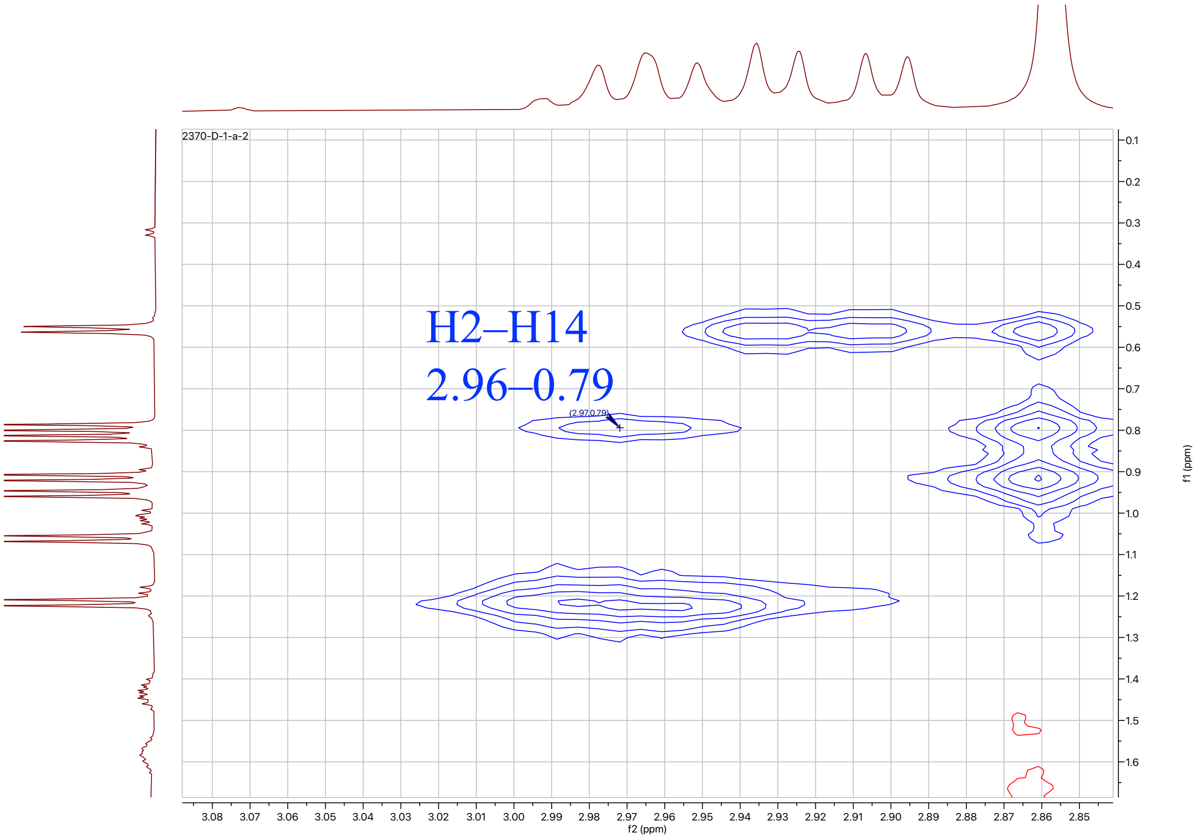

**Figure S39.** ROESY spectrum of boavistamide A (**1**) showing clear cross-peaks between H-2 and H-14**.**

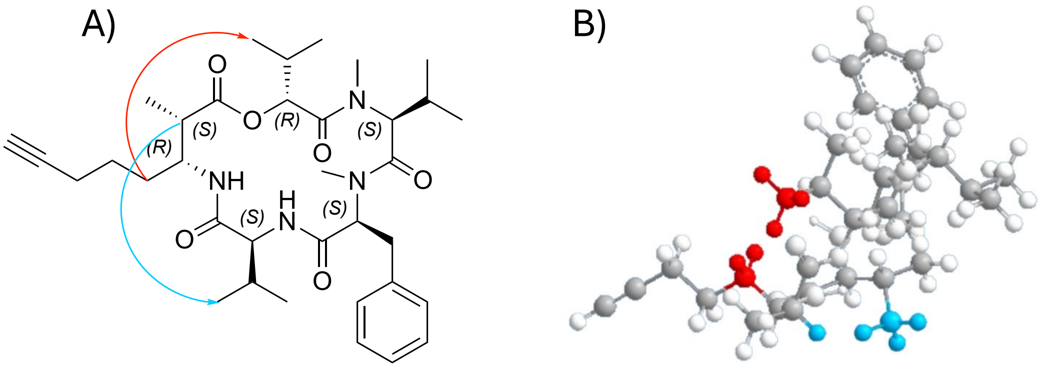

**Figure S40.** Visualization of key inter-residue ROESY correlations in boavistamide A (**1**). (A) 2D structure highlighting ROESY correlations between the AMOYA unit and L-Val (blue) and between the AMOYA unit and D-HIVA (red), shown for the (2*S*,3*R*) AMOYA configuration. (B) 3D model of the (2*S*,3*R*) AMOYA configuration illustrating the corresponding spatial proximity between AMOYA H-2 and the Val methyl group (blue) and between the AMOYA C4 methylene region and the HIVA methyl group (red). The 3D model is consistent with the experimentally observed ROESY correlations.

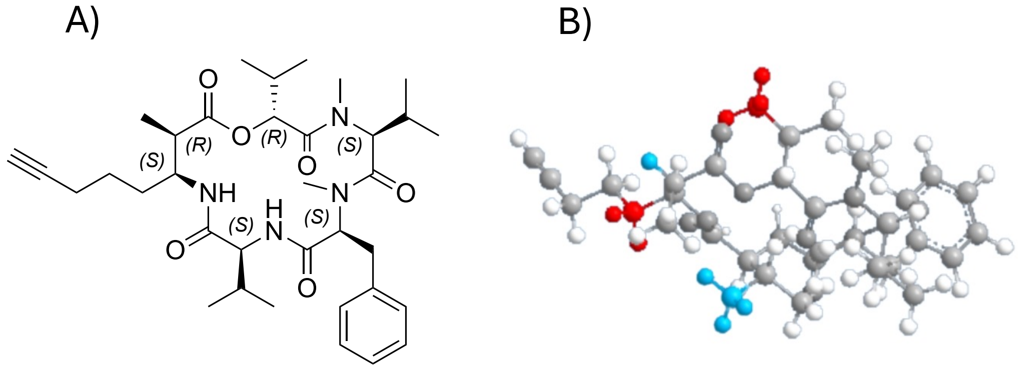

**Figure S41.** Evaluation of the alternative (2*R*,3*S*) AMOYA configuration in boavistamide A (**1**). (A) 2D structure corresponding to the alternative (2*R*,3*S*) AMOYA configuration. (B) 3D model of the (2*R*,3*S*) configuration illustrating that these spatial proximities are not simultaneously observed. See Figure S40 for comparison.

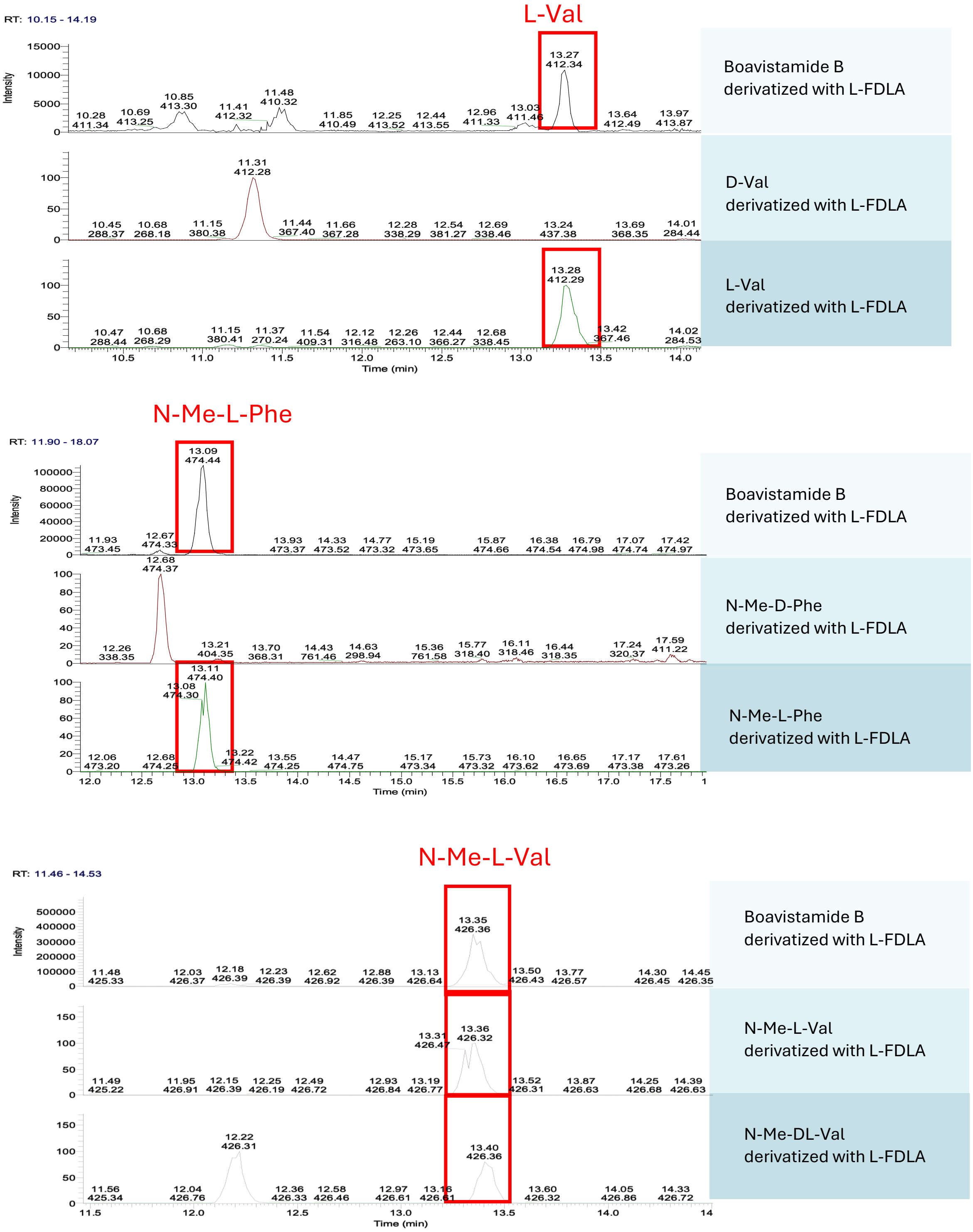

**Figure S42.** Marfey’s analysis of boavistamide B (**2**). LC-MS chromatogram shows the presence of L-Val (m/z 412.34 $[M+H]^+$, RT = 13.27 min), *N*-Me-L-Phe (*m/z* 474.44 $[M+H]^+$, RT = 13.09 min), and *N*-Me-L-Val (*m/z* 426.36 $[M+H]^+$, RT = 13.35 min), identified by comparison with authentic amino acid standards derivatized with L-FDLA. These results support the assignment of L-configurations for all chiral centers in the corresponding residues of boavistamide B (**2**).

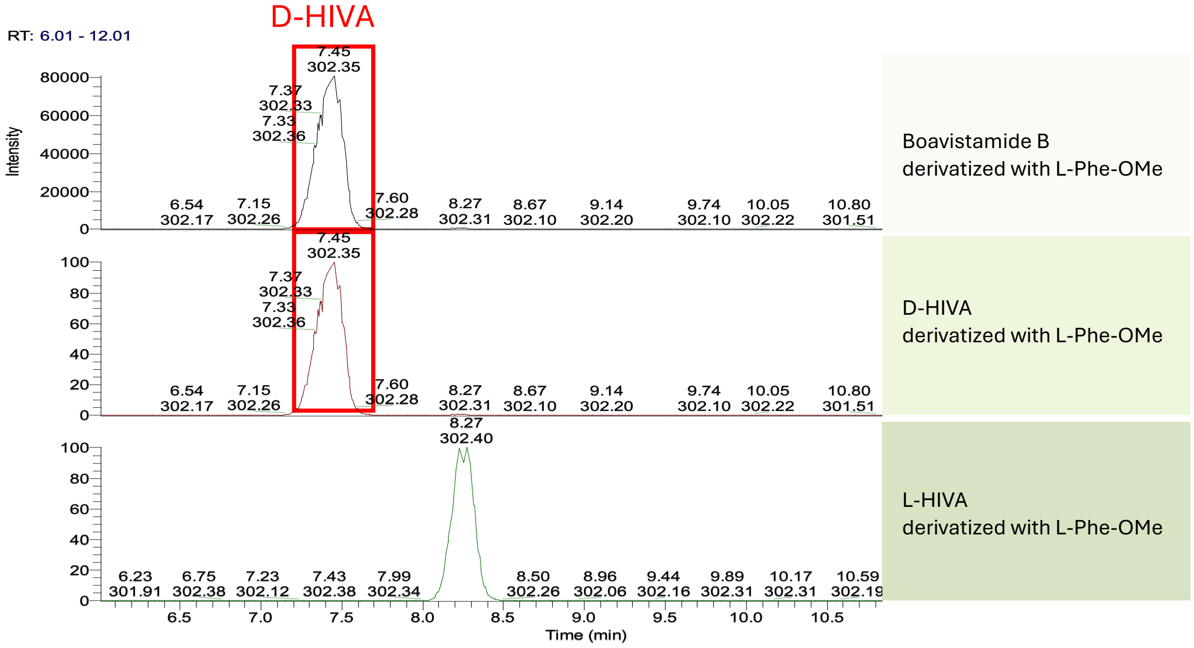

**Figure S43.** L-Phe-OMe derivatization of boavistamide B (**2**). LC-MS analysis revealed a peak at *m/z* 302.35 $[M+Na]^+$ with a retention time of 7.45 min, corresponding to the D-HIVA derivatized, confirming the D-configuration of the hydroxy acid residue.

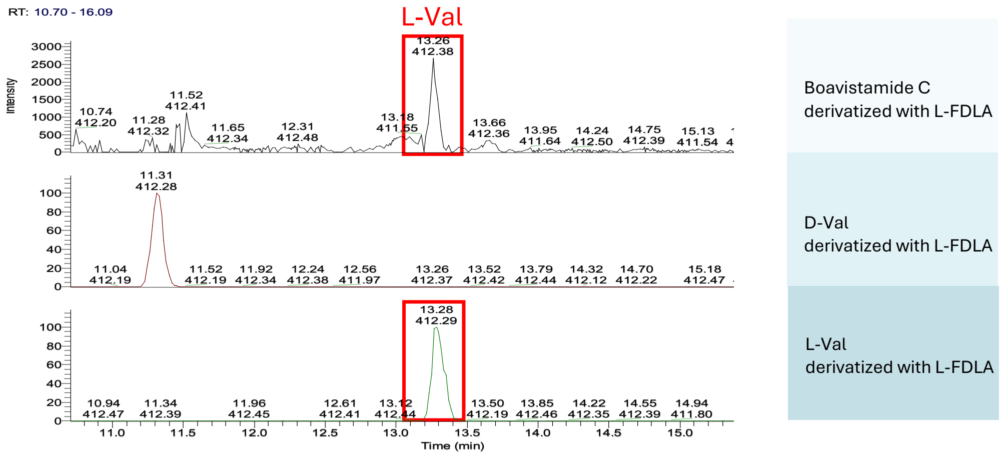

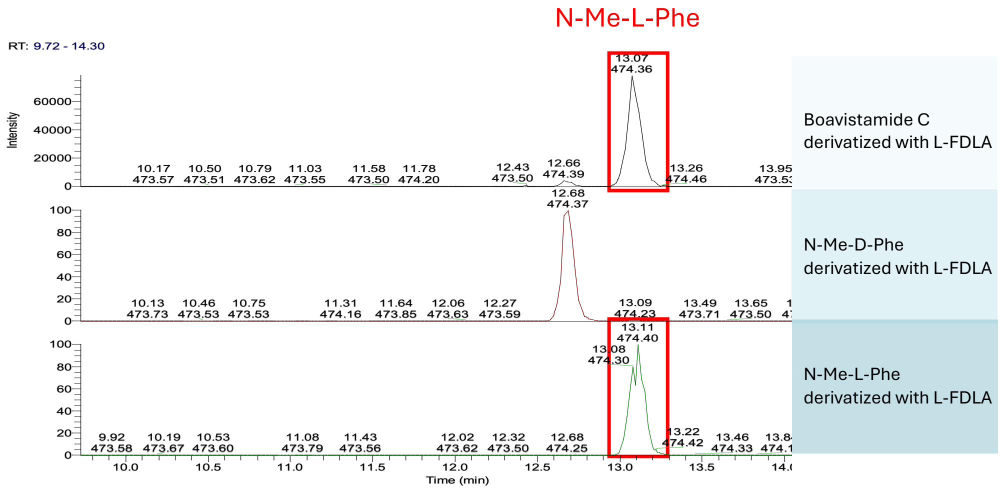

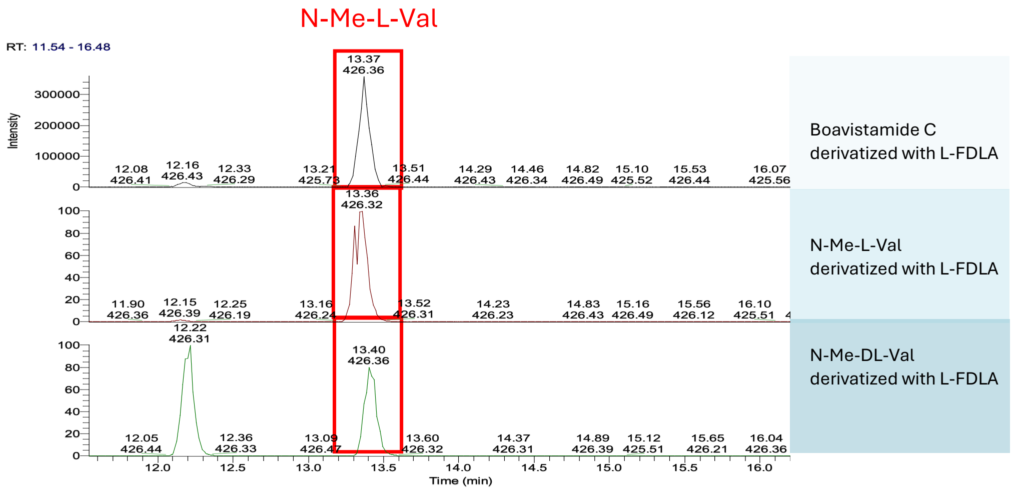

**Figure S44.** Marfey's analysis of boavistamide C (**3**). LC-MS chromatogram shows the presence of L-Val (*m/z* 412.36 $[M+H]^+$, RT = 13.26 min), *N*-Me-L-Phe (*m/z* 474.36 $[M+H]^+$, RT = 13.07 min), and *N*-Me-L-Val (*m/z* 426.36 $[M+H]^+$, RT = 13.36 min), identified by comparison with authentic amino acid standards derivatized with L-FDLA. These results support the assignment of L-configurations for all chiral centers in the corresponding residues of boavistamide C.

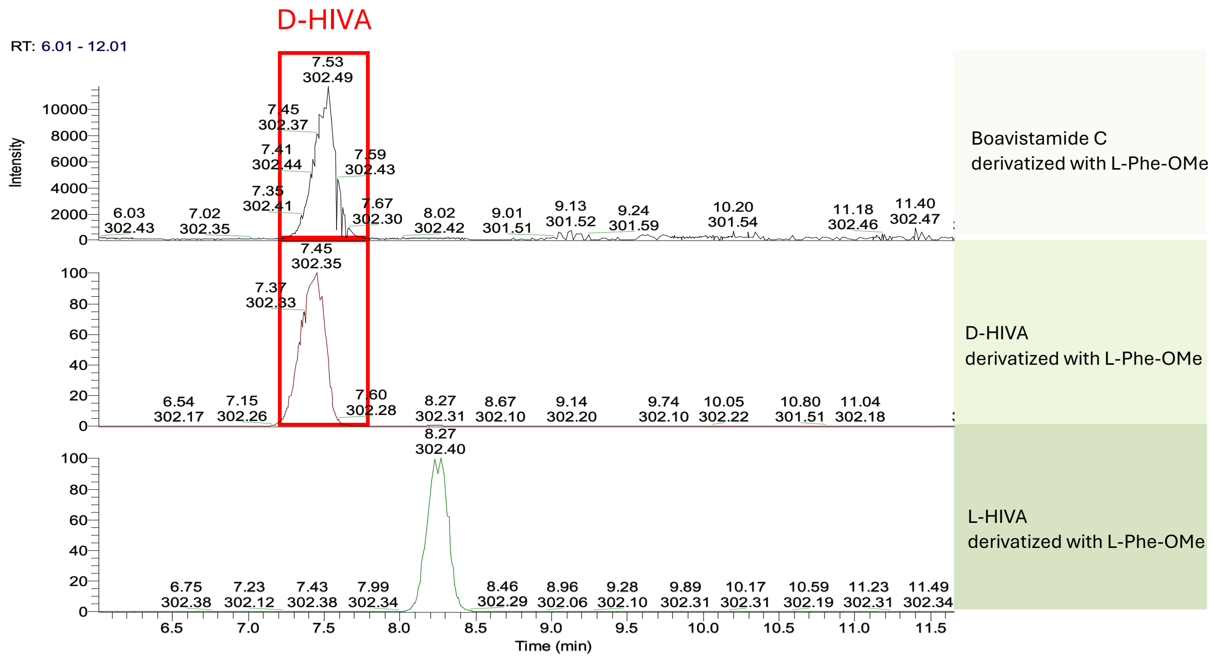

**Figure S45.** L-Phe-OMe derivatization of boavistamide C (**3**). LC-MS analysis reveals a peak at *m/z* 302.49 $[M+Na]^+$ with a retention time of 7.53 min, corresponding to the derivatized D-HIVA, confirming the D-configuration of the hydroxy acid residue.

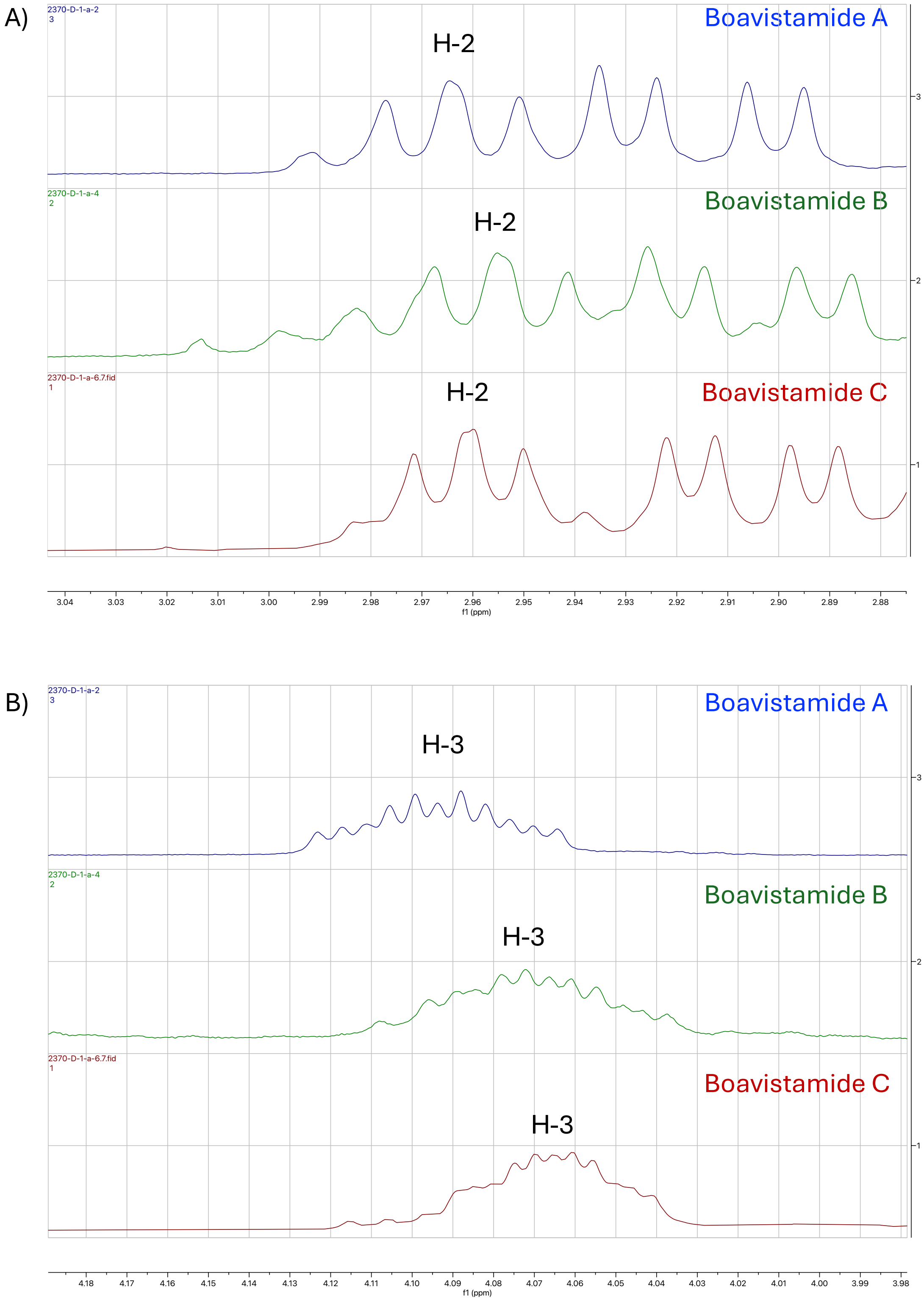

**Figure S46.** $^1$H NMR expansions of the H-2 (A) and H-3 (B) resonance regions of the AMOYA unit in boavistamide A (**1**) and the corresponding protons in boavistamides B (**2**) and C (**3**). Similar multiplet patterns are observed across the three analogs, consistent with conserved stereochemical features of this residue.

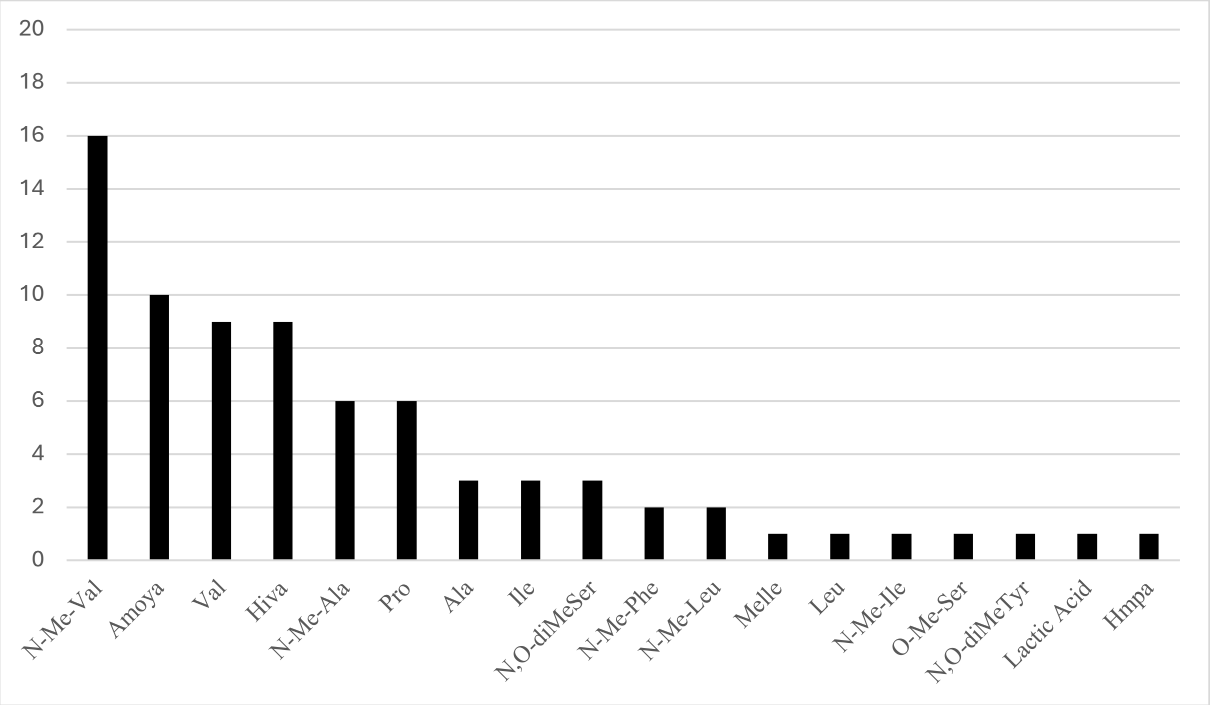

**Figure S47.** Frequency of the residues in nine AMOYA-containing cyclic depsipeptides.

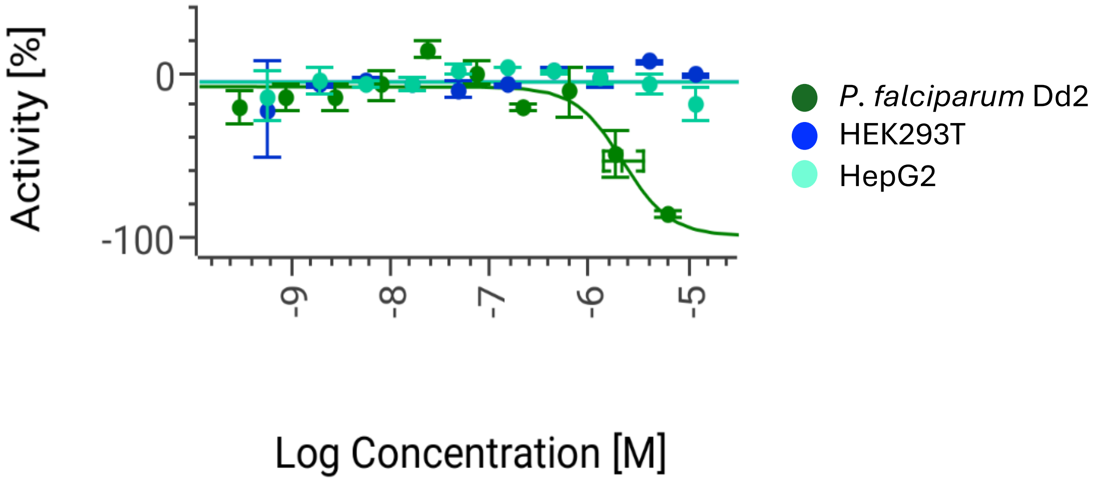

**Figure S48.** Dose response curves of boavistamide A (**1**) against *Plasmodium falciparum* Dd2, HEK293T, and HepG2 cell lines.

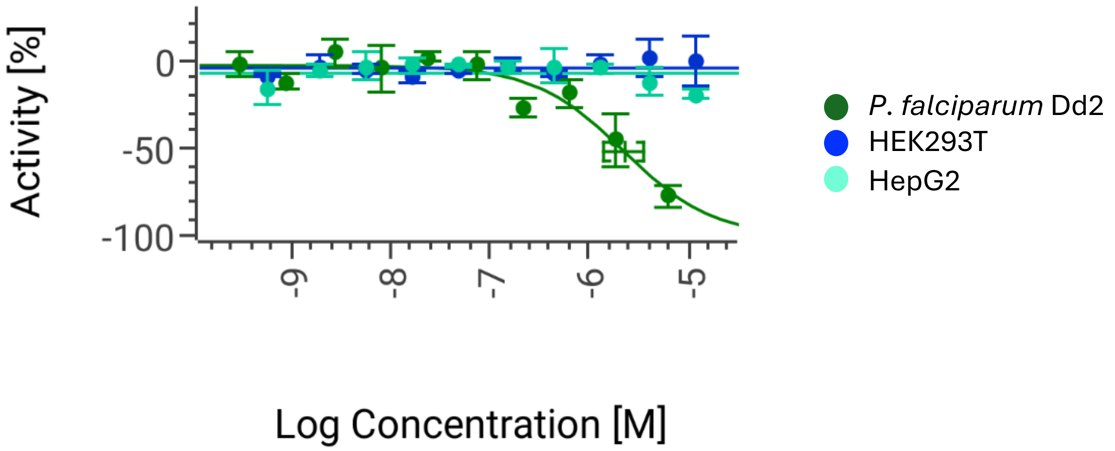

**Figure S49.** Dose response curves of boavistamide B (**2**) against *Plasmodium falciparum* Dd2, HEK293T, and HepG2 cell lines.

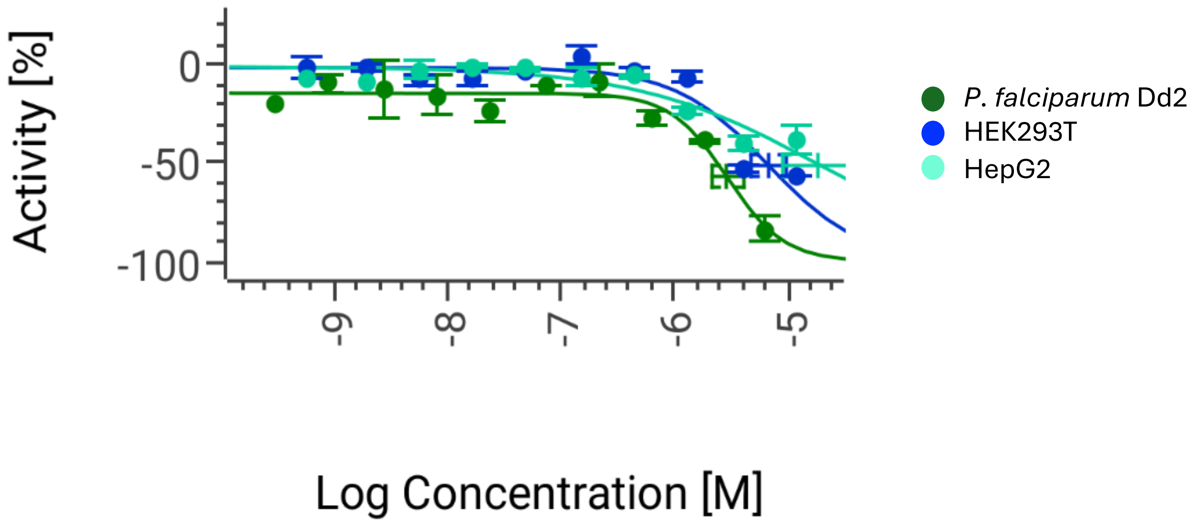

**Figure S50.** Dose response curves of boavistamide C (**3**) against *Plasmodium falciparum* Dd2, HEK293T, and HepG2 cell lines.

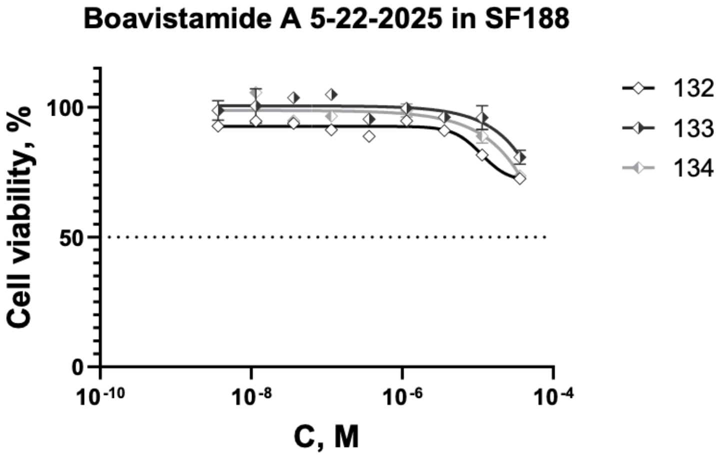

| | 132 | 133 | 134 |
|---|---|---|---|
| IC50 | 1.114e-005 | 0.02177 | 0.004532 |

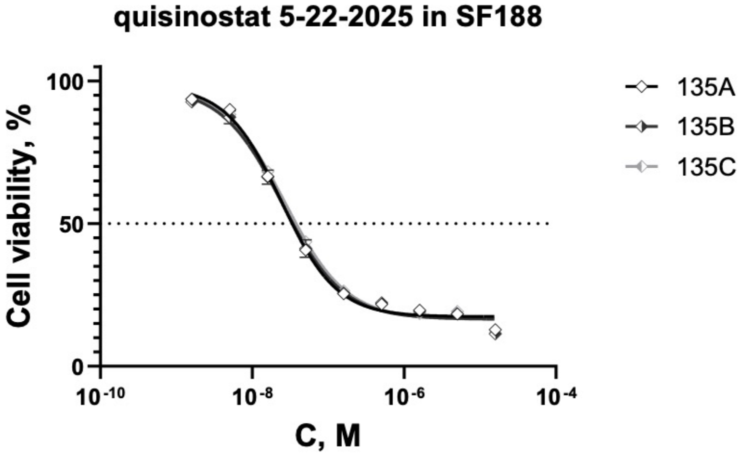

| | 135A | 135B | 135C |
|---|---|---|---|
| IC50 | 2.419e-008 | 2.472e-008 | 2.685e-008 |

**Figure S51.** Concentration response curves of cell viability for glioblastoma cells SF188 treated with boavistamide A (**1**) (top) and positive control quisinostat (bottom)

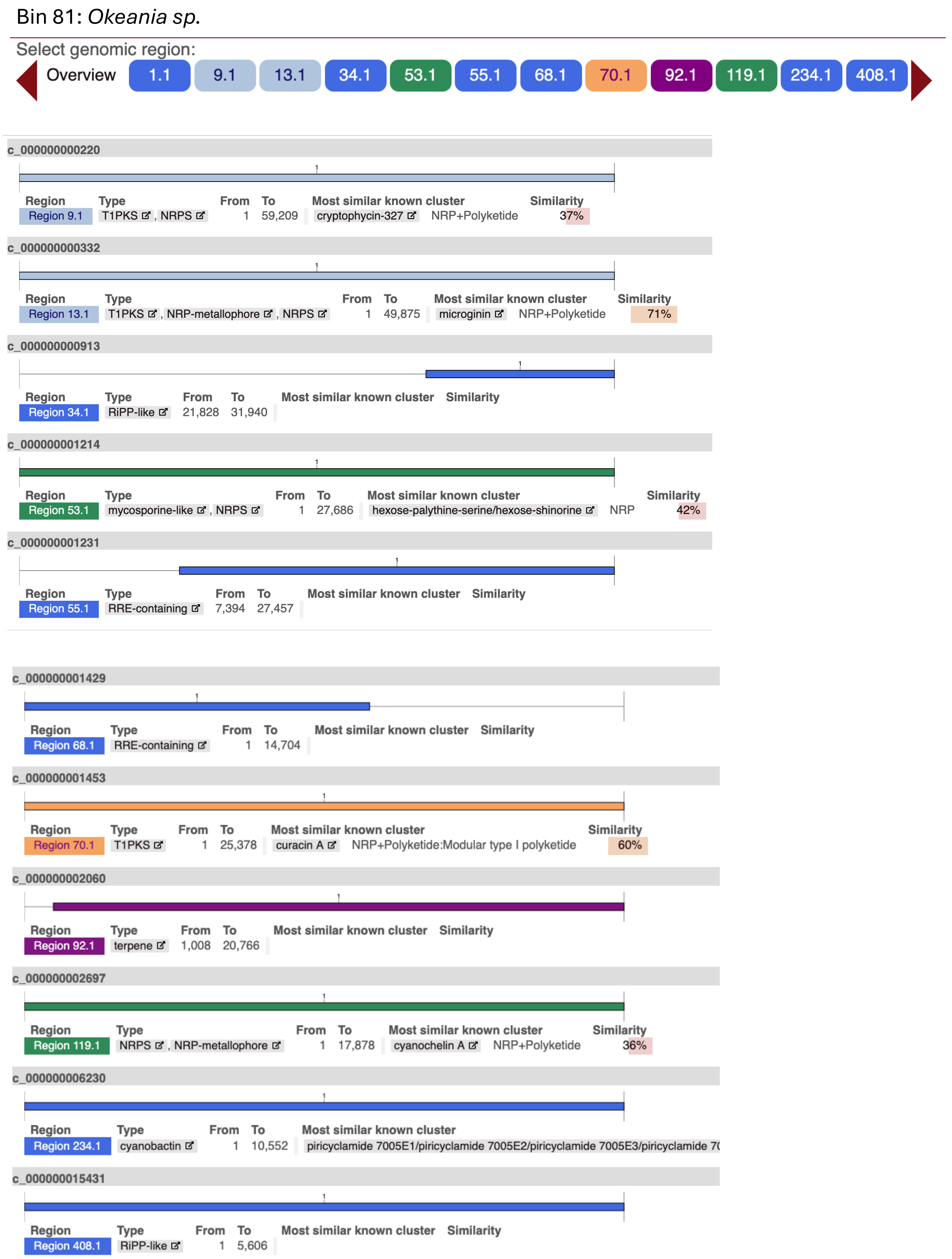

**Figure S52.** AntiSMASH (v7.0) results.

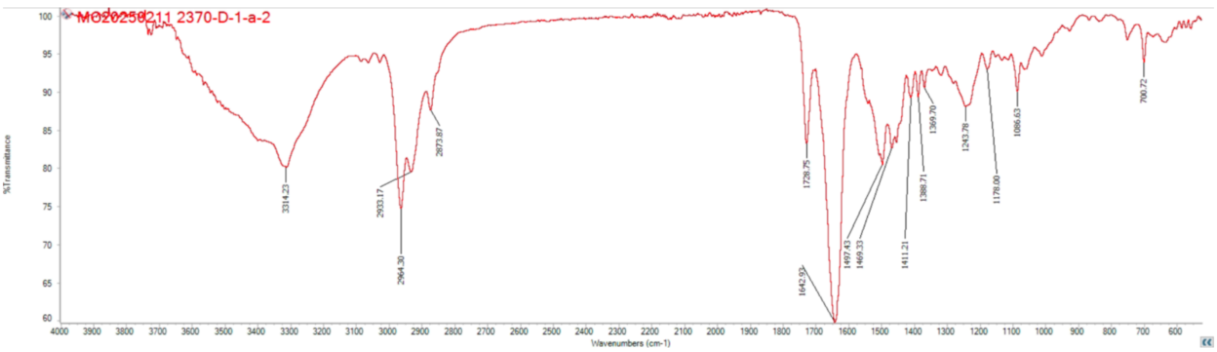

**Figure S53.** IR spectrum of boavistamide A (**1**).

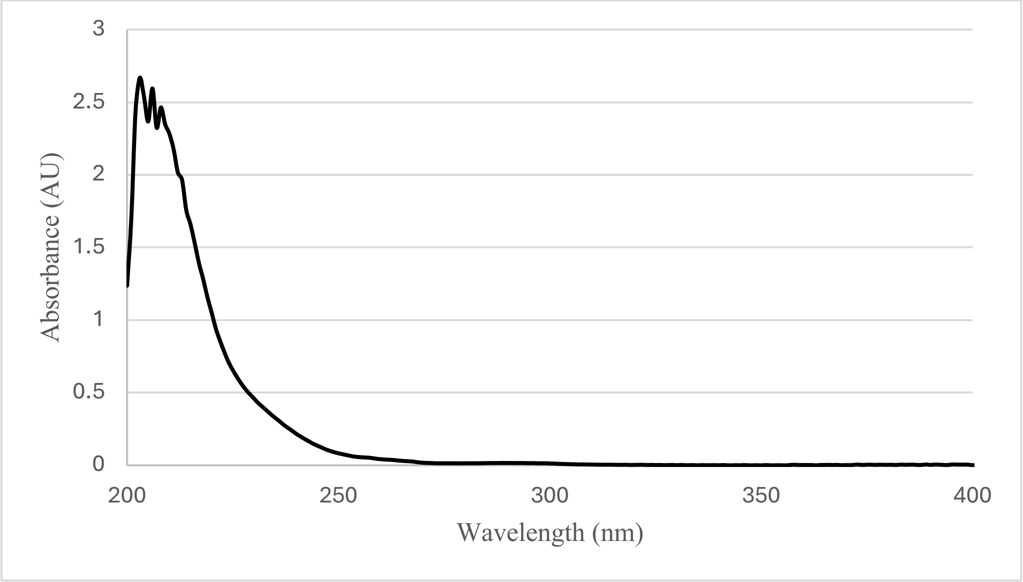

**Figure S54.** UV spectrum of boavistamide A (**1**) in MeOH.

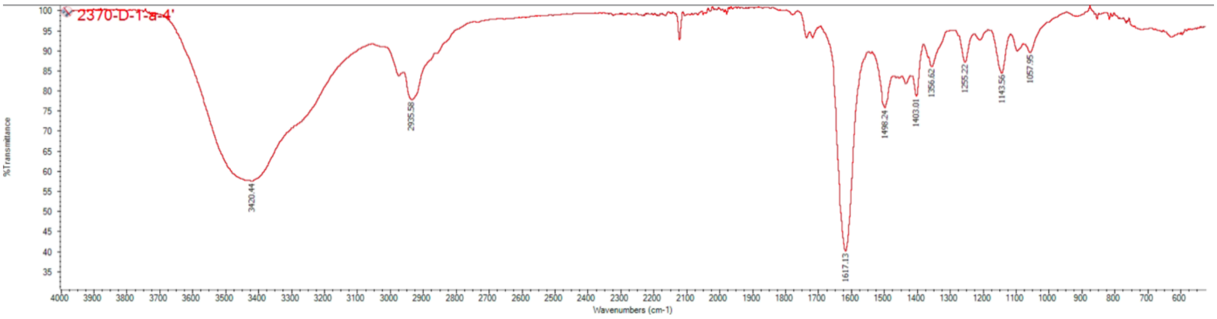

**Figure S55.** IR spectrum boavistamide B (**2**).

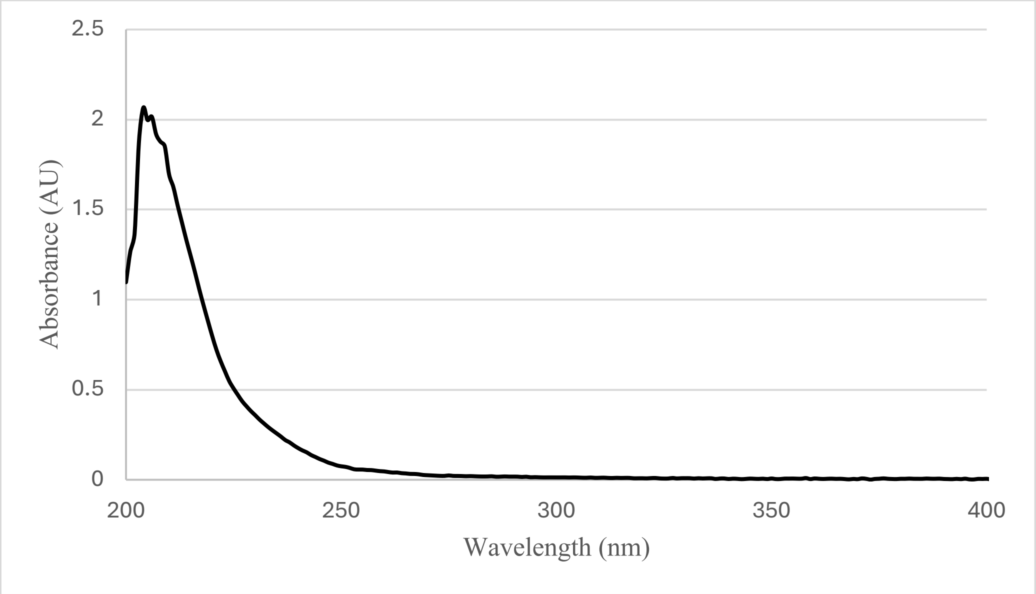

**Figure S56.** UV spectrum of boavistamide B (**2**) in MeOH.

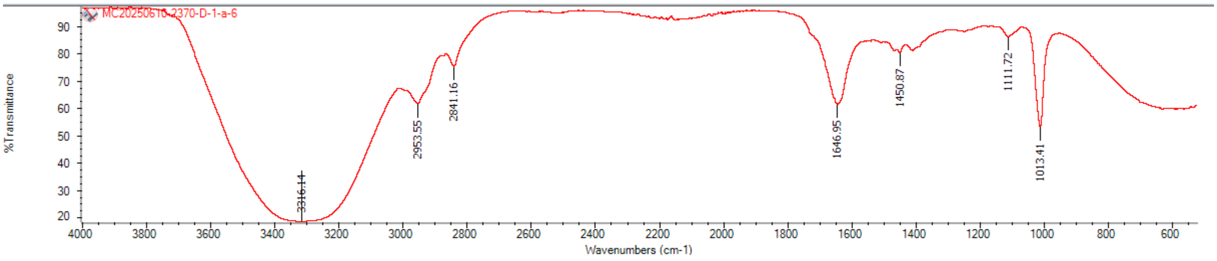

**Figure S57.** IR spectrum boavistamide C (**3**).

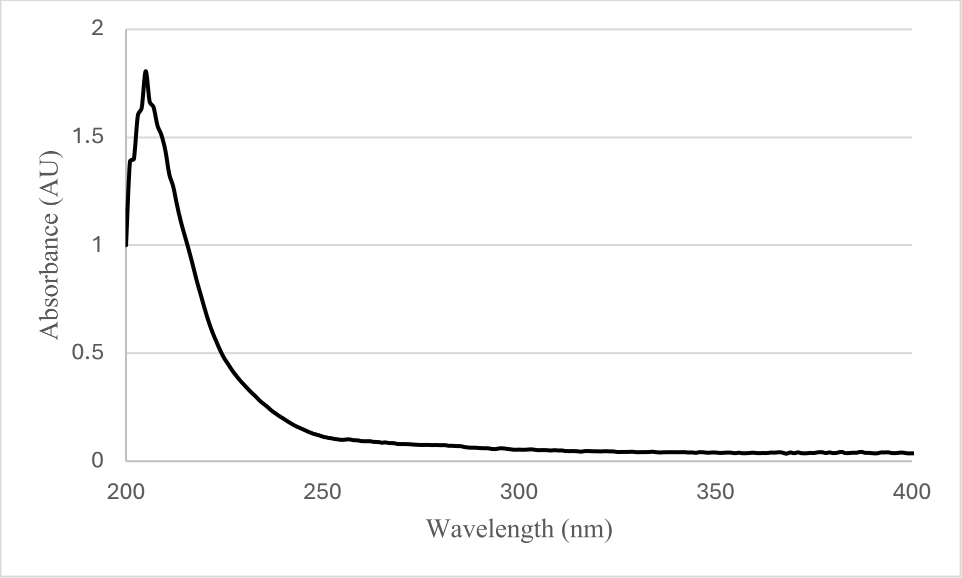

**Figure S58.** UV spectrum of boavistamide C (**3**) in MeOH.

Monti, L.; Liu, L. J.; Varricchio, C.; Lucero, B.; Alle, T.; Yang, W.; Bem-Shalom, I.; Gilson, M.; Brunden, K. R.; Brancale, A.; Caffrey, C. R.; Ballatore, C. Structure-Activity Relationships,

Tolerability and Efficacy of Microtubule-Active 1,2,4-Triazolo[1,5-a]Pyrimidines as Potential Candidates to Treat Human African Trypanosomiasis. *bioRxiv* **2023**, 2023.03.11.532093. https://doi.org/10.1101/2023.03.11.532093

Trager, W.; Jensen, J. B. Human Malaria Parasites in Continuous Culture. *Science* **1976**, 193 (4254), 673–675. https://doi.org/10.1126/science.781840

Hirumi, H.; Hirumi, K. Continuous Cultivation of *Trypanosoma brucei* Blood Stream Forms in a Medium Containing a Low Concentration of Serum Protein without Feeder Cell Layers. *J. Parasitol.* **1989**, 75 (6), 985–989.
